# High-Affinity Protein Binder Design via Flow Matching and In Silico Maturation

**DOI:** 10.64898/2026.01.19.700484

**Authors:** Qilin Yu, Liangyue Guo, Xiayan Qin, Xikun Huang, Baihui Tian, Hongzhun Wang, Yu Liu, Yunzhi Lang, Di Wang, Zhouhanyu Shen, Jie Lin, Mingchen Chen

## Abstract

The de novo design of high-affinity protein binders remains a central challenge in protein engineering and therapeutic discovery. While deep generative models have advanced backbone generation and interface design, achieving picomolar or nanomolar binding affinities typically demands extensive experimental screening or iterative in vitro maturation. A general computational approach for the direct production of high-affinity binders has remained elusive. Here, we introduce an integrated framework that synergizes PPIFlow, a flow-matching-based generative model, with a novel in silico maturation strategy. PPIFlow employs a pairformer architecture to explicitly reason over pair-wise geometric and chemical interactions, modeling protein backbone rigid-body transformations as continuous flows. To bridge the affinity gap, we implement a dedicated in silico affinity maturation stage that combines interface rotamer enrichment with partial flow refinement to optimize energetic packing. This pipeline is further accelerated by AF3Score, a score-only adaptation of AlphaFold3 that enables high-fidelity and computationally efficient candidate prioritization. Across a diverse set of therapeutic targets, this synergistic approach consistently produces picomolar and nanomolar affinity binders without experimental affinity maturation. Notably, the framework proves highly effective for the de novo design of single-domain antibodies (VHHs), producing sub-nanomolar binders across multiple targets. These results establish that coupling robust backbone generation with focused in silico maturation renders the purely computational design of high-affinity binders feasible.

## 1 Introduction

Protein–protein interactions (PPIs) underpin nearly all cellular processes and represent a primary class of therapeutic targets. The ability to design proteins that bind specific epitopes with high affinity and precision would revolutionize applications in pathway modulation, diagnostics, and biologic drug development [1]. Traditional binder discovery, relying on immunization [2], library screening [3, 4], and directed evolution [5, 6], remains resource-intensive and often offers limited control over the targeted epitope or the resulting molecular geometry.

Deep learning has recently catalyzed a paradigm shift in *de novo* protein design. Generative approaches utilizing structure hallucination [7, 8], inverse folding [1, 9], and diffusion models [10, 11] have enabled the creation of novel folds and functional binders. Notable frameworks such as RFdiffusion and BindCraft have significantly improved backbone sampling and diversity, and design of binders [10, 12, 13, 14]. However, a critical bottleneck remains: most computationally designed binders exhibit only modest affinities, typically requiring extensive experimental screening or iterative *in vitro* maturation to achieve therapeutic relevance.

Achieving high-affinity binding is fundamentally more challenging than generating plausible interfaces. High-affinity PPIs require exacting geometric complementarity, optimized side-chain packing, and cooperative interactions that are difficult for backbone-centric generative models to capture [14]. Antibodies, which naturally achieve strong and specific protein–protein interactions, therefore represent a particularly stringent benchmark for *de novo* design. While recent advances in *de novo* antibody design, such as RFantibody and Germinal, have expanded the accessible space for complementarity-determining regions (CDRs) [11, 15], generation of high-affinity binders and single-domain antibodies (VHHs) with high success rate and without the need for library-based directed evolution remains an elusive goal.

Flow matching has recently emerged as a powerful alternative to diffusion, offering superior training stability and efficient sampling by learning continuous mappings between simple distributions and complex structures [16, 17, 18, 19]. Here, we introduce **PPIFlow**, a flow-matching–based framework and integrated design workflow for the *de novo* generation of high-affinity binders targeting precise epitopes (Fig. 1a). PPIFlow integrates a pairformer module to explicitly model interface contacts and utilizes a multi-stage training scheme for robust generalization across diverse protein scaffolds. To achieve *in silico* affinity maturation, we introduce three key innovations: (i) **interface rotamer enrichment** to identify energetically optimal residues; (ii) **partial flow refinement** to optimize interface packing by perturbing and regenerating existing designs; and (iii) **AF3Score**, an efficient score-only adaption of AlphaFold3 (Fig. 1c) for high-confidence complex prioritization[20].

**Figure 1:**
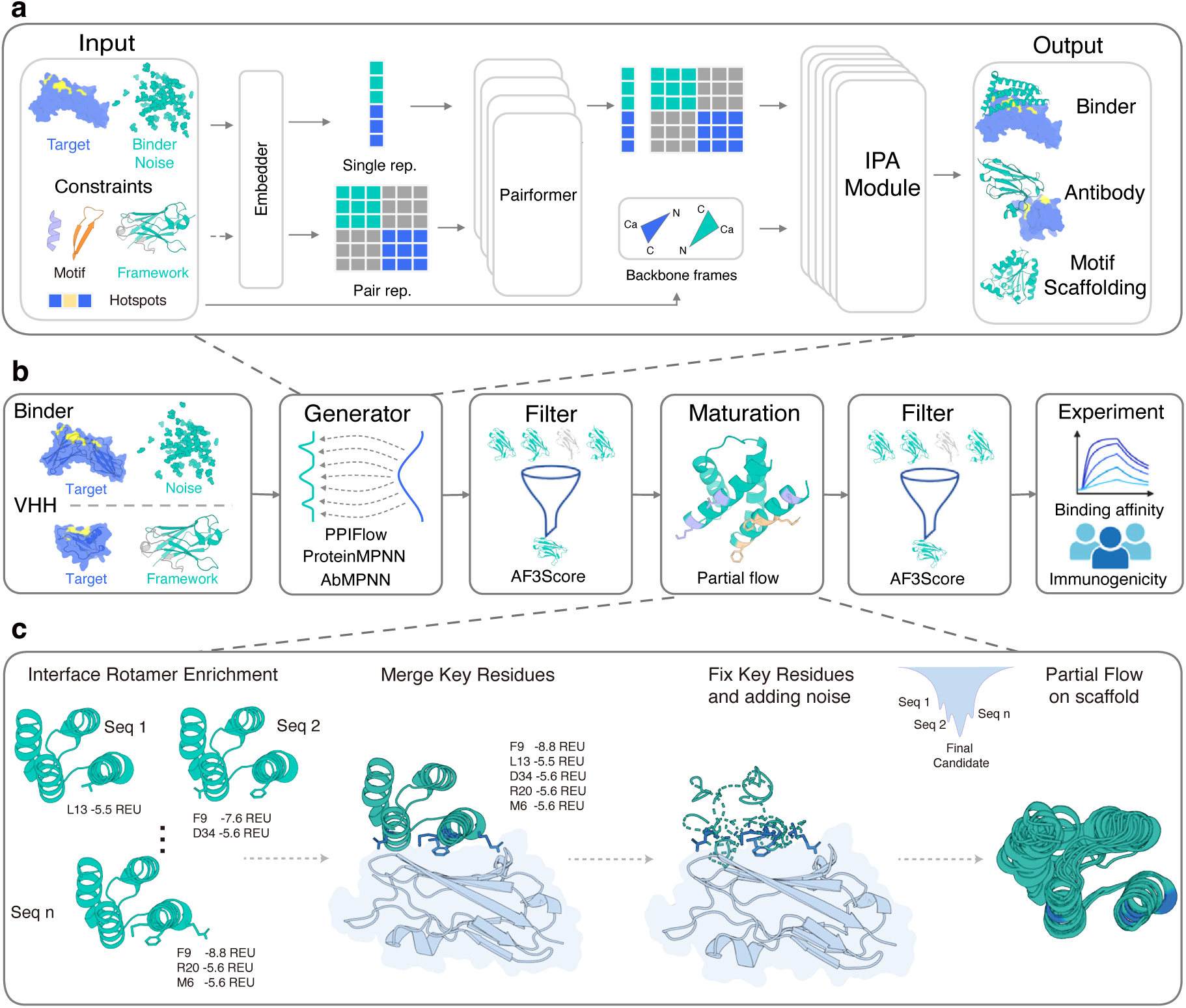
PPIFlow model architecture and high-affinity design workflow. a, Overview of the PPIFlow model framework. b, End-to-end design pipeline. PPIFlow-generated backbones undergo sequence design via ProteinMPNN (monomers, motifs, binders) or AbMPNN (VHHs, scFvs). High-confidence candidates are identified via AF3Score filtering and prioritized for *in silico* maturation and template-free AlphaFold3 validation. c, *In silico* maturation strategy: (1) interface rotamer enrichment to identify energetically favorable residues; (2) selection and merging of key interface residues; (3) fixation of critical residues with concomitant noise addition to neighboring backbone regions; and (4) partial flow-based refinement of unconstrained regions to optimize interfacial packing.

Using this integrated platform, we demonstrate unprecedented performance, successfully generating pM-affinity binders for six out of seven diverse therapeutic targets and pM to nM-affinity VHHs for seven out of eight targets. These results establish PPIFlow as a reliable platform for the *de novo* design of protein and antibody therapeutics with epitope precision.

## 2 Results

### 2.1 A flow matching based protein generation model

PPIFlow models protein binder design as a continuous rigid-body flow. We represent protein backbones as collections of rigid residue frames in SE(3) [21] (Fig. 1a). Unlike diffusion-based approaches [10], PPIFlow employs flow matching to directly learn the translational and rotational velocity fields that transport random distributions to physically realistic structures [16, 18]. This formulation provides a geometrically consistent parameterization of rigid-body dynamics, enabling stable training and precise conditioning on interface constraints.

The pairformer module jointly updates residue-level and pairwise embeddings to explicitly capture structured geometric relationships across binding interfaces [22]. These representations are decoded into frame updates via Invariant Point Attention (IPA) [21], balancing generation accuracy with parameter efficiency (Fig. 2c).

**Figure 2:**
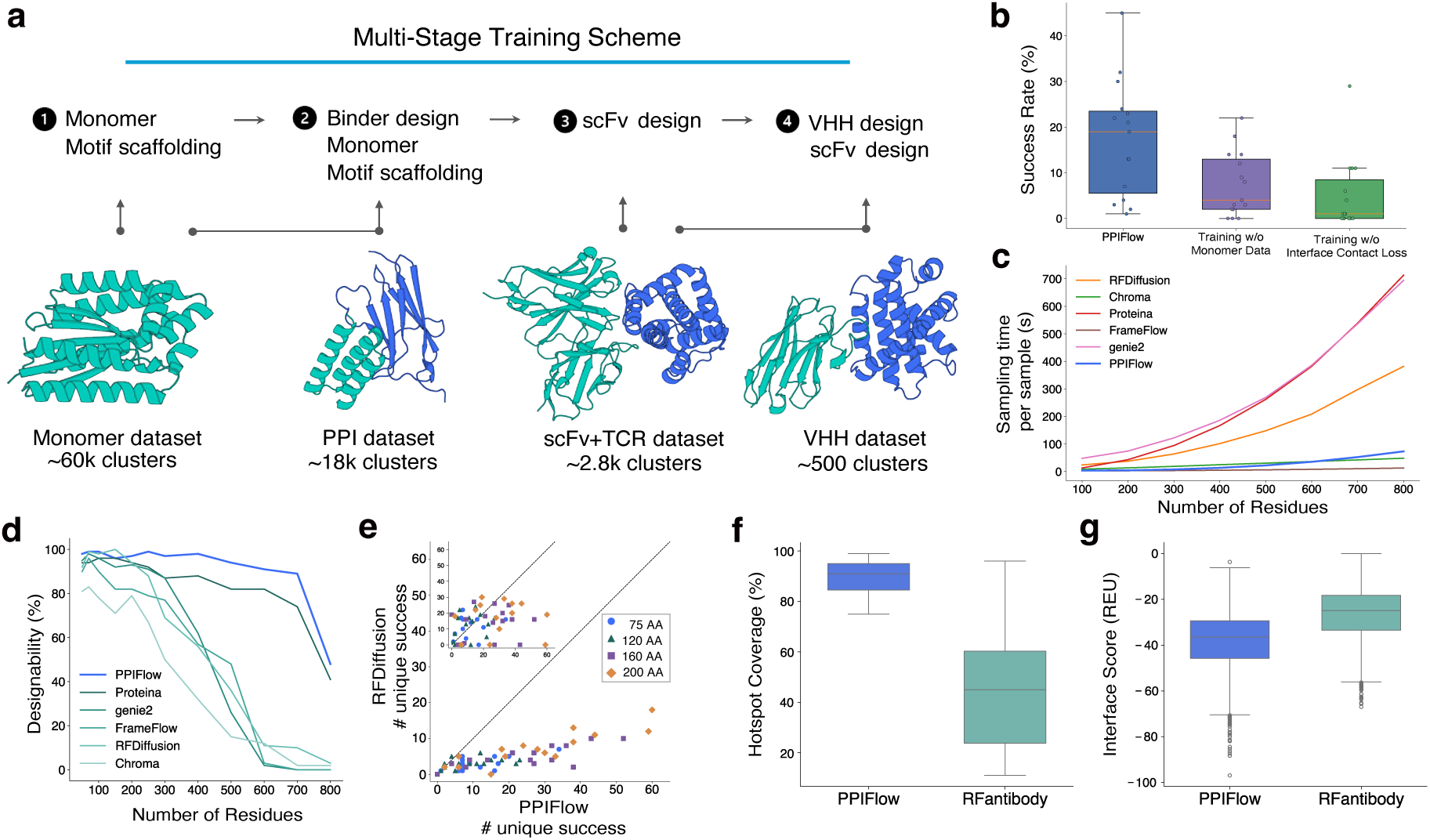
Training curriculum and *in silico* performance benchmarks. a, Multi-stage training scheme of PPIFlow. The foundation model is first trained on a monomeric protein dataset for structure recovery and scaffolding. Subsequent fine-tuning stages progressively adapt the model to protein–protein interactions (PPI), scFv generation, and VHH design, leveraging both general and antibody-specific datasets to preserve framework integrity. b, Ablation analysis of the training objective, highlighting the necessity of monomer dataset integration and interface contact loss for optimal binder design. c, Inference efficiency comparison of PPIFlow against representative generative architectures. d, Designability benchmarks for monomers. e, Designability benchmarks for binders, inset: comparison between PPIFlow (x-axis) & BindCraft (y-axis). f–g, VHH design quality metrics, including predicted hotspot coverage (**f**) and Rosetta interface scores (**g**) for generated antibody–antigen complexes.

Training follows a staged curriculum that adapts the model from general backbone generation to constrained PPI design (Fig. 2a). Optimization relies on a flow-matching objective [17] supplemented by auxiliary geometric loss terms to promote interface stability. We validated these design choices through ablation studies, and demonstrated that both staged curriculum and geometric constraints are critical for achieving robust binder generation (Fig. 2b).

### 2.2 PPIFlow generates high-designability proteins

PPIFlow performs robustly across five generative tasks. For monomers (up to 700 residues), the model achieved *>*90% design success (scRMSD *<* 2 Å vs. ESMFold) while maintaining structural novelty and diversity (Fig. 2d; Fig. S1) [23, 24, 10, 18, 13]. In motif scaffolding, PPIFlow outperformed state-of-the-art methods on MotifBench (score 42.38 vs. 28.6 for RFdiffusion) [25], solving challenging multi-segment cases (e.g., 4JHW, 4XOJ) where prior approaches failed (Fig. S2) [10, 26].

We further validated PPIFlow on minibinder design (75/120/160/200 residues) against 15 diverse targets. The model surpassed recent open-source methods like RFdiffusion, BindCraft and BoltzGen in design success, novelty, and diversity (Fig. 2e and Fig. S3) [12, 27]. Notably, PPIFlow shows a pronounced advantage for larger binders, as reflected by both AF3Score-based designability and Rosetta-derived interface energetics. This length-dependent improvement likely stems from the multi-stage training strategy combined with PPIFlow’s outstanding monomer designability (Fig. S3).

In VHH and antibody design, PPIFlow generated diverse CDRs while preserving input frameworks (median RMSD 0.18 Å) (Fig. S4, S5). The model targeted epitopes precisely (median hotspot coverage *>* 0.9) (Fig. 2f), yielding complexes with well-packed, high-quality interactions (Fig. 2g).

### 2.3 High-affinity design via interface rotamer enrichment and partial flow

While generative models have streamlined binder design [28], achieving picomolar affinity typically requires experimental optimization. To bridge this gap, we developed a pipeline integrating interface rotamer enrichment with partial flow refinement (Fig. 1c). In the **initial generation stage**, PPIFlow generates backbones targeting a predefined epitope (Fig. 1b). We design sequences using ProteinMPNN [29] or AbMPNN [30] and filter candidates via AF3Score to retain structurally plausible designs.

During maturation, we identify energetic “anchor” residues (*<* −5 Rosetta Energy Units) on the interface [31]. We fix these rotamers while perturbing the backbone to an intermediate flow state (*t* = 0.6). Crucially, this plasticity allows the backbone to escape local minima and resolve steric clashes imposed by the anchors, reshaping the binding interface rather than merely repacking side chains [32]. We then regenerate perturbed regions and redesign unconstrained sequences (*T* = 0.1). This active remodeling accommodates enriched residues and significantly improves binding compatibility (Fig. S3). Matured designs are re-evaluated via AF3Score, Rosetta Interface Analyzer, and template-free AlphaFold3. We prioritize the top 30 candidates, selected by a composite score (AF3 ipTM × 100 − Rosetta interface score), for experimental validation.

### 2.4 *De novo* design of high-affinity mini-binders

To experimentally validate the pipeline, we designed *de novo* binders against seven therapeutically relevant targets: IL7RA, IFNAR2, IL17A, PD-L1, TRKA, PDGFR, and VEGFA (Fig. 3a). Spanning diverse structural classes and functional roles in oncology and autoimmune disease, these targets provide a stringent benchmark. We characterized 30 candidates per target; all 210 expressed successfully, with 76 (36.2%) binders showing affinities better than 1 *µ*M by Bio-Layer Interferometry (BLI) (Fig. 3b,c; Fig. S6; Table S7), spanning diverse structures and sequences (Table S6,S7). Notably, we identified picomolar-affinity binders for six of the seven targets without post-design experimental optimization.

**Figure 3:**
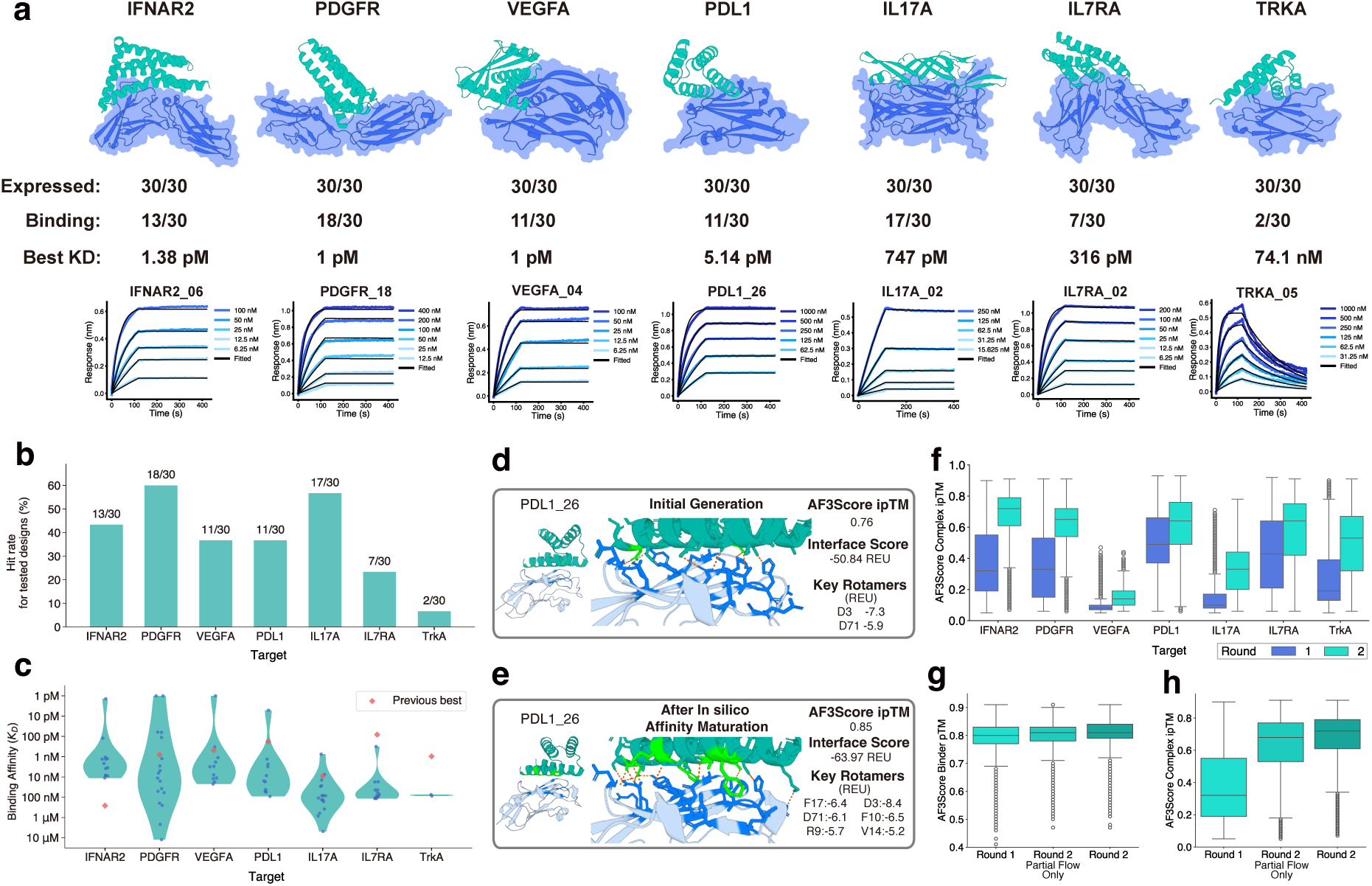
Experimental validation and high-affinity binder generation. a, Structural models of the highest-affinity *de novo* binders for each target. Accompanying data include expression success, binding hit rates, peak affinities, and representative BLI sensorgrams. b–c, Experimental performance across seven therapeutic targets, showing (**b**) hit rates and (**c**) the distribution of binding affinities (violin plots), demonstrating consistent picomolar-to-nanomolar potency. d–e, Illustration of interface side-chain packing after (**d**) initial generation and after (**e**) *in silico* affinity maturation for PDL1 26. f, Comparison of AF3Score profiles between initial Round 1 designs and matured Round 2 designs, illustrating the systematic quality improvement achieved through *in silico* maturation. g–h, Performance comparison of design strategies for IFNAR2. Metrics for AF3Score pTM (**g**) and ipTM (**h**) demonstrate that interface rotamer enrichment combined with partial flow refinement markedly outperforms both Round 1 baselines and partial flow alone.

Success stems largely from the *in silico* maturation stage. Structural comparisons reveal that interface rotamer enrichment followed by partial flow creates compact, highly connected interfaces featuring specific hydrogen bonds, salt bridges, and hydrophobic interactions formed during refinement (Fig. 3d,e; Fig. S9). Across all targets, this maturation systematically improved folding confidence and interface quality, significantly increasing AF3Score pTM and ipTM values (Fig. 3f). In the case of IFNAR2, interface rotamer enrichment markedly outperformed baseline pipelines, yielding superior interface metrics (Fig. 3g,h).

### 2.5 *De novo* design of high-affinity VHHs

We extended PPIFlow to the *de novo* generation of single-domain antibodies (VHHs) against eight therapeutic targets: CCL2, HNMT, PDGFR, 1433E, BHRF1, S100A4, EFNA1, and IL13. To prioritize developability, we sampled frameworks from five therapeutically validated scaffolds, generating and maturing CDRs via the PPIFlow-VHH pipeline (see Supplementary Information).

BLI characterization of 240 candidates (30 per target) confirmed 100% expression success. We identified binders for seven of the eight targets, with several achieving exceptional potency without optimization. Notably, VHH CCL2 21 reached 250 pM affinity, while binders for HNMT, PDGFR, and 1433E achieved single-digit nM affinities (Fig. 4a). The overall hit rate was 33.8% (81/240), with 27.9% (67/240) showing affinities better than 1 *µ*M, and these successful designs span diverse structures and sequences (Fig. 4b,c; Fig. S7; Table S6,S8).

**Figure 4:**
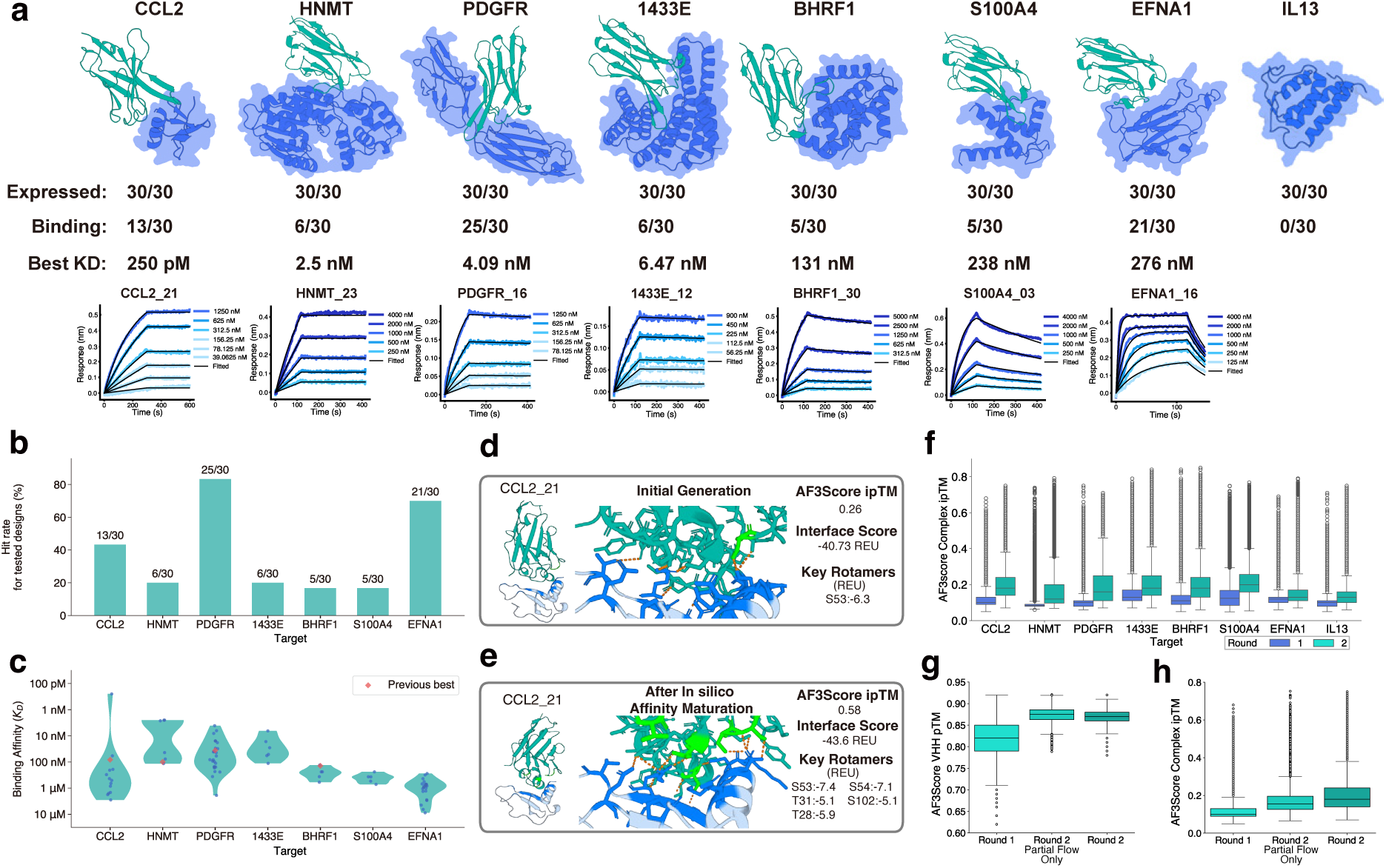
Computational design and experimental validation of high-affinity VHHs. a, Structural models of the highest-affinity *de novo* VHHs for each target, paired with expression data, binding hit rates, peak affinities (*K_D_*), and representative BLI sensorgrams. b–c, Experimental performance across VHH targets, showing (**b**) hit rates and (**c**) affinity distributions (violin plots), highlighting the discovery of sub-nanomolar binders for CCL2, PDGFR, and 1433E. d–e, Illustration of interface side-chain packing after (**d**) initial generation and after (**e**) *in silico* affinity maturation for CCL2 21. f, Comparison of AF3Score profiles between initial (Round 1) and matured (Round 2) VHH designs, demonstrating consistent improvement in predicted interface quality across all targets. g–h, Evaluation of design strategies for the CCL2 target. AF3Score pTM (**g**) and ipTM (**h**) metrics confirm that the full maturation protocol, integrating interface rotamer enrichment with partial flow, significantly outperforms baseline generative and refinement-only pipelines.

As with general protein binders, AF3Score analysis confirms that *in silico* maturation, combining interface rotamer enrichment with partial flow, was critical. This stage systematically improved structural confidence and interface quality (Fig. 4f), evidenced by compact side-chain packing and energetically favorable interactions (Fig. 4d,e; Fig. S10). Benchmarks on CCL2 show that interface rotamer enrichment significantly outperforms baselines, yielding higher AF3Score ipTM values and superior experimental outcomes (Fig. 4g,h).

## 3 Discussion

The ability of PPIFlow to generate picomolar affinity binders and VHHs purely *in silico* marks a shift from “design-then-screen and optimize” toward de novo binder generation. Traditionally, bridging the “affinity gap” required massive experimental libraries or directed evolution. Recent retrospective analyses have shown that AlphaFold3 interface confidence scores are strongly predictive of experimental binding success and can therefore serve as an effective structural consistency filter [11]. By incorporating AF3Score, a score-only adaptation of AlphaFold3, PPIFlow bypasses MSA construction and iterative recycling, accelerating evaluation by approximately two orders of magnitude compared to standard inference [20]. This enables rapid, high-fidelity candidate prioritization without the computational burden of traditional protocols.

A central conceptual advance of this work is the integration of global backbone topology sampling with local interface optimization. While flow matching explores diverse architectures, interface rotamer enrichment acts as a computational analogue of affinity maturation. This hybrid approach resolves the tension between broad fold exploration and the stringent geometric requirements of high-affinity binding, addressing the observation that high in silico designability alone does not guarantee experimental success [28, 27].

Although PPIFlow outperforms existing methods such as BindCraft on backbone feasibility metrics, its superior experimental hit rates also stem from the pipeline’s synergy. Furthermore, PPIFlow significantly reduces the computational burden associated with binder design. Unlike protocols such as BindCraft that typically necessitate computationally expensive full AlphaFold predictions for filtering, our workflow leverages AF3Score to rank complexes directly, accelerating candidate prioritization by approximately 100-fold. Unlike standard generation methods, PPIFlow robustly accommodates the constraints imposed during “Interface Rotamer Enrichment”, allowing the subsequent “Partial Flow” stage to refine packing without breaking the fold. Thus, it is not the generator in isolation, but the holistic integration of pairformer, energy-guided anchoring, and flow-based refinement that bridges the gap between computational plausibility and picomolar potency.

Limitations remain. PPIFlow currently operates as a backbone-only generator, meaning side-chain packing is handled post-generation [33]. While our maturation pipeline mitigates this, a fully atomistic generative framework could more natively capture the highly specific hydrogen-bonding networks and hydrophobic packing required for certain complex interfaces. We also observed target-dependent variation in success rates. For example, IL13 remained challenging. In such cases, the interface rotamer enrichment stage may fail to identify stable energetic anchors (*<* -5 REU), preventing effective partial flow refinement. Consistent with this, VHHs generated against IL13 showed systematically lower confidence scores (Fig. 4f).

Finally, the current model is limited to protein–protein complexes; extending the framework to nucleic acids, glycans, and small molecules is essential. Nevertheless, PPIFlow establishes a generalizable platform for rapid, purely computational binder design, significantly reducing reliance on experimental iteration.

## 4 Data and materials availability

The code and relevant data for PPIFlow will be available at https://github.com/Mingchenchen/PPIFlow. AF3Score is available at https://github.com/Mingchenchen/AF3Score: A Score-Only Adaptation of AlphaFold3 for Biomolecular Structure Evaluation. All relevant data will be available at Zenodo.

## Supplementary Information

## Appendix A Supplementary Methods

### A.1 Notation

We denote the number of residues in a protein structure by *N*, and each residue is represented by a *frame T* = (*x, r*) ∈ SE(3), where *x* ∈ ℝ^3^ denotes a translation vector and *r* ∈ SO(3) a rotation matrix. Our model generates backbone atoms, i.e. Ω = {*N, C, C_α_, O*}, and we denote by *z* ∈ ℝ^3^ the global coordinates of a single atom. For binder design tasks, we denote the target backbone atoms and the binder backbone atoms by *B* and *T* respectively. The set of atom pairs between the target and the binder whose backbone atoms are separated by a distance less than 10 Å is denoted by *P* which is used in loss computation.

We establish the following notation for operators employed in the subsequent algorithms. Capitalized operator names are exclusively reserved for modules containing learnable parameters. For instance, LINEAR signifies a parameterized linear transformation, defined by a weight matrix *W* and a bias vector *b*. The term LAYERNORM specifically denotes layer normalization applied across the channel dimensions. Conversely, lowercase names are utilized for functions that operate without learnable parameters (e.g., `sigmoid`, `softmax`) and standard non-parameterized mathematical operations (e.g., `sin`, `cos`, and `exp`).

### A.2 Datasets

#### A.2.1 Data Curation

We curated a unified structural dataset comprising three complementary data sources: monomeric protein structures, protein–protein interaction (PPI) complexes, and antibody-like binders, including VHH and scFv.

Monomeric protein structures were obtained from two sources: protein domains curated in the CATH database [34] and single-chain structures from the RCSB Protein Data Bank (PDB) [35]. We filtered the data based on the following criteria:

- non canonical amino acid ratio less than 20%.
- Sequence length is between 25 and 512.
- Coil percent is less than 0.8
- Radius of gyration is within the lowest 96% of the distribution.

Sequence redundancy was reduced by clustering all sequences using MMseqs2 [36]. After filtering and deduplication, the monomer dataset comprised approximately 60k clusters, spanning a broad range of lengths and folds. This dataset was used primarily for unconditional structure learning, enabling the model to learn general protein structures.

We curated a protein–protein interaction (PPI) dataset consisting of both chain–chain interactions (CCI) and domain–domain interactions (DDI). We filter the data based on the following criteria:

- number of contact pairs larger than 80 where contact pair is defined as interchain residues within the distance of 10 Å.
- non canonical amino acid ratio less than 20%.
- structure similarity between target and binder is less than 0.6.
- binder radius of gyration divided by length less than *k*, where *k* = 0.15 if binder size is larger than 100, *k* = 0.16 if binder size is between 70 and 100, *k* = 0.18 if binder size is less than 70.
- binder main chain SASA ration is less than 17.
- Sequence length is less than 570.
- Binder sequence length is less than 256 and target sequence length is larger than 30.

After filtering and redundancy removal, the combined CCI + DDI dataset contained approximately 18k interaction pairs. This dataset was used to train the model to capture general interface geometry and inter-chain spatial organization, which are essential for downstream binder generation.

**Table S1:**
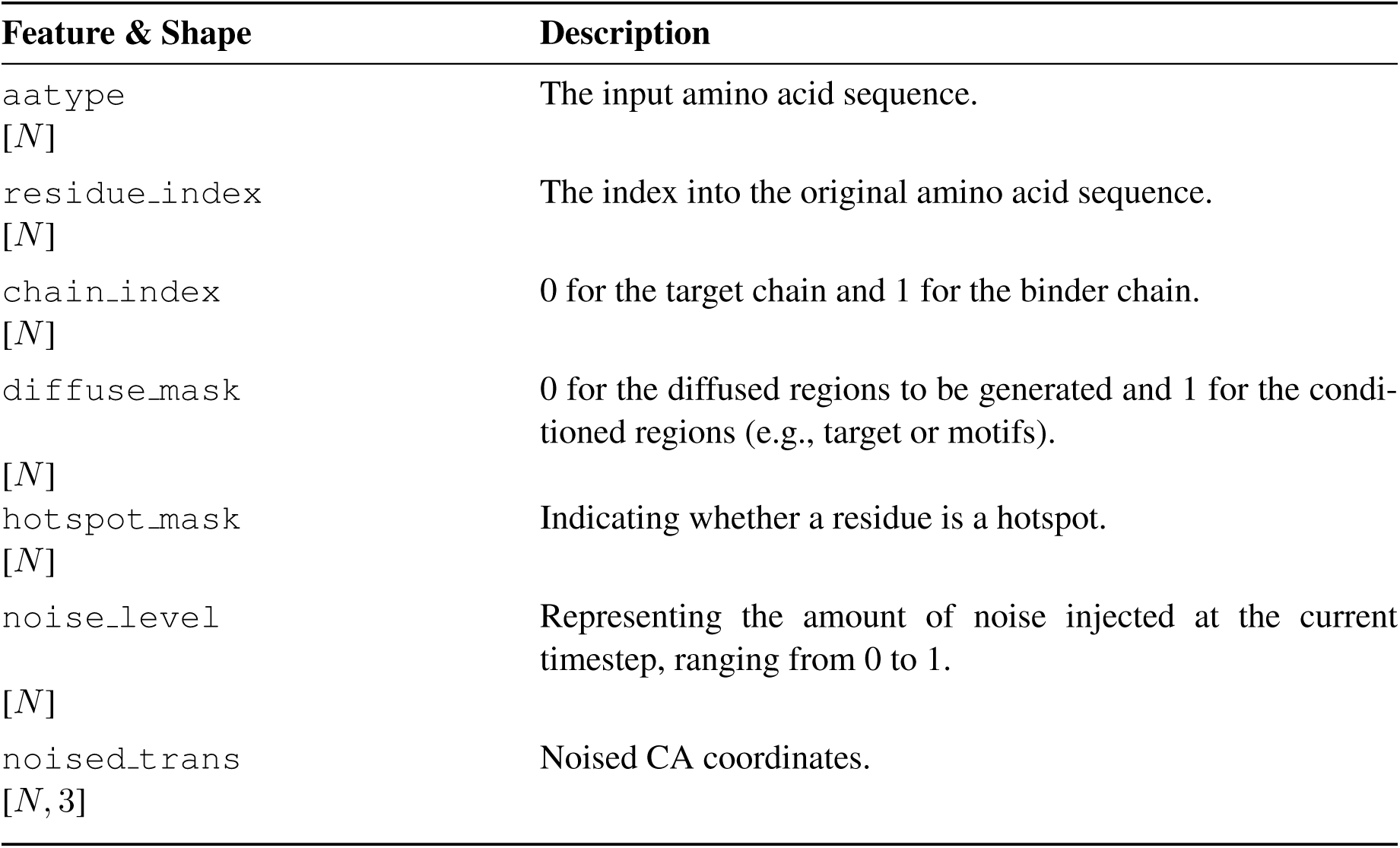
Input features to the model. *N* is the number of residues.

To specialize the model for antibody-like binder design, we curated a dataset of antibody, VHH and TCR structures from SAbDab [37]. The dataset comprised approximately 6.9k antibody structures, 1.2k VHH structures, and 350 TCR structures, clustered into 2.6k, 500 and 200 clusters, respectively.

#### A.2.2 Input Features

In Table S1, we list the input features to the model. These features will be embedded into the single embedding network and pair embedding network.

### A.3 Input Embeddings

Initial embedding details are presented in Algorithm 1 and Algorithm 2.

**Algorithm 1:**
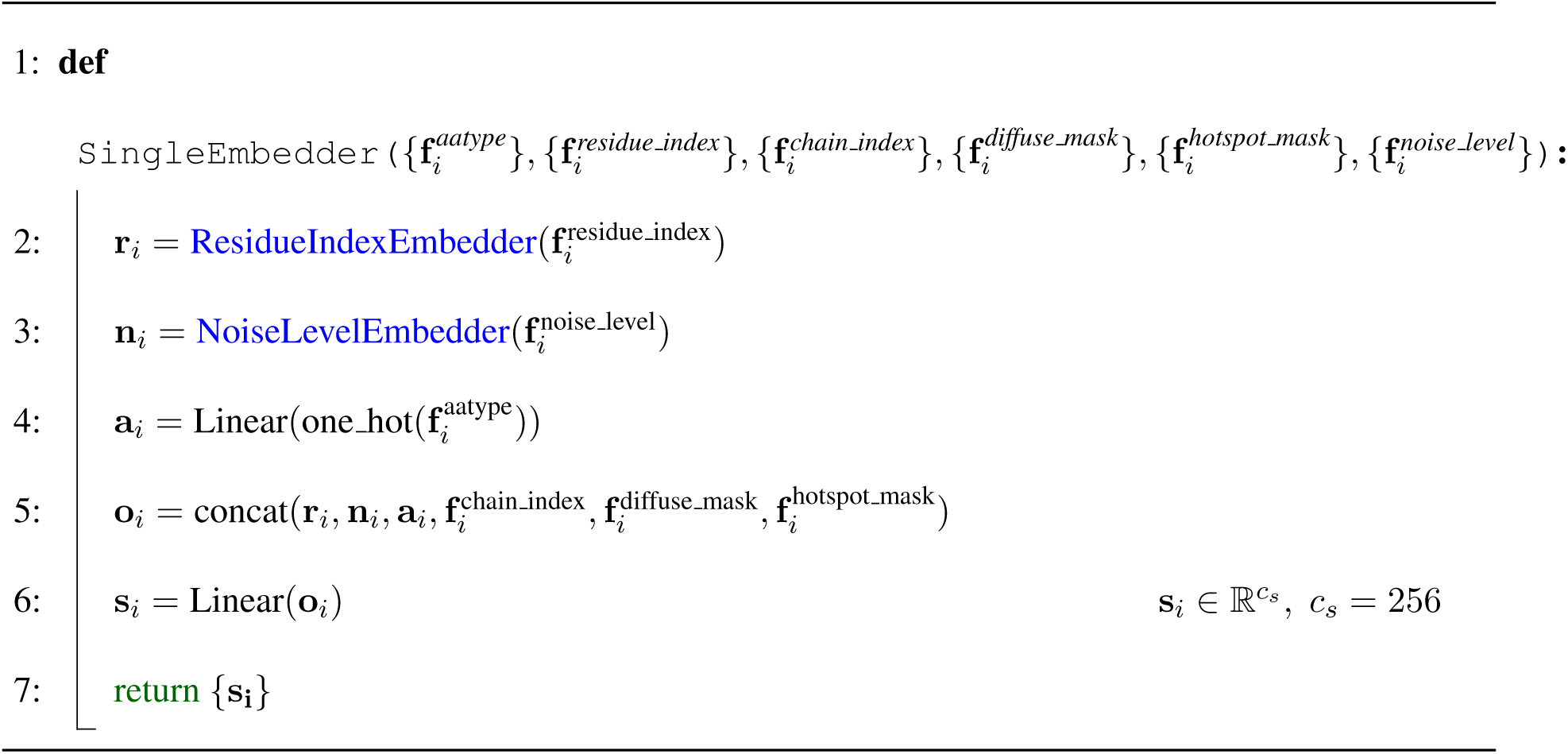
Embeddings for initial single representations.

**Algorithm 2:**
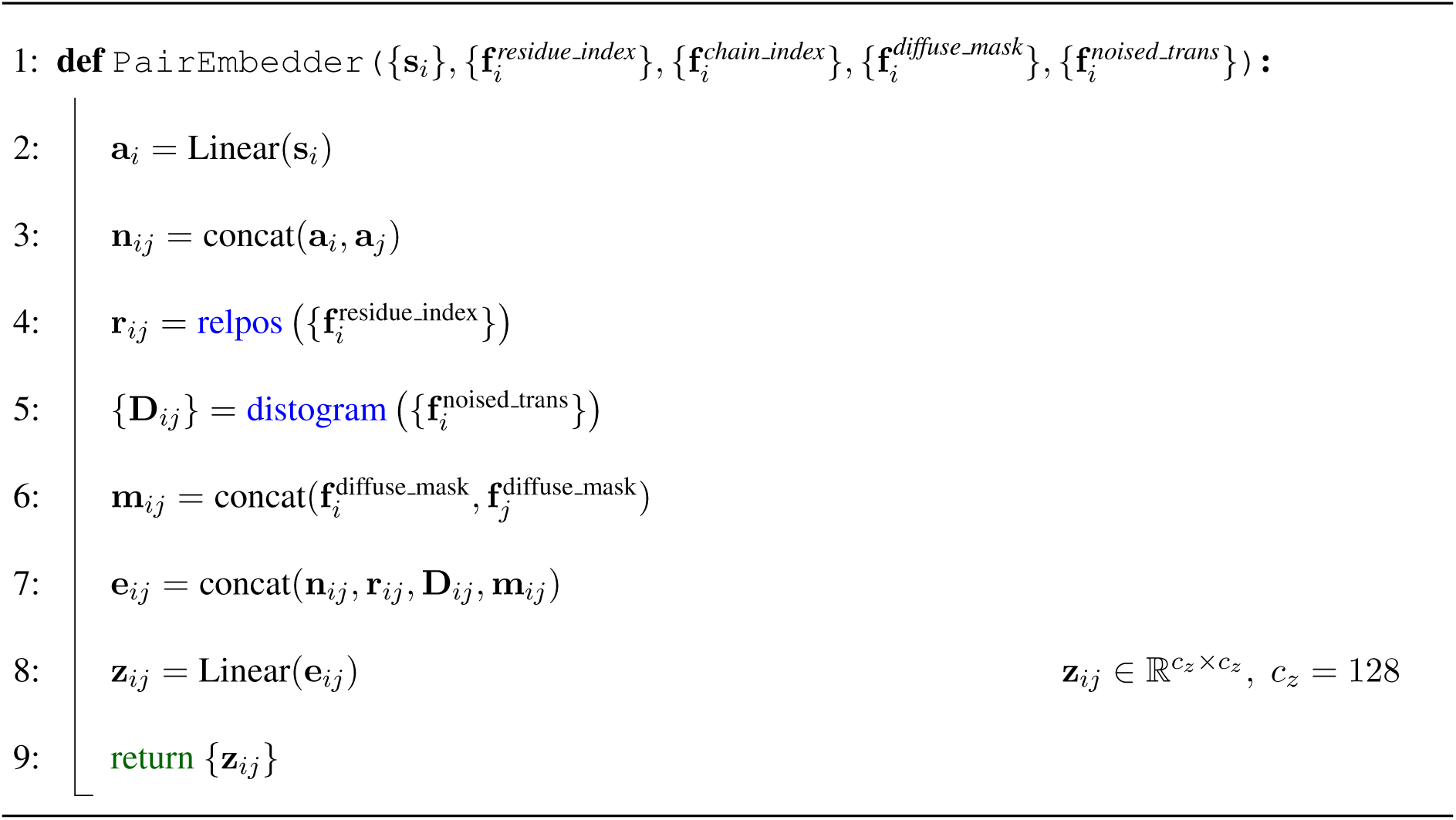
Embeddings for initial pair representations.

**Algorithm 3:**
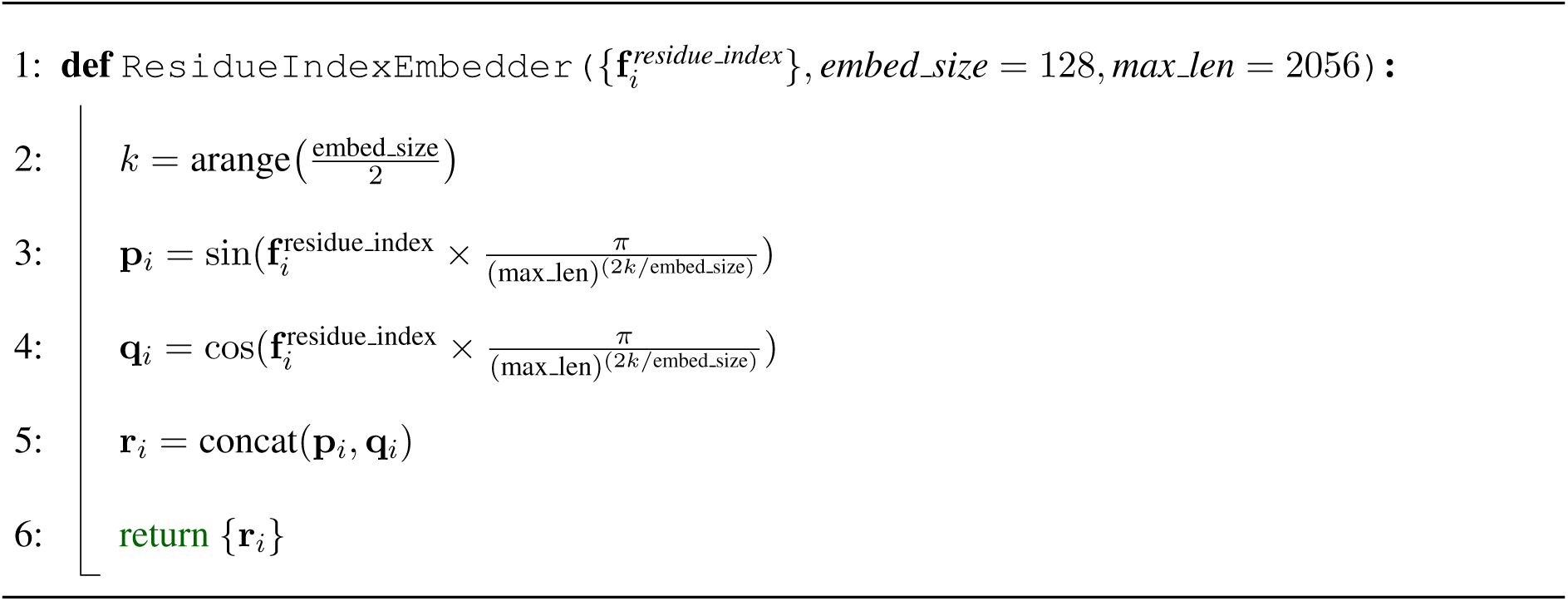
Embeddings for residue index.

**Algorithm 4:**
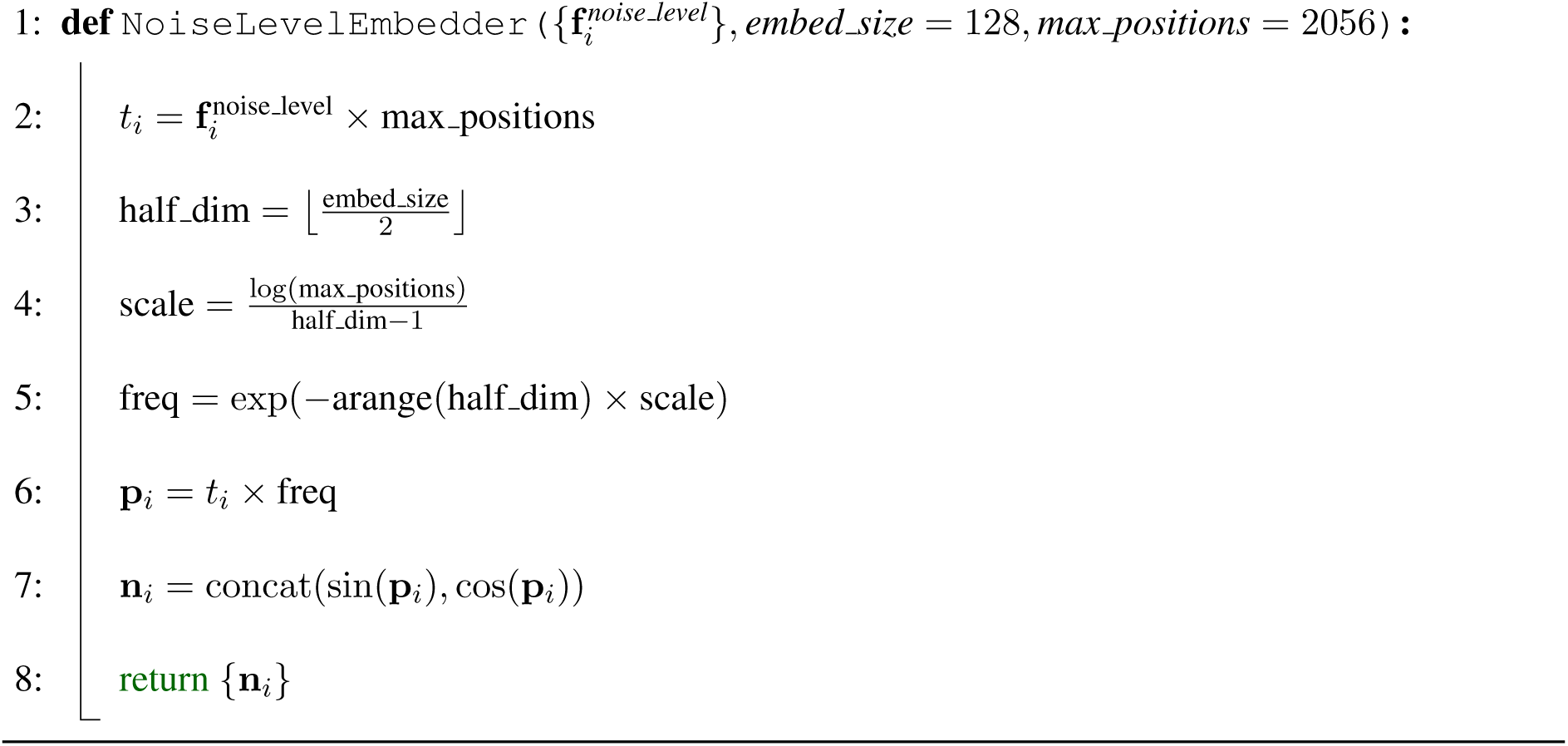
Embeddings for noise level.

**Algorithm 5:**
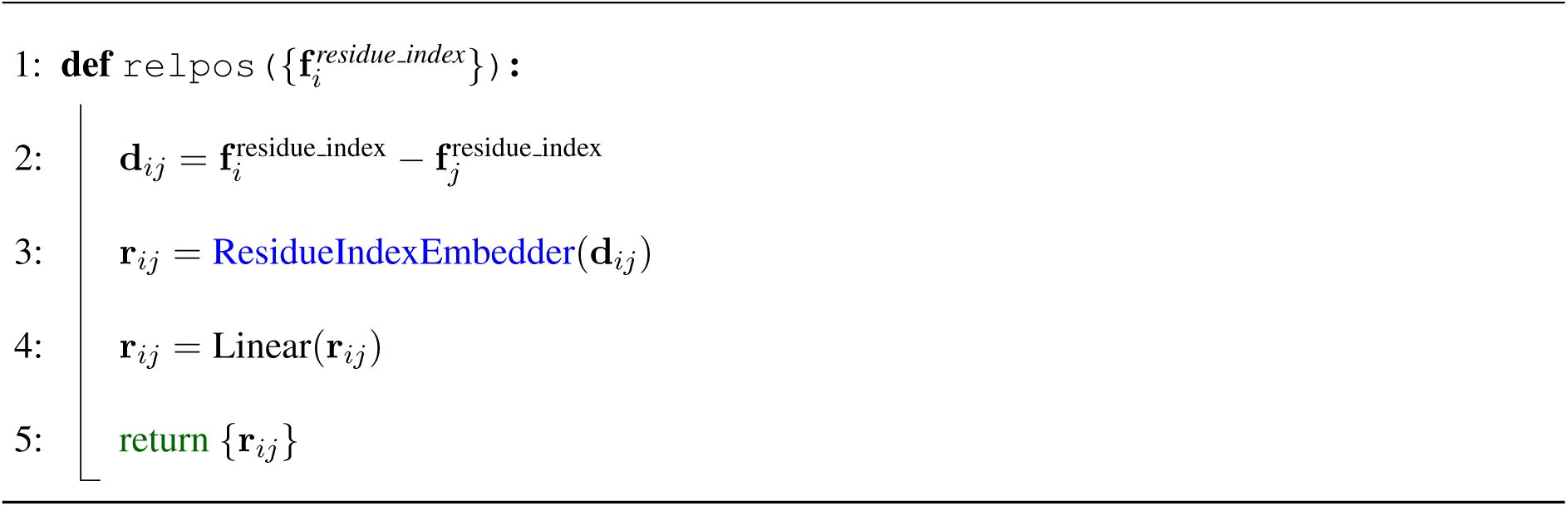
Relative position encoding.

**Algorithm 6:**
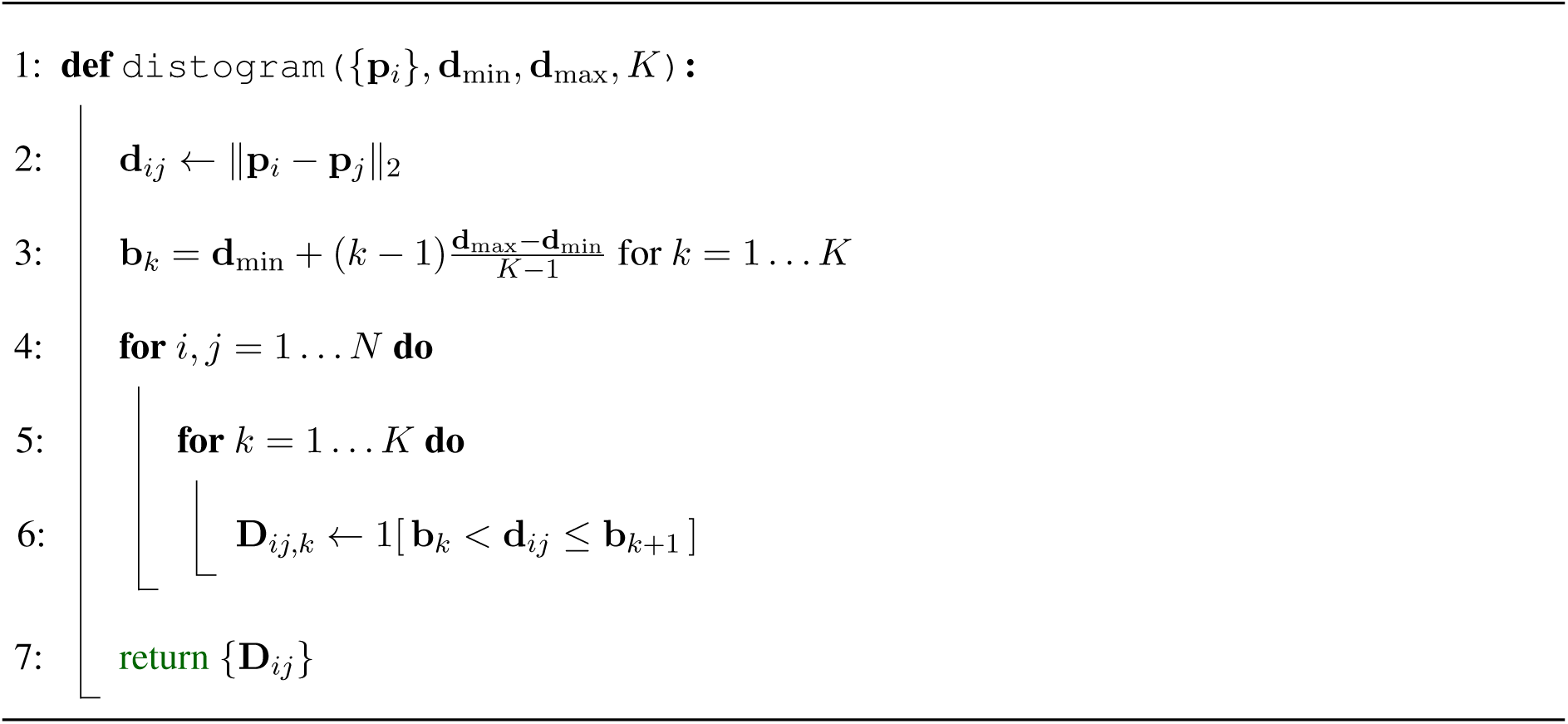
Compute distogram from atomic coordinates.

### A.4 Model Architecture

Our model follows a hierarchical architecture that integrates both sequence-level and structure-level reasoning. Specifically, we employ two key components that have become standard in protein structure modeling: the **Pairformer** and the **Invariant Point Attention (IPA)** module.

The Pairformer module is used to jointly model residue-wise and pairwise representations through interleaved attention and transition layers. It allows effective information exchange between the one-dimensional sequence features and the two-dimensional pairwise geometric features, enabling the network to reason about both local and long-range residue interactions. We adopt the standard Pairformer design as introduced in AlphaFold-style architectures, without modification. We use 4 Pairformer blocks in our model.

Following AlphaFold [21], we use the Invariant Point Attention module to transform pairwise relational information into 3D coordinate updates. The IPA performs attention over residue embeddings in SE(3)-invariant space, ensuring that the resulting structural representations are equivariant to rotations and translations. This design allows the model to learn spatial relationships in a physically meaningful and symmetry-preserving manner. Our model includes 6 IPA layers.

Our model consists of 23.2M parameters. Together, these modules enable our model to integrate sequence context, pairwise geometry, and spatial reasoning in a unified framework for protein structure generation and refinement.

### A.5 Loss Functions

#### Flow matching loss in SE(3)

Following the flow-matching framework, the model learns to predict the continuous velocity field that transforms a noisy structure toward its clean counterpart. Each residue is represented in SE(3) space by a translation vector *x* ∈ ℝ^3^ and a rotation matrix *r* ∈ SO(3).

Given a clean structure **T**_1_ and a noisy structure **T***_t_* sampled along the flow trajectory, the target velocity is defined as the time derivative of the flow:

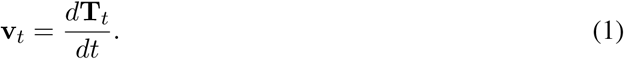

The SE(3) flow-matching loss minimizes the discrepancy between predicted and ground-truth velocities on both translational and rotational components:

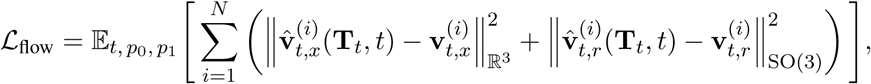

where *p*_0_ and *p*_1_ denote noise distribution and data distribution respectively, *p*_0_ = *N* (0*, I*_3_)⊗U(SO(3)), *t* ∼ U([0, 1]).

Following FrameFlow [18], the model predicts the clean frames **T**_1_ given the corrupted frames **T***_t_* at time *t*, and the velocity can be approximated as

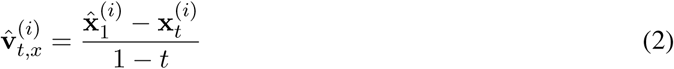

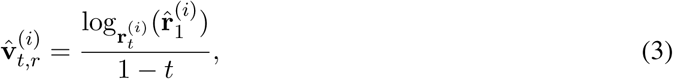

where log 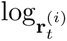 is the logarithmic map on SO(3). As a result, the objective can be reparameterized as

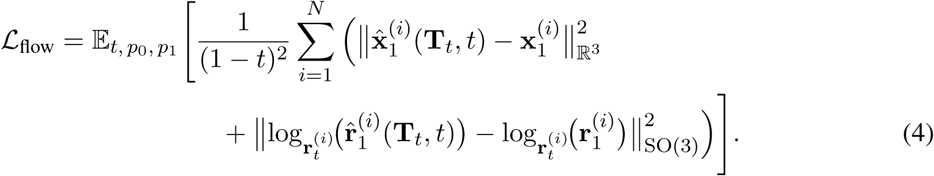

#### Interface contact loss

We introduce interface contact loss to improve the learning of target to binder interaction. Since we observed that some sampled results positioned binders too far from the target, leading to weak contacts, we designed this interface contact loss to improve the quality of interfaces. This loss is defined on residue pairs across binder and target. We examine on interchain residue pairs within 10 Å in native structures, and compare the predicted pairwise distances of these pairs against true pairwise distances. Let 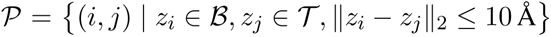, where *z_i_* and 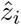 are the ground-truth and predicted coordinates of backbone atoms respectively. Then the interface contact loss can be written as

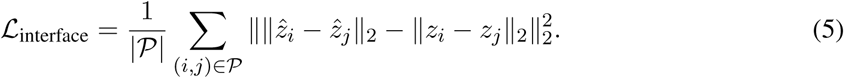

#### Backbone atom loss

The backbone atom coordinates are reconstructed from the predicted translation and rotation vectors, and we compute the mean squared error (MSE) between the predicted and ground-truth coordinates of the backbone atoms. The backbone atom loss is defined as

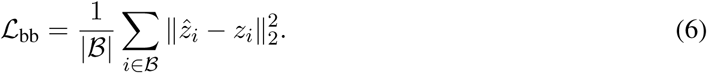

#### Binder pairwise distance loss

By minimizing binder pairwise distance loss, the model is trained to accurately predict the spatial relationship (the pairwise distances) between residues in the binder, which is crucial for generating physically realistic proteins. This loss is defined as

**Algorithm 7:**
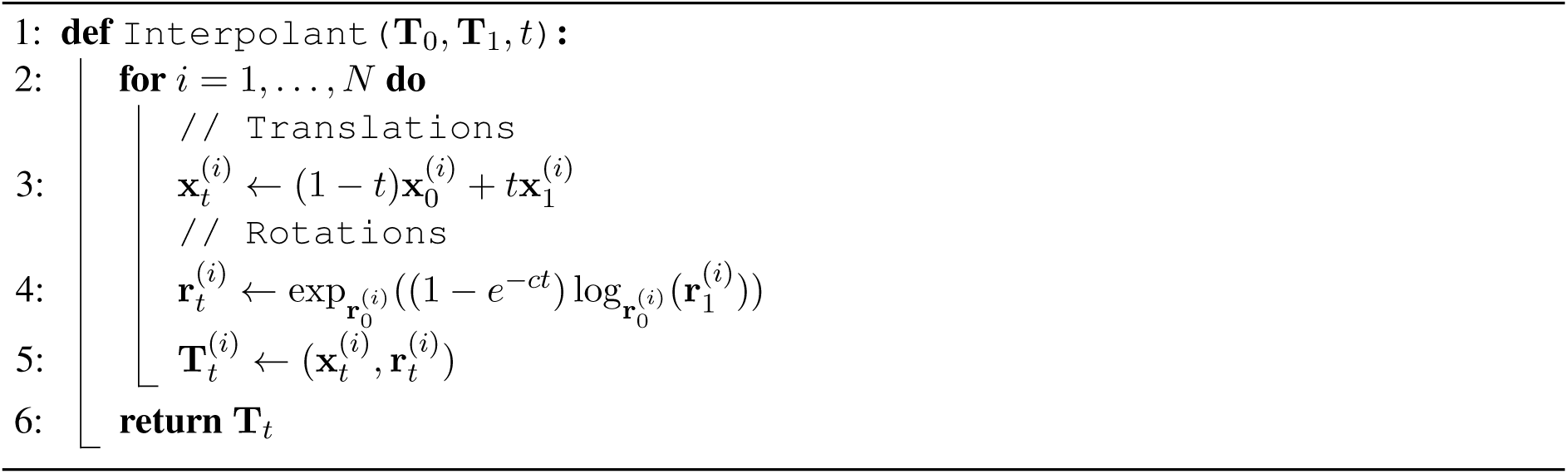
SE(3) flow matching interpolation.

**Algorithm 8:**
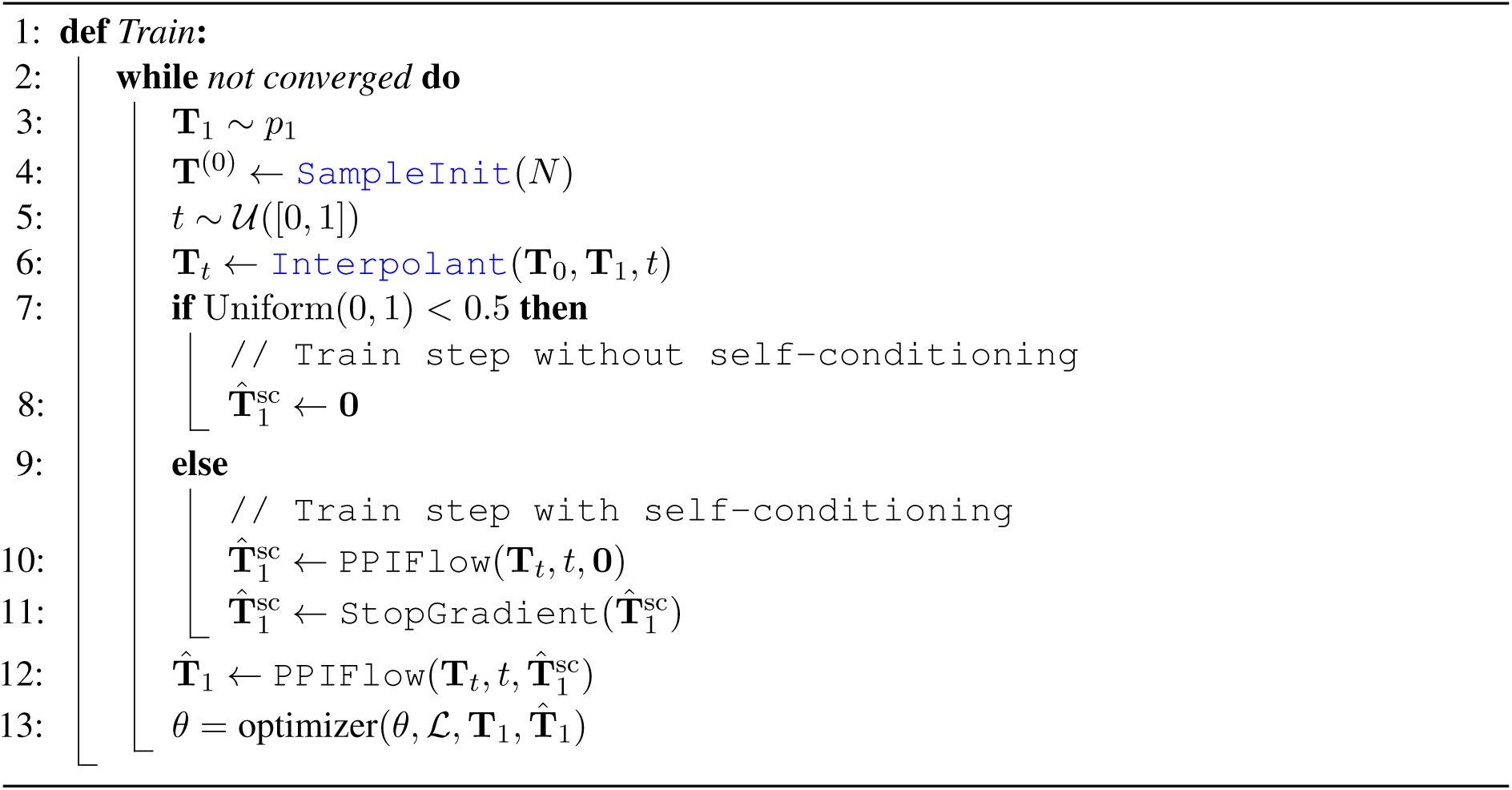
PPIFlow training.

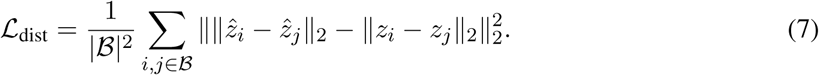

### A.6 Training and Inference Details

To enable our model to support diverse protein generation tasks—including monomer generation, motif scaffolding, binder design, scFv generation, and VHH generation—we adopt a multi-stage training strategy. The model is trained progressively in four stages, with each stage initializing from the checkpoint of the previous one. This staged training allows the model to first acquire general protein structural priors and then gradually specialize toward increasingly complex multi-chain and antibody-specific generation tasks. Algorithm 8 summarizes the training procedure. Below we introduce each training stage in detail.

#### The first stage: monomer pretraining with motif scaffolding

In the first stage, the model is trained on a large-scale monomeric protein dataset to learn general protein structures. During this stage, we jointly train the model on two tasks:

- Monomer generation, where the model learns to generate full-length single-chain protein structures.
- Motif scaffolding, where a subset of residues corresponding to functional or structural motifs is provided as conditioning input, and the model generates a complete protein scaffold consistent with the motif geometry.

During this stage, 20% of the samples are generated unconditionally, while for the remaining 80%, 10%–90% of residues are randomly selected as motifs, and the corruption is applied only to the non-motif residues. By mixing motif scaffolding with unconditional monomer generation, the model learns to preserve local structural constraints while maintaining global fold coherence. This stage establishes a strong structural prior and serves as the foundation for all subsequent finetuning stages.

#### The second stage: binder design finetuning

In the second stage, the model is finetuned for binder design where the model generates a binder protein conditioned on a target protein. Training data consist of a mixture of monomer structures and protein–protein complex (PPI) structures. The training objective is formulated as a multi-task mixture of: monomer generation, motif scaffolding, and binder design. Specifically, we downsample monomer data to 25% since the total amount of monomers is much larger than that of complexes. If a sample is monomer, then we follow the motif sample strategy used in the first stage. If a sample is a complex, then we generate the binder unconditional for 50% of the time and with motifs otherwise. For binder design task, we define target interface as residues whose CA position is within 10 Å to the binder. During training, we randomly select 0–20% percent of the interface residues as hotspots, and at least 3 hotspots are selected.

Including monomer and motif scaffolding tasks alongside binder design helps prevent catastrophic forgetting and stabilizes training (refer to ablation study). This stage equips the model with the ability to reason about inter-chain geometric constraints and protein–protein interfaces, which are essential for downstream antibody-related tasks.

#### The third stage: scFv finetuning with framework conditioning

In the third stage, we finetune the model for single-chain variable fragment (scFv) generation, conditioned on a given antigen. Training data are collected from SAbDab and TCR. In this stage, the model is trained to generate an scFv structure given an antigen structure as input.

To support antibody–antigen systems and framework-aware generation, we introduce two key architectural modifications: chain group encoding and framework-aware pair embedding. Antibody–antigen complexes often contain multiple chains. Instead of using raw chain IDs, we replace them with chain group IDs, where all antigen chains belong to one group and all antibody chains belong to another. This modification enables the model to better distinguish inter-group (antigen–antibody) interactions from intra-group interactions while remaining invariant to arbitrary chain labeling. To explicitly control framework generation, we provide a reference framework structure as part of the input. We augment the pair embedding module with an additional layer that extracts pairwise distance features within the reference framework. These features are injected into the model’s pair representations, allowing the model to condition on the reference framework geometry.

As a result, the model is trained to generate both the framework and CDR regions simultaneously, while ensuring that the generated framework remains highly consistent with the provided reference. This design allows controlled antibody generation without freezing or hard-constraining the framework coordinates.

#### The fourth stage: VHH finetuning

In the final stage, the model is finetuned for VHH (single-domain antibody) generation. Training data are drawn from SAbDab and TCR datasets, including both scFV and VHH. This stage reuses the architectural modifications introduced in the third stage, including chain group encoding and framework-aware pair embeddings. Because VHHs consist of a single variable domain, this stage further refines the model’s ability to generate compact, stable antibody structures while preserving antigen-binding specificity.

**Algorithm 9:**
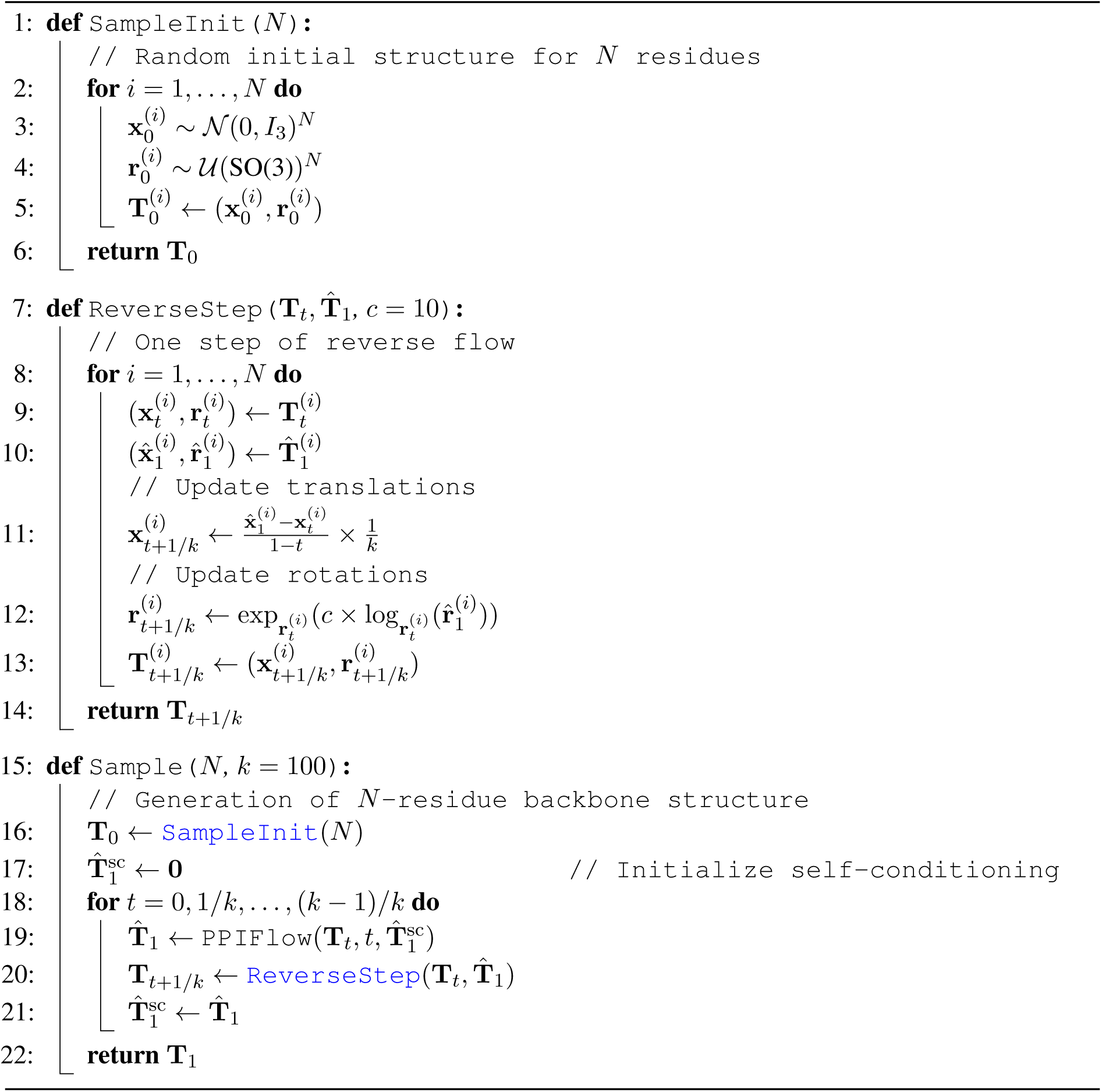
PPIFlow inference.

### A.7 Ablation Studies

To rigorously assess the contribution of key components in our binder design methodology, we conducted an ablation study focusing on two critical aspects: the training data strategy and the objective function.

#### Training data strategy ablation

Our primary training strategy involves jointly utilizing data from both binder design tasks and monomer structures. To evaluate the necessity of this data mixture, we performed an ablation where the model was trained exclusively on data relevant to the binder design task, omitting the monomer structure data. We denote this variant as w/o Monomer Data.

#### Loss function ablation

The full objective function incorporates an interface contact loss designed to explicitly guide the generation of favorable binding interfaces. We tested a variant, labeled w/o Interface Contact Loss, where this interface contact term was removed from the total loss during the training process.

The results demonstrate that both ablations lead to a degradation in the performance of the binder design task (Fig. 2b). Specifically, the w/o Monomer Data model exhibited a significant drop in pTM, indicating that the inclusion of monomer data is crucial for robust structural stability. Similarly, the w/o Interface Contact Loss variant showed diminished ability to generate high-quality interfaces, confirming that the explicit supervision provided by the interface contact loss is essential for optimizing inter-chain interactions.

These findings validate the necessity of both the mixed data training strategy and the interface contact loss term for achieving state-of-the-art results in protein binder design.

## Appendix B Design Workflow

### B.1 Binder design workflow

In the first design round, binder backbones were generated using PPIFlow and subsequently sequenced with ProteinMPNN. To balance sequence diversity and structural fidelity, ProteinMPNN was applied with a low sampling temperature (T = 0.2). As ProteinMPNN tends to under-sample bulky hydrophobic residues, such as phenylalanine, methionine, and tryptophan, which are often critical for stabilizing protein–protein interfaces, we adopted a dual-sequence design strategy. For each backbone, 8 sequences were generated using the vanilla ProteinMPNN checkpoint, while an additional 8 sequences were generated with bias toward large side-chain residues to enhance hydrophobic interactions at the interface. All designs were evaluated using AF3Score and filtered based on binder pTM and complex ipTM metrics to retain only structurally and functionally plausible complexes.

Candidates passing the first filtering stage were further optimized through an maturation process. Interface residues were analyzed using Rosetta residue-wise interaction energy calculations, and for each backbone, residues exhibiting strong energetic contributions (total interaction energy *<* -5 Rosetta Energy Units) across the 16 designed sequences were merged to form an interface rotamer enriched design. To further refine the structural context of these enriched interfaces, we applied partial flow, a flow-matching analogue of partial diffusion. In this step, key interface residues were fixed, remaining backbone is perturbed to an intermediate flow state (*t* = 0.6). The perturbed regions were regenerated using PPIFlow. The unconstrained regions were then redesigned using the ProteinMPNN soluble checkpoint to improve folding stability (T = 0.1).

The matured designs were re-evaluated using AF3Score and filtered again based on binder pTM and complex ipTM. Surviving candidates were further assessed using AlphaFold3, and Rosetta interface analyzer. Top 30 candidates defined by a composite score (AF3 ipTM*100 - Interface Score) were selected for experimental validation.

### B.2 VHH design workflow

For each target, 25,000 VHH backbone structures were sampled using PPIFlow-VHH. For each backbone, 8 CDR sequences were designed using ABMPNN with a sampling temperature of 0.5 to increase sequence diversity. All designs were evaluated using AF3Score, and candidates with ipTM *>* 0.2 were retained for the second-round optimization. Interface residues were then analyzed using Rosetta residue-wise interaction energy calculations to identify key residues for binding.

In the second-round, for each backbone, interface residues with strong energetic contributions, total interaction energy *<* −5 Rosetta Energy Units, across the 8 first-round sequences were merged to construct a set of enriched interface rotamers. Partial flow was subsequently applied to the remaining regions of the VHH backbone, while enriched key interface residues were held fixed, to regenerate 8 refined backbone conformations. For each refined structure, 4 sequences were redesigned using ABMPNN at a lower temperature (T = 0.1) to favor high-confidence sequence–structure compatibility.

All second-round designs were re-evaluated using AF3Score, and candidates with ipTM *>* 0.5 and pTM *>* 0.8 were subjected to further structural consistency validation with AlphaFold3. Designs were filtered based on agreement between AF3-predicted and designed structures (DockQ *>* 0.49), AF3 pTM *>* 0.8, and AF3 ipTM *>* 0.7. For each target, VHH candidates passing these filters were ranked by a composite score (AF3 ipTM*100 - Interface Score). Top 30 candidates were selected for experimental validation based on the composite score.

## Appendix C Experimental Materials & Methods

### C.1 Binder experimental methods

#### C.1.1 Gene Synthesis and Subcloning

The target DNA sequence was optimized for codons and synthesized by GenScript Biotechnology Co., Ltd. (Nanjing, China). The synthesized gene was subcloned into the pIVEX vector for protein expression in a cell-free system. Subcloning into the pIVEX vector yielded the following product: MSG-[protein]-GSGSHHWGSTHHHHHH, where the C-terminal SNAC cleavage tag and 6×His affinity tag are underlined, respectively.

#### C.1.2 Cell-Free Protein Expression

Cell-free protein synthesis reactions were assembled by combining S30 cell lysate, synthesis buffer, required enzymes, and plasmid DNA in a 24-deep-well plate. Each reaction had a final volume of 5 mL and was incubated at 25^◦^C for 16 hours. Subsequently, the reaction mixture was then centrifuged at 4000 rpm for 5 minutes and the supernatant was collected for purification.

#### C.1.3 Cell-Free Protein Purification and Analysis

Target protein was obtained by Ni column purification using Hamilton instruments (Hamilton, MICRO LAB Starlet). The purified protein was then dialyzed in the desired buffer (either PBS, pH 7.4 or 50 mM Tris, 150 mM NaCl, 10% Glycerol, pH 8.0). Before subpackage and storage, the target protein was sterilized through a 0.22 µm filter. Protein concentration was determined by the A280 protein assay using bovine serum albumin (BSA) as the standard. Protein purity was verified by standard SDS-PAGE electrophoresis. For SDS-PAGE analysis: samples were mixed with 5× reducing loading buffer (300 mM Tris-HCl pH 6.8, 10% SDS, 30% glycerol, 0.5% bromophenol blue, and 250 mM DTT). Proteins were separated on a 12% homogeneous SDS-PAGE gel (GenScript, Cat. No. M00668) and visualized accordingly.

#### C.1.4 Target protein preparation

Since the binders designed in this study were expressed with His-tag, target proteins were preferentially selected without a His-tag to avoid potential interference during affinity assessment. Most of these target proteins were obtained with an Fc-tag or in untagged forms. For IL-17A, commercially available proteins lacking alternative tags were unavailable; therefore, biotin-labeled, His-tagged IL-17A variants were purchased. In such cases, affinity measurements were performed using immobilized antigens to evaluate binder binding. All purchased target proteins were commercially available reagents with validated biological activity.

#### C.1.5 Binding characterization of binders

Initial screening of binder designs for target protein binding was performed using Biolayer Interferometry (BLI, Gator Bio). For target proteins without the His-tag (TrkA, IFNAR2, IL-7RA, PD-L1, PDGFR, and VEGF-A), His-tagged binders were prepared at 5 µg/mL in PBSTA buffer (10 mM Na_2_HPO_4_·12H_2_O, 2 mM KH_2_PO_4_, 137 mM NaCl, 2.7 mM KCl, 0.05% Tween-20, 0.1% BSA, pH 7.4) for loading. Target proteins were diluted to different concentrations in PBSTA buffer (VEGF-A: 50 nM; IL-7R: 100 nM;

**Table S2:**
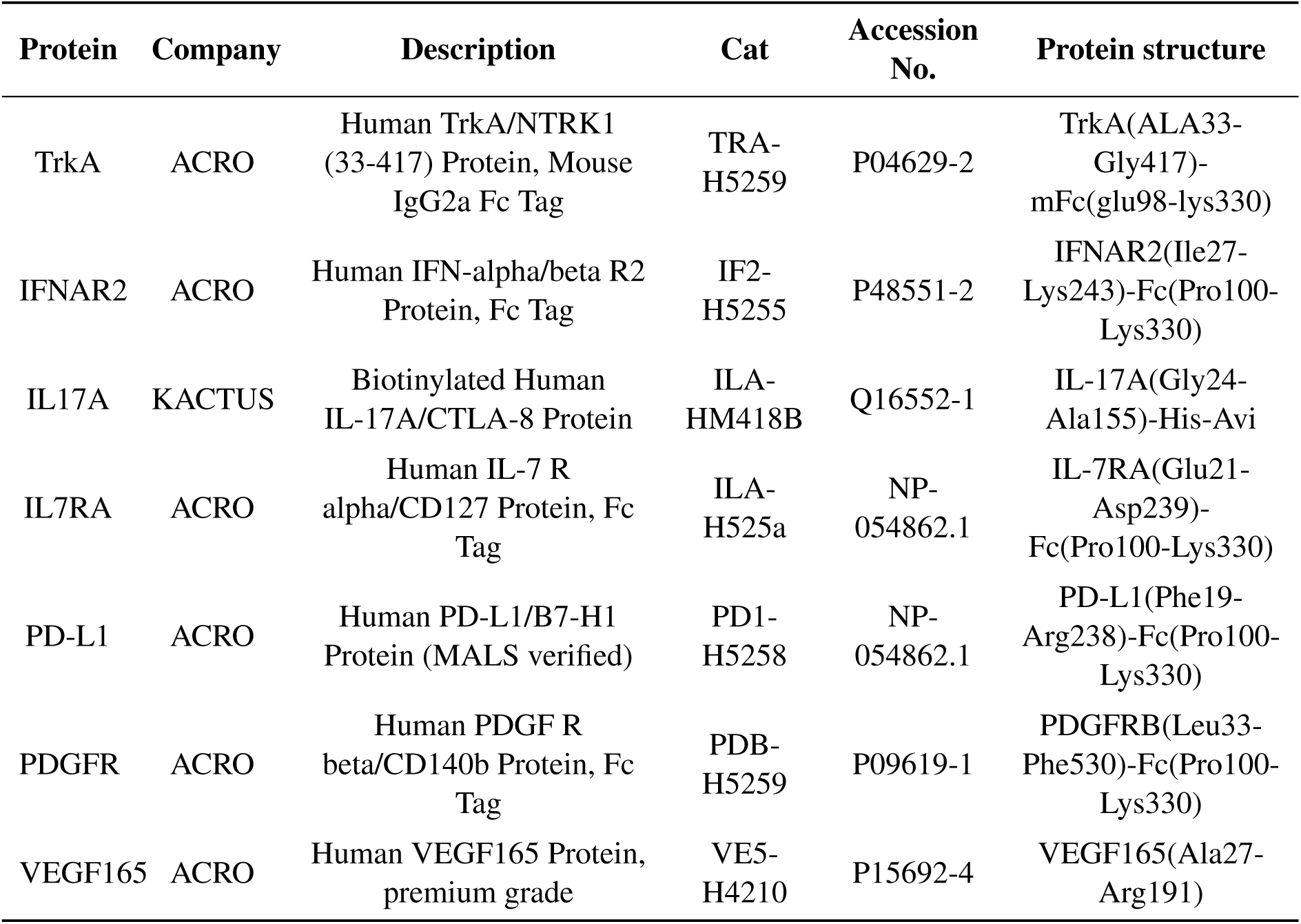
Binder design target protein.

IFNAR2: 200 nM; PDGFR: 400 nM; TrkA & PD-L1: 1000 nM each). Anti-His sensors (Gator Bio, Cat. No. 20-5066) were saturated with His-tagged binders. The BLI assay sequence was as follows: 60 s baseline, 120 s ligand loading, 60 s post-loading baseline, 120 s association phase, 300 s dissociation phase, and 30 s regeneration. Baseline-subtracted signals (in nanometers) were calculated to prioritize binders with specific binding to target proteins for further characterization. To determine the affinity of selected binder designs: His-tagged binders (5 µg/mL in PBSTA) were immobilized onto Anti-His sensors for 120 s. Serial dilutions of target proteins were prepared as follows: VEGF-A (50 nM to 3.125 nM), IL-7RA and IFNAR2 (400 nM to 6.25 nM), PDGFR (800 nM to 12.5 nM), and TrkA & PD-L1 (1000 nM to 15.625 nM). Diluted target proteins were allowed to associate with the immobilized ligand for 120 s, followed by dissociation in PBSTA buffer for 300 s. After background subtraction using buffer-only (PBSTA) control curves, dissociation constants (*K_d_*) were determined using the 1:1 binding model in the Results & Analysis curve-fitting module. For His-tagged target protein IL-17A: Biotin-labeled IL-17A was prepared at 5 *µ*g/mL in PBSTA. The design binders were diluted to 1000 nM in PBSTA. Streptavidin (SA) sensors (Gator Bio, 20-5006) were saturated with Biotin-labeled IL-17A for 60 s. The design binders were then allowed to associate with the immobilized ligand for 120 s, followed by a dissociation step in PBSTA for 300 s. The baseline subtracted signals (in nanometers) were calculated and used to prioritize target proteins specific binders for further characterization. To determine the affinity of selected designs, 5 *µ*g/mL Biotin-labeled IL-17A prepared in PBSTA was immobilized onto a Streptavidin (SA) sensor for 60 s. Serial dilutions of the design binders (1000 nM to 15.625 nM) were allowed to associate with the immobilized ligand for 120 s, followed by a dissociation step in PBSTA for 300 s. After background subtraction using buffer-only (PBSTA) control curves, *K_d_* values were determined using the 1:1 binding model in the Results & Analysis curve-fitting module.

### C.2 VHH experimental methods

#### C.2.1 Framework selection

VHH frameworks are randomly selected from 7EOW (caplacizumab), 8Z8V (ozoralizumab), 5JDS (en-vafolimab), 7XL0 (vobarilizumab), and 8COH (gefurulimab), highlighting their diverse structural architectures and clinical feasibility. For scFv design, frameworks are randomly selected from 6ZQK (trastuzumab), 6NOU (ixekizumab), 6TCS (omalizumab).

#### C.2.2 Gene Synthesis and Subcloning

VHH DNA sequences were codon-optimized and synthesized by GenScript Biotechnology Co., Ltd. (Nanjing, China). The synthesized genes were subcloned into the pCDNA3.4 vector for protein expression in CHO cell system. Subcloning into the pCDNA3.4 vector results in the following product: MGWSCIILFLVATATGVHS (signal peptide)-[Nanobody]-<u>GGGGSHHHHHH</u>, with the 6×His affinity tag underlined.

#### C.2.3 VHHs Expression in CHO Cells

Transfection-grade plasmids were prepared in large quantities using a GenScript self-developed plasmid extraction kit for CHO cell expression. Cells were maintained at 37^◦^C with 5% CO_2_ on an orbital shaker. One day before transfection, the cells were seeded at an appropriate density. On the day of transfection, DNA and Reagent (self-developed by GenScript) were mixed at an optimal ratio and then added into cells ready for transfection. Approximately 24 hours post transfection, Feed was added to each sample.

#### C.2.4 Protein Purification and Analysis

Cell culture broth was centrifuged and followed by filtration. Filtered cell culture supernatant was loaded onto an affinity purification column at an appropriate flowrate. After washing and elution with appropriate buffers, the eluted fractions were pooled and buffer exchanged to the final formulation buffer (PBS, pH 7.2). The purified protein was analyzed by SDS-PAGE and SEC-HPLC analysis to determine the molecular weight and purity. The concentration was determined by A280 method.

#### C.2.5 Target protein preparation

Since the VHHs designed in this study were expressed with a His-tag, target proteins were preferentially selected without a His-tag to avoid potential interference during affinity assessment. Most of these target proteins were obtained with an Fc-tag or in untagged forms. For CCL2 and BHRF1, commercially available proteins lacking alternative tags were unavailable. Therefore, BHRF1 was expressed with a His-SUMO tag in HEK293 cells, followed by tag cleavage. Biotin-labeled, His-tagged CCL2 was purchased. In such cases, affinity measurements were performed using immobilized antigens to evaluate binder binding.

#### C.2.6 Binding characterization of VHHs

Initial screening of VHH designs for target protein binding was performed using BLI (Gator Bio).

For target proteins without His-tag (PDGFR, 1433E, IL-13, BHRF1, S100A4 and HNMT), His-tagged VHHs were prepared at 5 *µ*g/mL in PBSTA buffer. Target proteins were diluted to different concentrations in PBSTA buffer (PDGFR: 5000 nM; IL-13, 1433E, BHRF1, S100A4 and HNMT: 1000 nM each). Anti-His sensors (Gator Bio, Cat. No. 20-5066) were saturated with His-tagged VHHs. The BLI assay sequence was as follows: 60 s baseline, 120 s ligand loading, 60 s post-loading baseline, 120 s association phase, 300 s dissociation phase, and 30 s regeneration. Baseline-subtracted signals (in nanometers) were calculated to prioritize VHHs with specific binding to target proteins for further characterization. For EFNA1(1000 nM), the signal value detected by the HIS1K sensor was too low. Therefore, the ProA sensor was used for detection, and the detection conditions were the same as those of the HIS1K sensor.

**Table S3:**
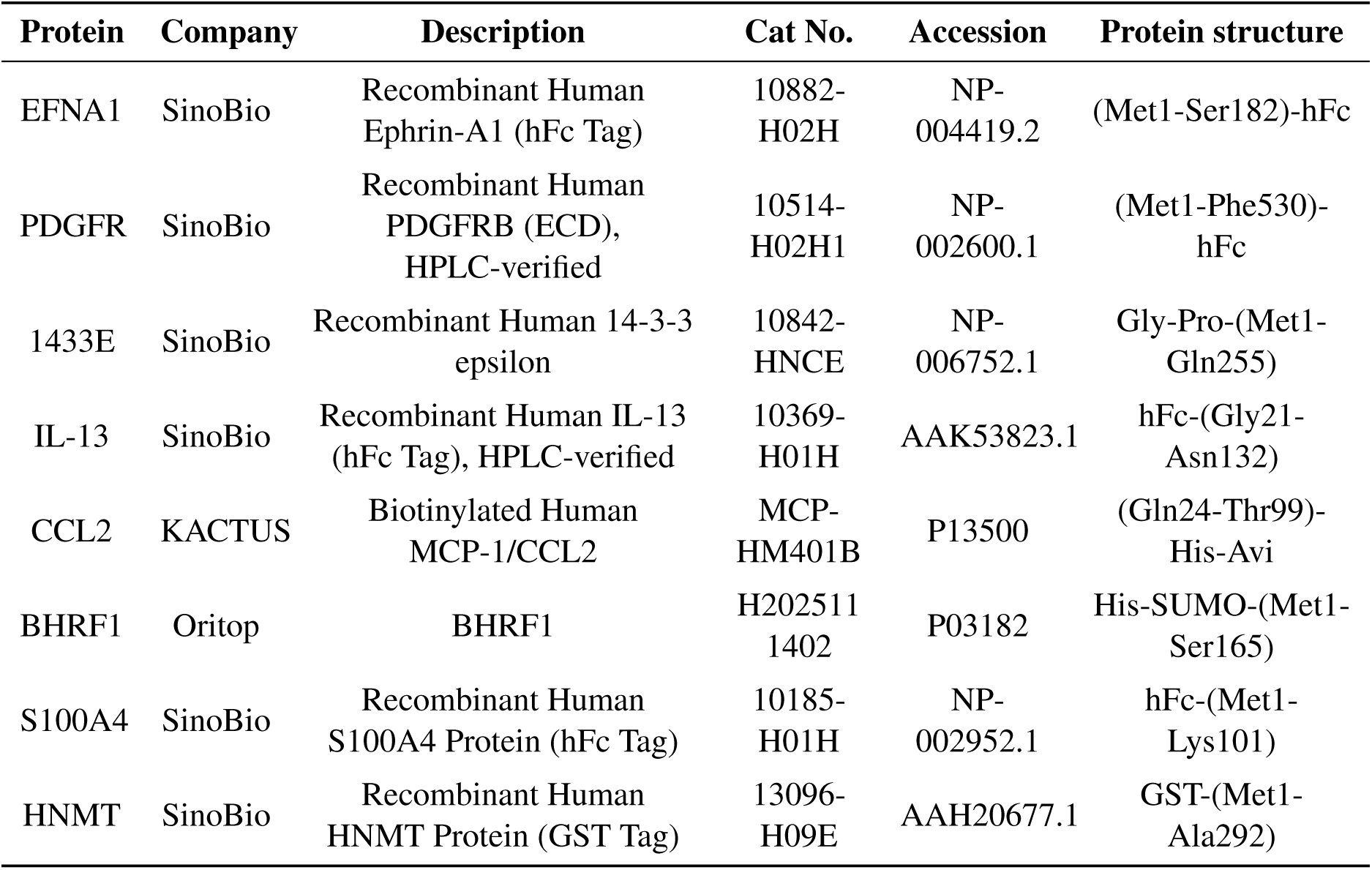
VHH design target protein.

To determine the affinity of selected designs, 5 *µ*g/mL His tag design VHHs prepared in PBSTA were immobilized onto Anti-His sensor for 120 s. Serial dilutions of the target proteins (PDGFR, IL-13 and BHRF1: from 5000 nM diluted to 78 nM, 1433E: from 3600 nM diluted to 56.25 nM, S100A4 and HNMT from 4000 nM diluted to 125 nM,) were then allowed to associate with the immobilized ligand for 120 s, followed by a dissociation step in PBSTA for 300 s. After background subtraction using buffer-only (PBSTA) control curves, *K_d_* values were determined using the 1:1 binding model in the Results & Analysis curve-fitting module.For EFNA1(from 4000 nM diluted to 125 nM), the ProA sensor was used for detection, and the detection conditions were the same as those of the HIS1K sensor. For His tag target protein CCL2: Biotin-labeled CCL2 was prepared at 5 *µ*g/mL in PBSTA. VHHs were diluted to 5000 nM in PBSTA. Streptavidin (SA) sensors (Gator Bio, 20-5006) were saturated with biotin-labeled CCL2 for 60 s. VHHs were then allowed to associate with the immobilized ligand for 120 s, followed by a dissociation step in PBSTA for 300 s. Baseline-subtracted signals (in nanometers) were calculated to prioritize specific VHHs for further characterization.

To determine the affinity of selected VHHs for CCL2: Biotin-labeled CCL2 (5 *µ*g/mL in PBSTA) was immobilized onto SA sensors for 60 s. Serial dilutions of VHHs (5000 nM to 78 nM) were allowed to associate with the immobilized ligand for 120 s, followed by dissociation in PBSTA buffer for 300s. After background subtraction using buffer-only (PBSTA) control curves, *K_d_* values were determined using the 1:1 binding model in the Results & Analysis curve-fitting module.

## Appendix D Supplementary Figures

**Figure S1:**
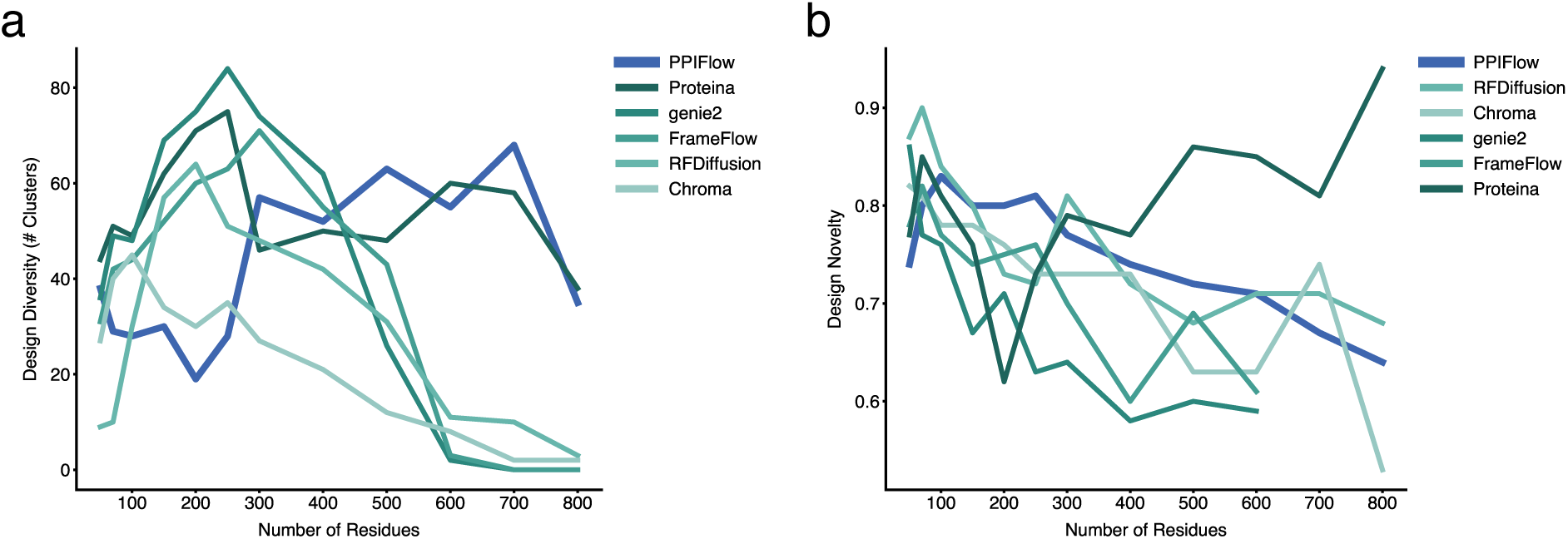
Benchmarking monomer generation performance across varying lengths. **(a)** Evaluation of structural diversity in monomer generation tasks for sequences ranging from 50 to 800 amino acids (100 designs per length). PPIFlow is compared against Proteina, Genie2, FrameFlow, RFDiffusion, and Chroma. **(b)** Structural novelty analysis quantified by the maximum TM-score of successful designs against the PDB databases.

**Figure S2:**
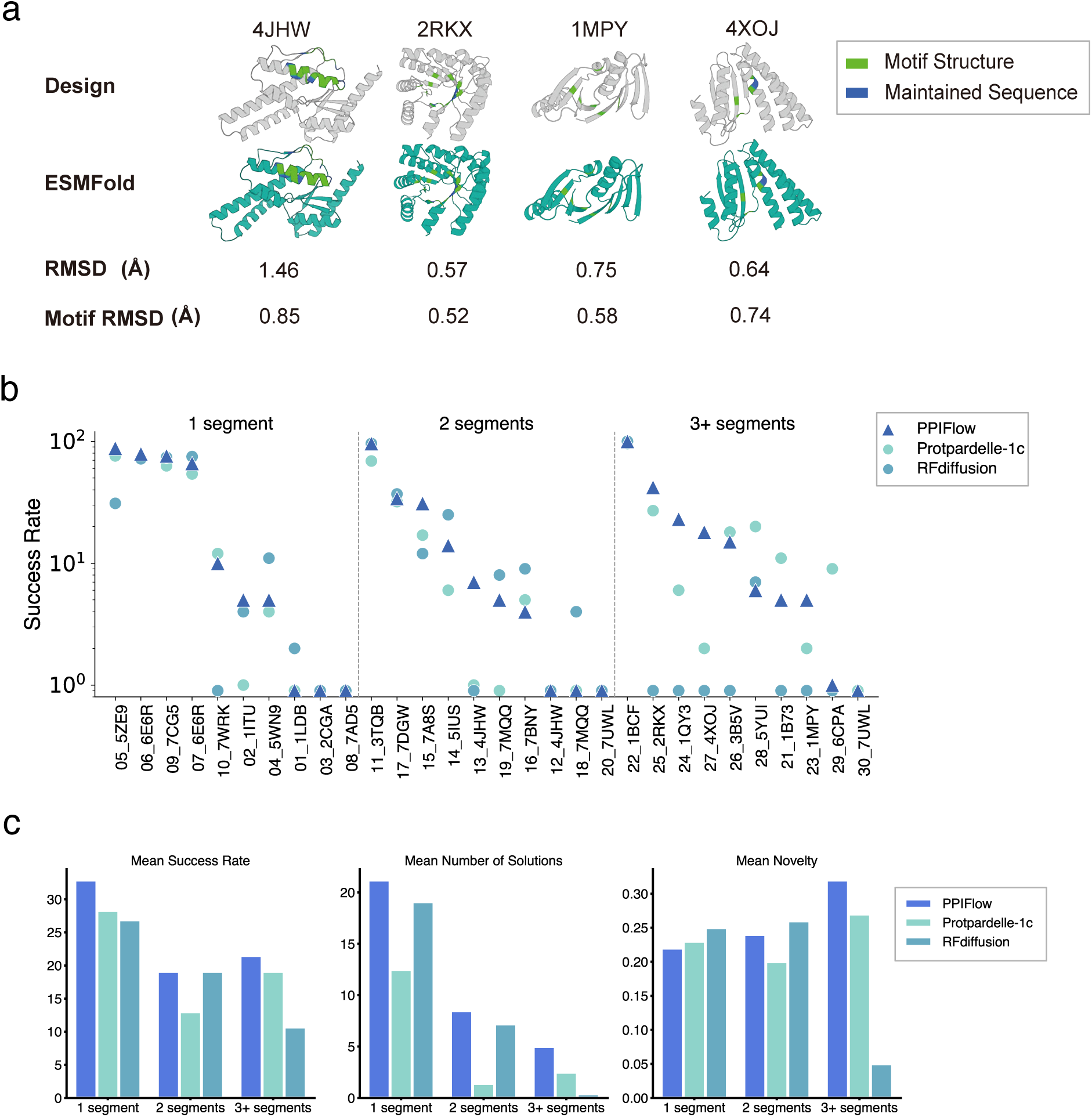
Assessment of motif scaffolding performance and structural fidelity. **(a)** Performance of PPIFlow on four representative scaffolding tasks (PDB IDs: 4JHW, 2RKX, 1MPY, and 4XOJ). Panels show the alignment between PPIFlow-generated backbones and ESMFold-refolded structures, annotated with global C*α* RMSD and motif-specific C*α* RMSD values. **(b)** Comparison of design success rates on the MotifBench dataset between PPIFlow, Protpardelle-1c, and RFDiffusion. **(c)** Comprehensive comparison of the three models across mean success rate, solution diversity, and structural novelty.

**Figure S3:**
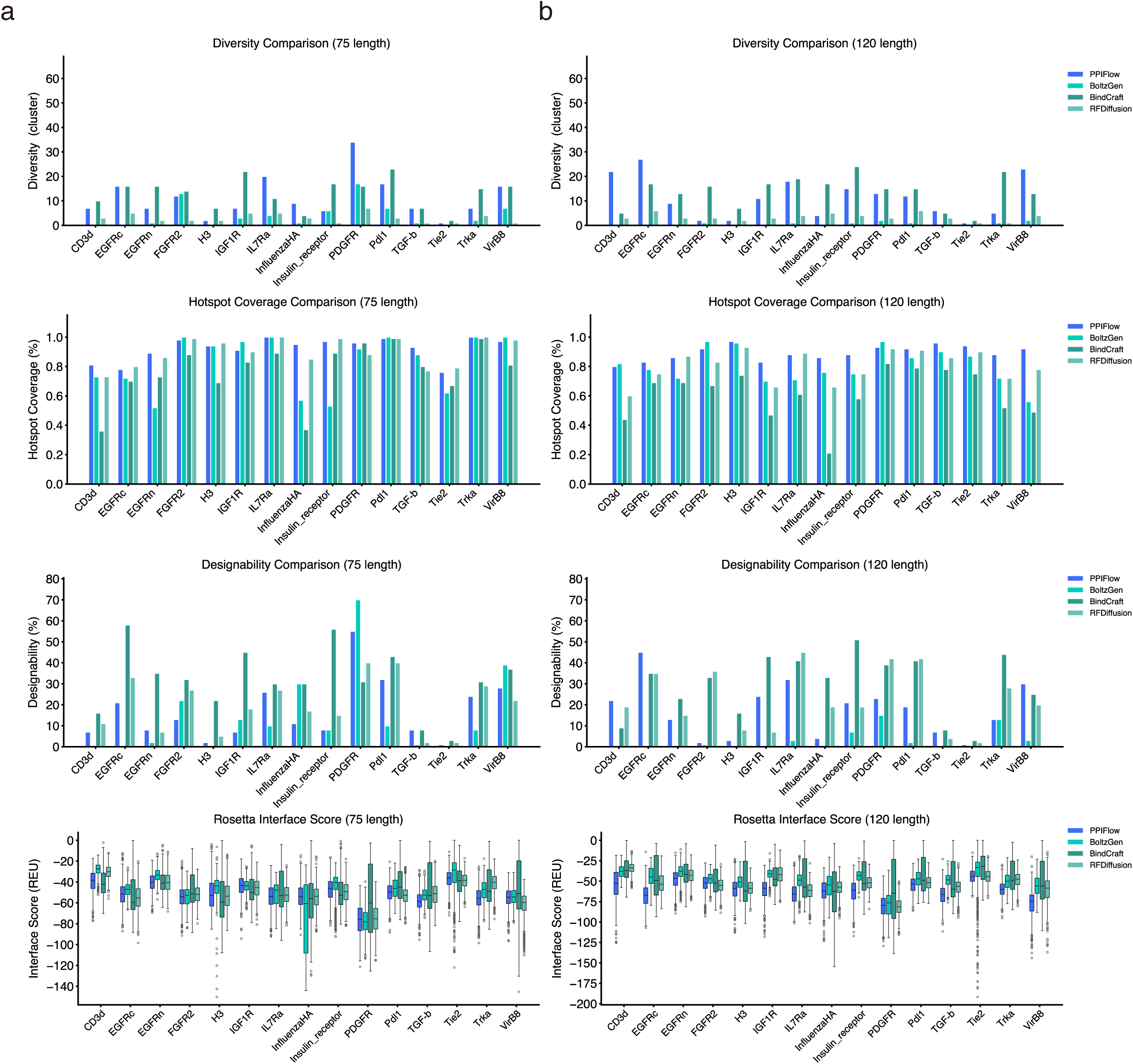
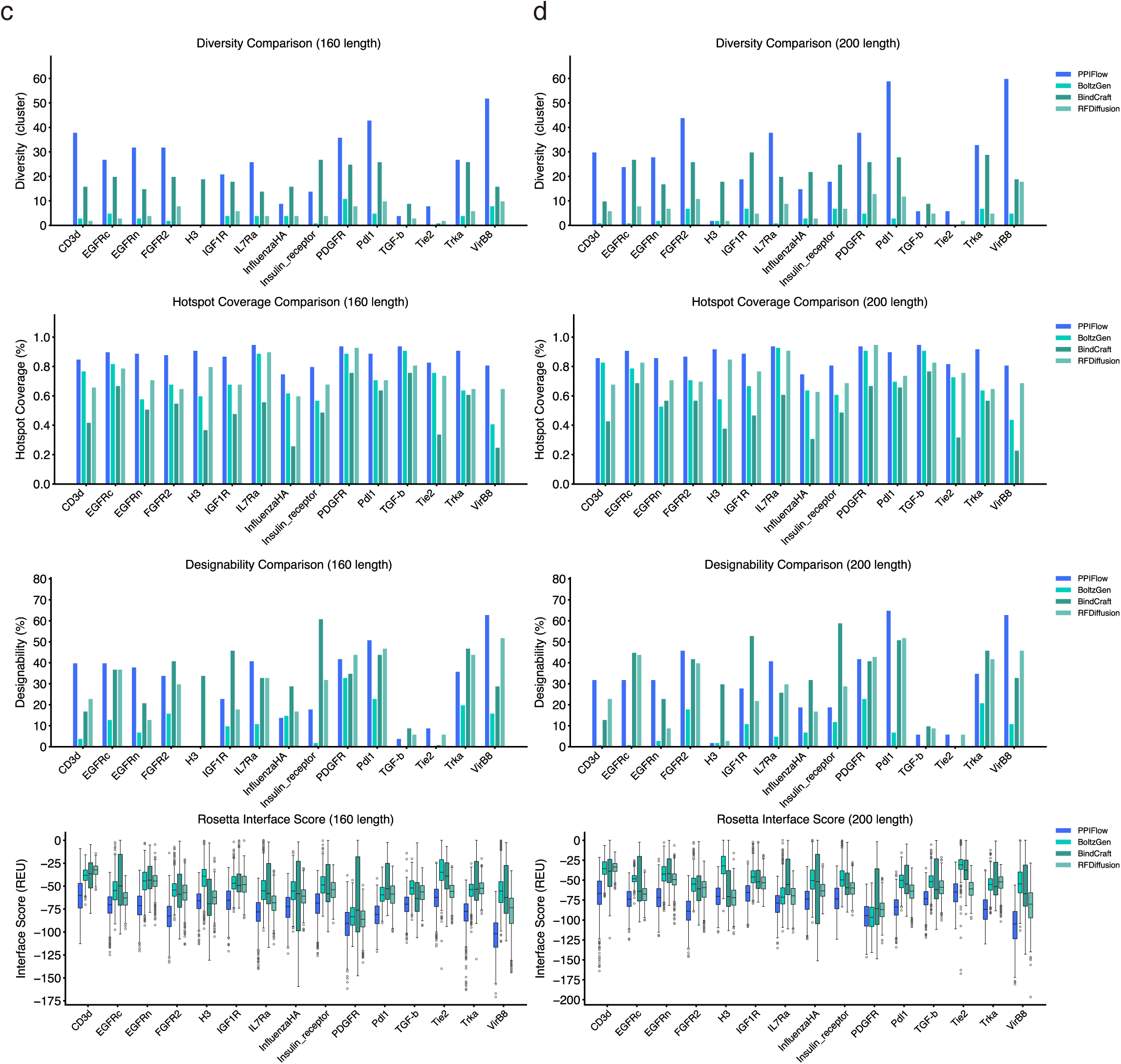
Comparative analysis of de novo binder design efficiency. Results are grouped by target sequence length: **(a)** 75 residues, **(b)** 120 residues, **(c)** 160 residues, and **(d)** 200 residues. For each group, four hierarchical performance metrics are evaluated (from top to bottom): *(i) Structural diversity*, quantifying the conformational breadth of the generated binder ensemble; *(ii) Hotspot coverage*, measuring the spatial conservation of critical target-interface residues; *(iii) Designability*, defined by the structural self-consistency (scRMSD) between generative backbones and forward-folded structures; *(iv) Binding energetics*, represented by Rosetta interface scores (REU). All subfigures share a common color scheme representing the four evaluated algorithms.

**Figure S4:**
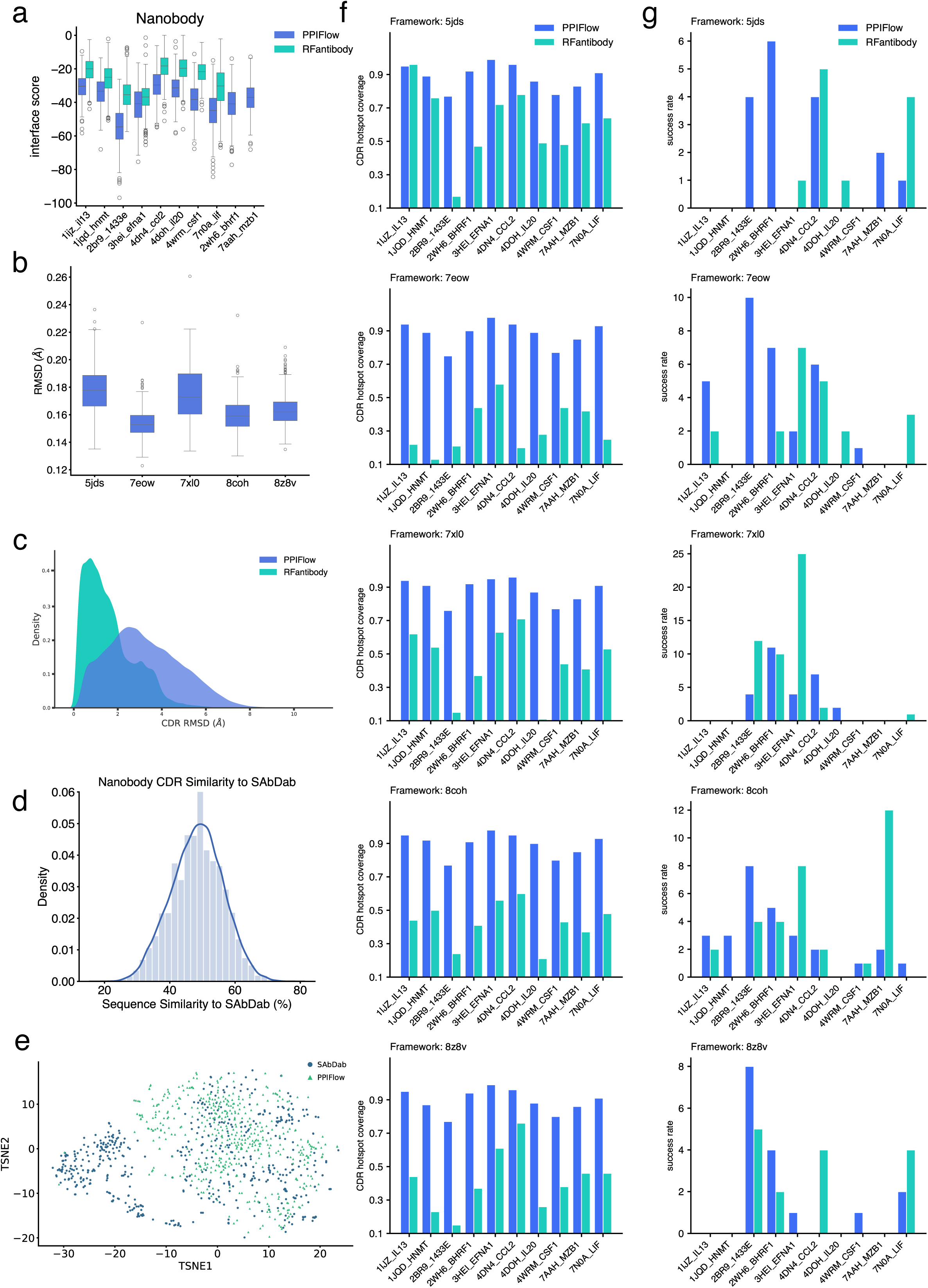
Benchmarking nanobody (VHH) design precision and diversity. **(a)** Distribution of Rosetta interface scores comparing PPIFlow and RFAntibody across nanobody design benchmarks. Notably, RFAntibody failed to generate valid binder candidates for the BHRF1 and MZB1 targets. **(b)** Structural recovery of framework regions (FRs), quantified by C*α* RMSD (Å) across multiple nanobody template PDB IDs, highlighting the backbone stability of the designs. **(c)** Conformational fidelity of CDR3 loops, calculated using the PyMOL alignment algorithm. Structures were pre-aligned via framework regions, followed by CDR3 RMSD calculation without further rotation to assess loop positioning accuracy. **(d, e)** Sequence and structural landscape analysis. (d) Sequence diversity of designed CDR3 loops relative to the SAbDab database (training set); (e) Structural clustering of the generated ensembles, illustrating the breadth of the sampled conformational space compared to known antibody repertoires. **(f, g)** Comprehensive evaluation of design quality across framework templates, reporting (f) CDR hotspot coverage efficiency and (g) design success rates across the diverse target panel.

**Figure S5:**
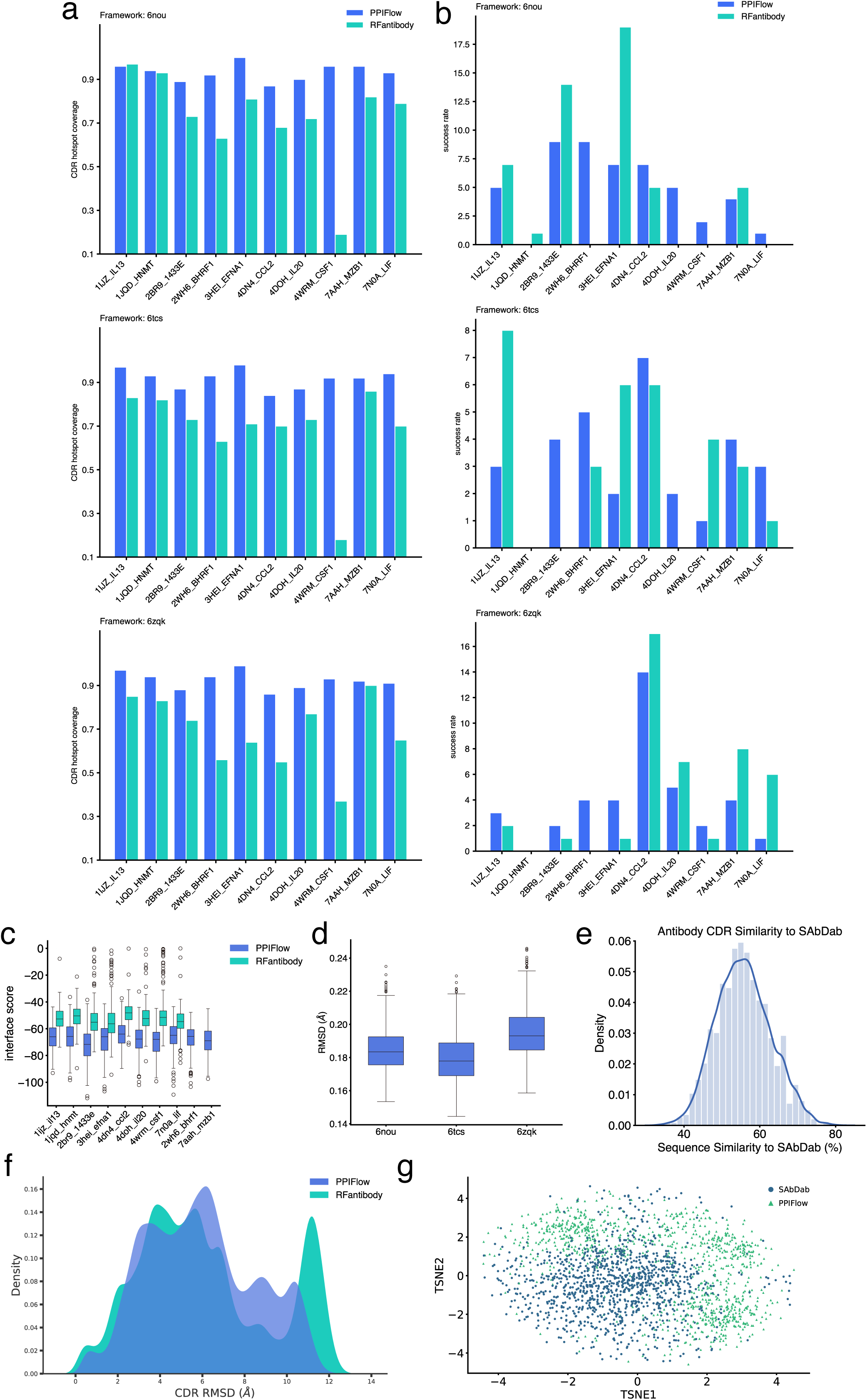
Benchmarking scFv design precision. (a, **b)** Detailed performance metrics for antibody design tasks across a library of diverse framework templates. Panels illustrate the efficiency of PPIFlow in maintaining CDR hotspot coverage and achieving high success rates for various antibody frameworks; **(c)** Distribution of Rosetta interface scores comparing PPIFlow and RFAntibody across scFv design benchmarks. Notably, RFAntibody failed to generate valid binder candidates for the BHRF1 and MZB1 targets. **(d)** Structural recovery of framework regions (FRs), quantified by C*α* RMSD (Å) across multiple template PDB IDs, highlighting the backbone stability of the designs. **(e)** Sequence diversity of designed CDR3 loops relative to the SAbDab database (training set); **(f)** Conformational fidelity of CDR3 loops, calculated using the PyMOL alignment algorithm. Structures were pre-aligned via framework regions, followed by CDR3 RMSD calculation without further rotation to assess loop positioning accuracy. **(g)** Structural clustering of the generated ensembles, illustrating the breadth of the sampled conformational space compared to known antibody repertoires.

**Figure S6:**
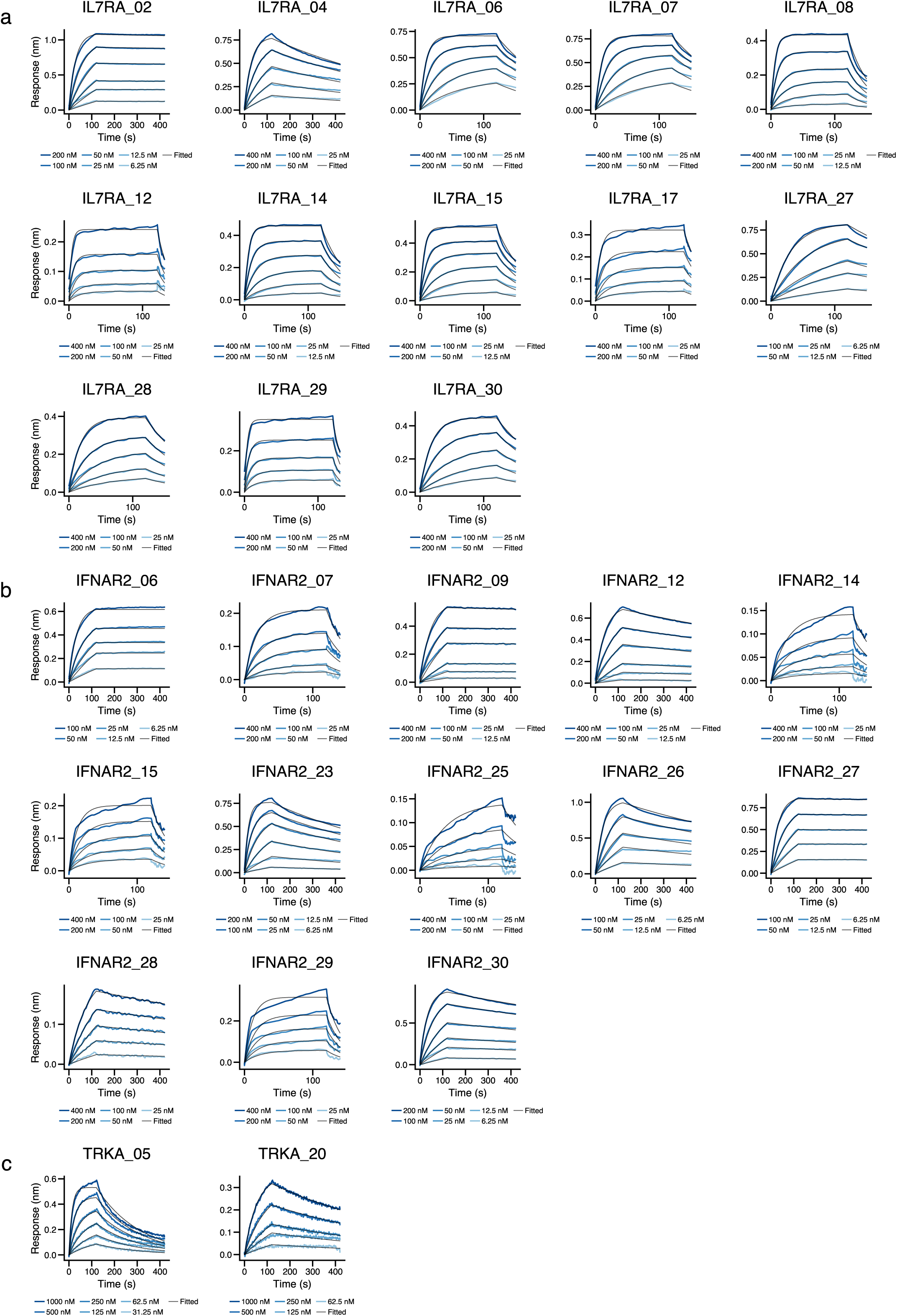
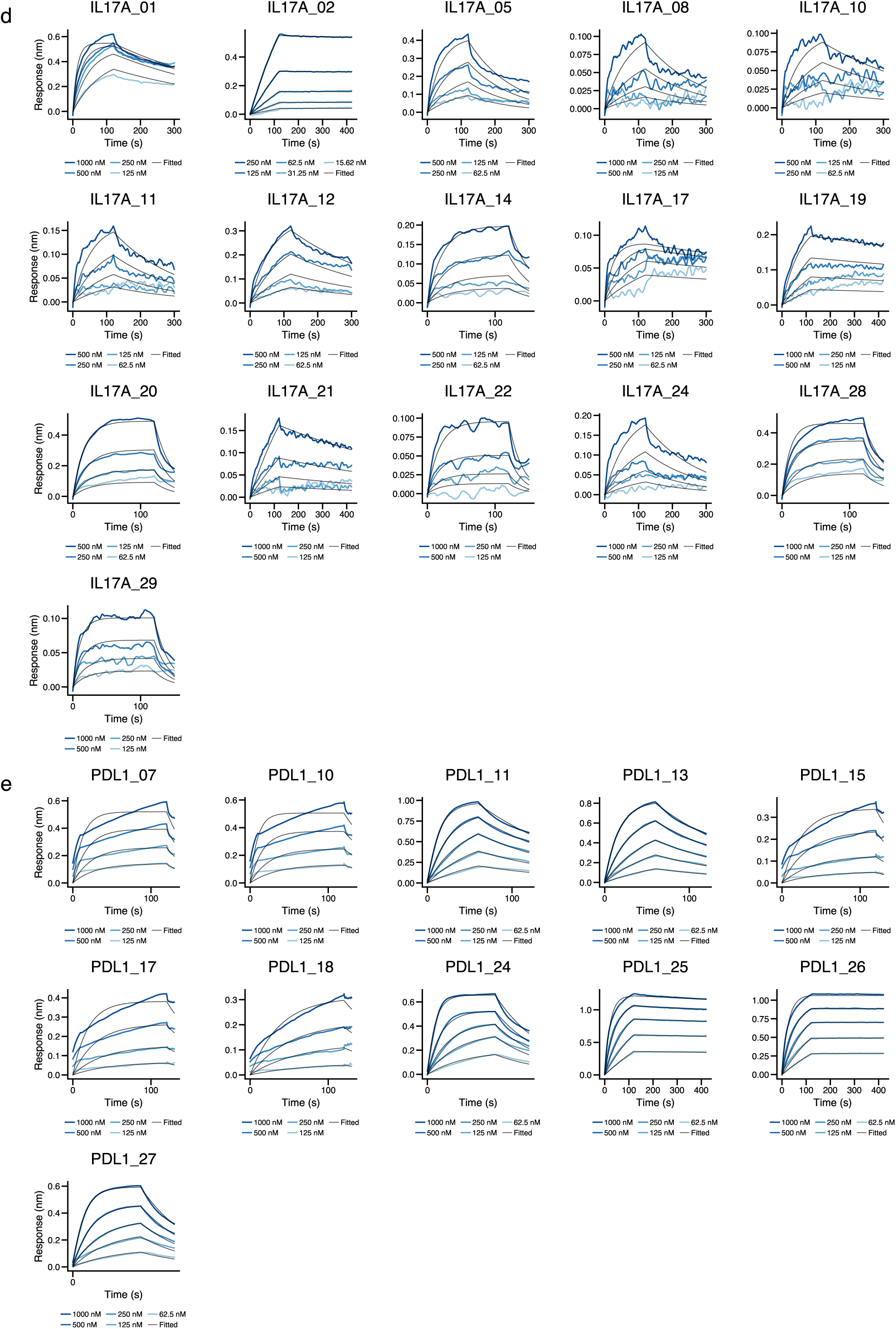
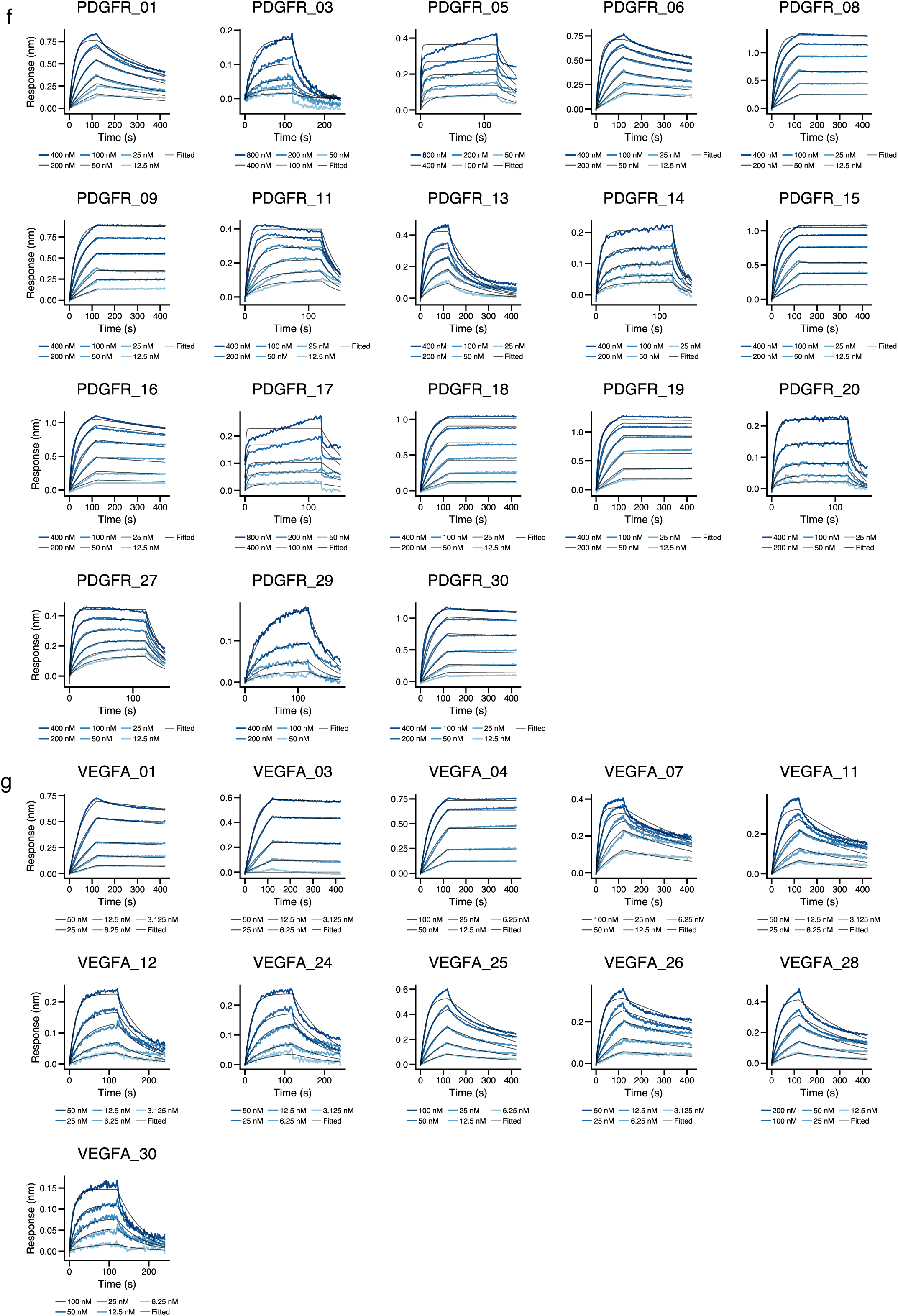
Bio-layer interferometry (BLI) kinetic analysis of de novo designed binders. Multi-cycle BLI sensorgrams illustrating the binding kinetics of synthetic binders against seven therapeutic targets: **(a)** IL7RA, **(b)** IFNAR2, **(c)** TrkA, **(d)** IL17A, **(e)** PDL1, **(f)** PDGFR, and **(g)** VEGFA. Raw data (colored lines) were globally fitted with a 1:1 binding model (black lines). Molar concentrations for each titration step are annotated below the respective sensorgrams.

**Figure S7:**
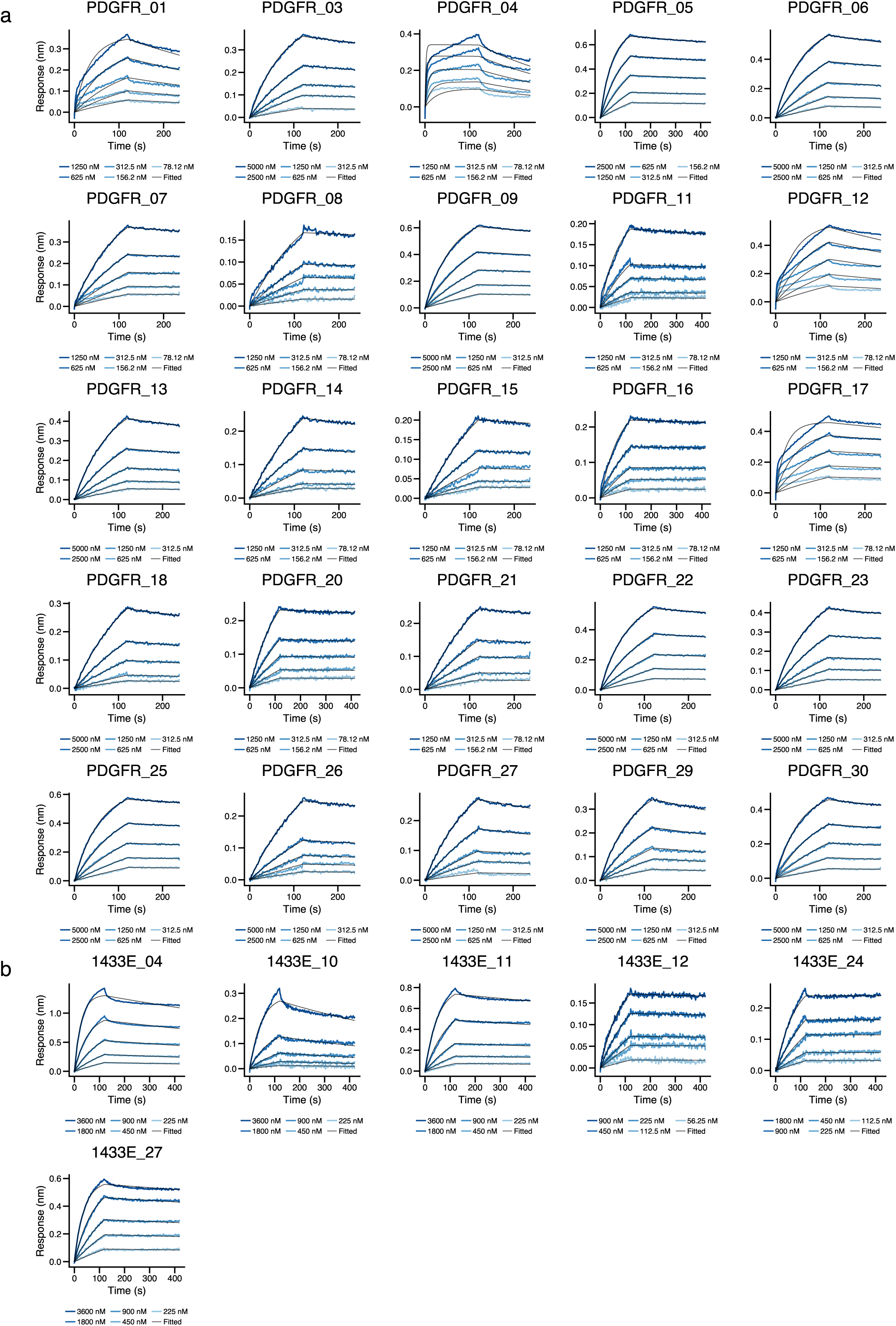
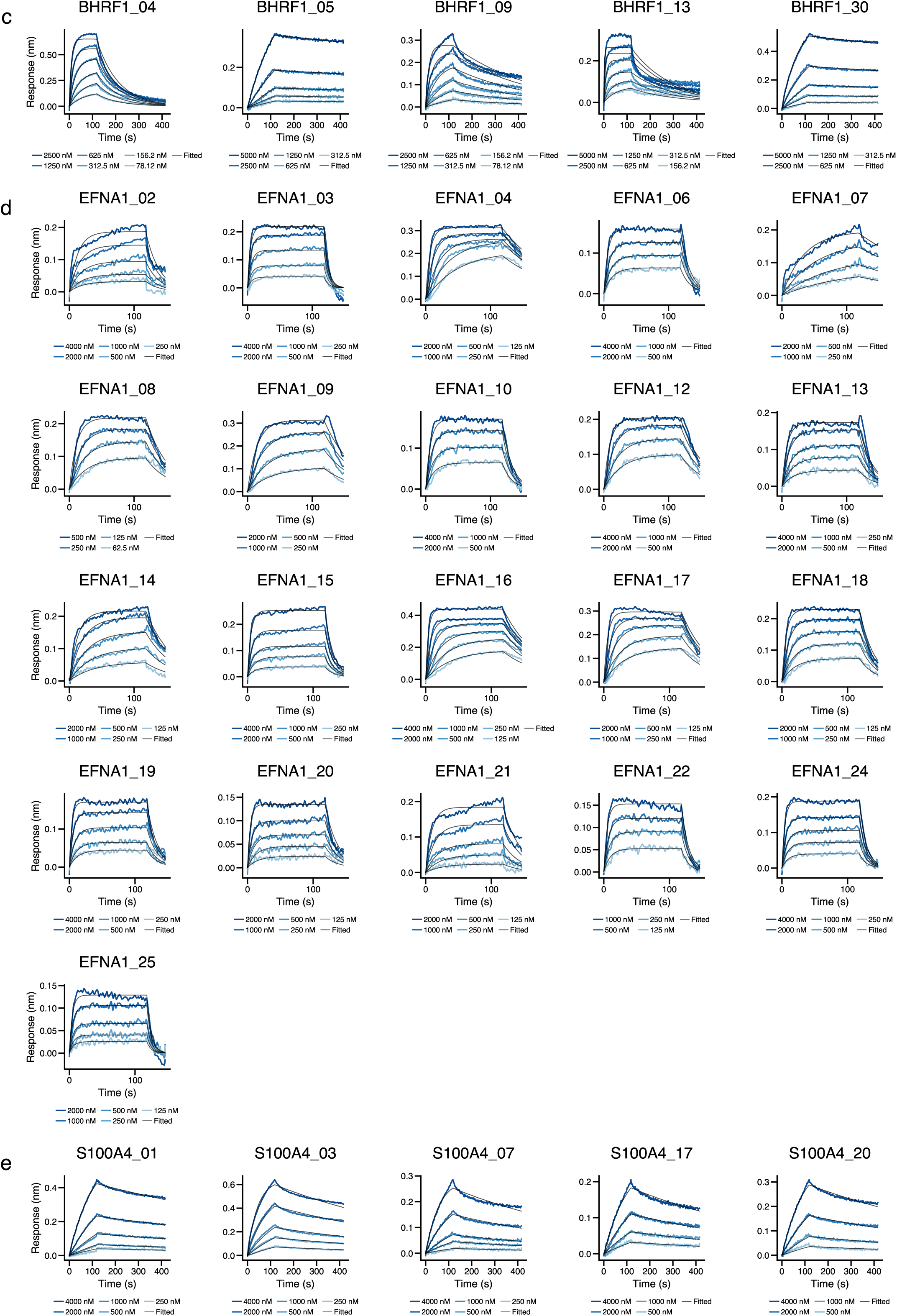
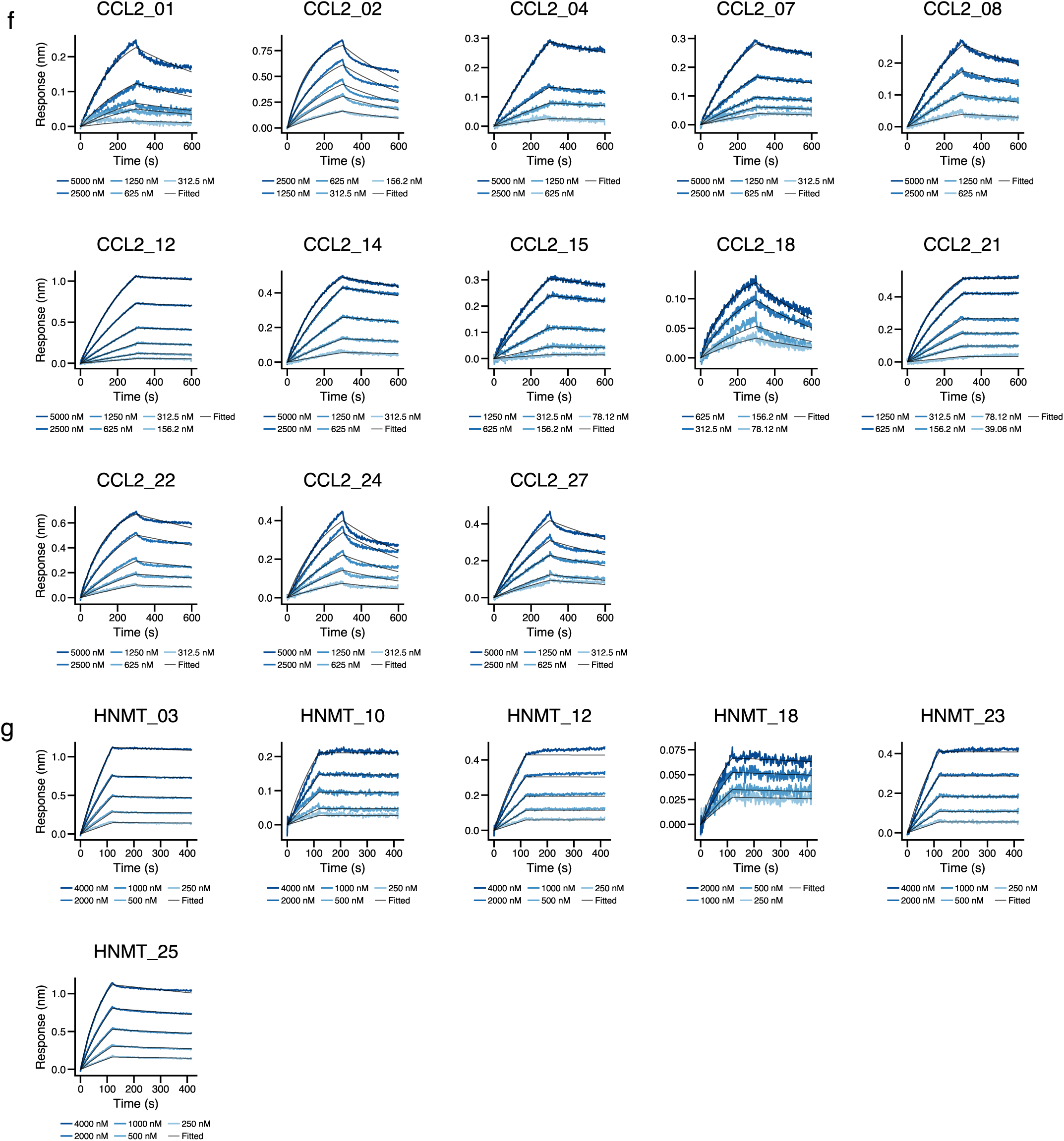
Kinetic characterization of designed VHH domains via BLI. Representative BLI sensorgrams showing the binding affinity of designed VHH candidates against: **(a)** PDGFR, **(b)** 1433E, **(c)** BHRF1, **(d)** EFNA1, **(e)** S100A4, **(f)** CCL2 and **(g)** HNMT. Multi-point kinetic titrations were performed to evaluate binding specificity. The data confirm high-affinity interactions and diverse dissociation profiles across the tested VHH-target complexes.

**Figure S8:**
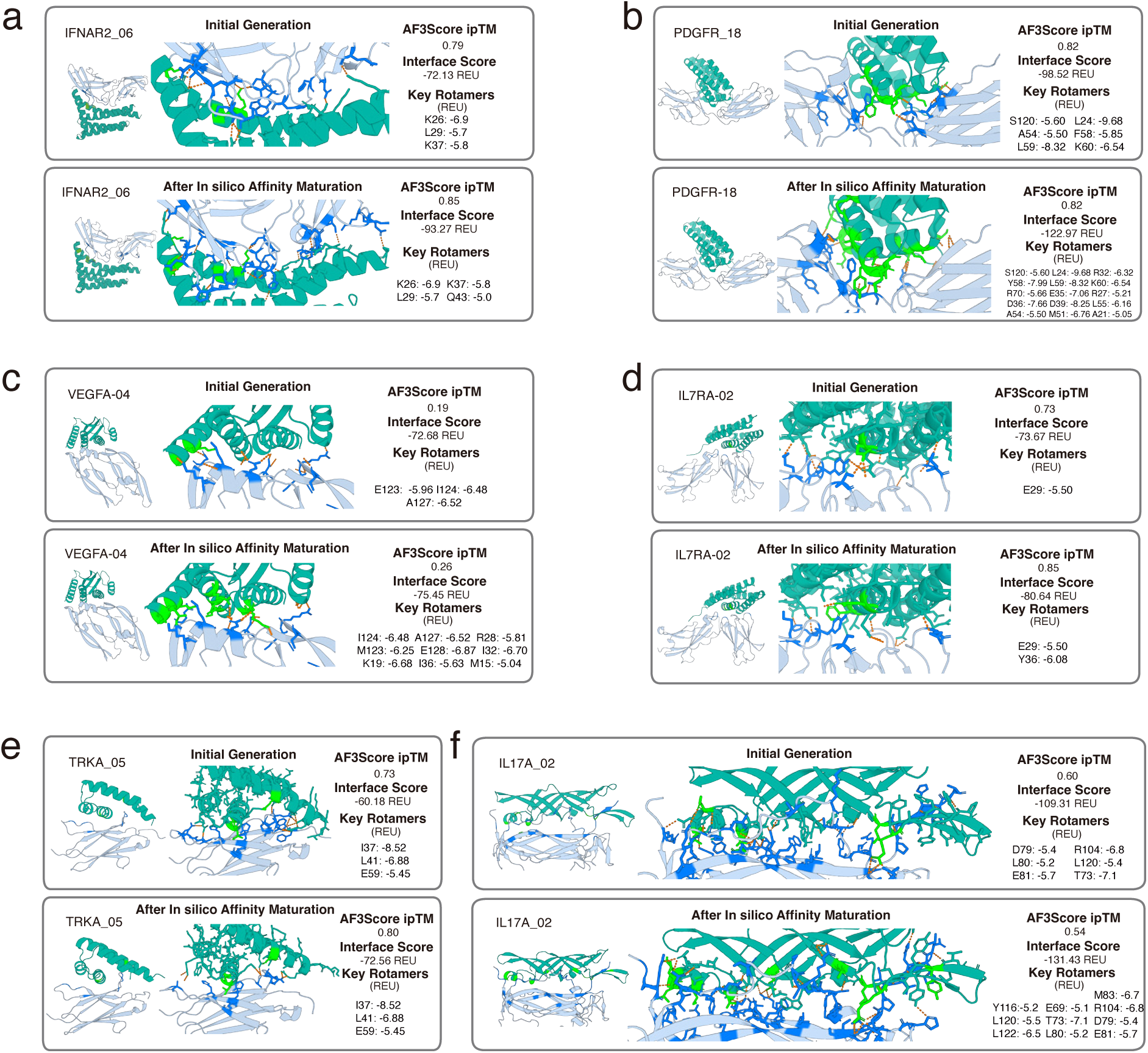
Affinity maturation and structural characterization of designed binders. (a–f) Optimization of lead candidates for each target. (Note: PD-L1 is omitted here as it is presented in the main figure). Each panel displays a comparative view between the initial generative model output (top) and the variant after affinity maturation (bottom). **Structural details:** Left panels illustrate the global fold and magnified interface interactions. Key residues on the designed binders are highlighted in *green*, with target-binding residues shown in *marine*. Hydrogen bonds are represented by *orange dashed lines*. **Computational metrics:** The corresponding AF3Score confidence scores and Rosetta interface energy scores are provided to the right of each structure. The interface decomposition highlights critical residues achieving total interaction energy *<* −5 Rosetta Energy Units (REU).

**Figure S9:**
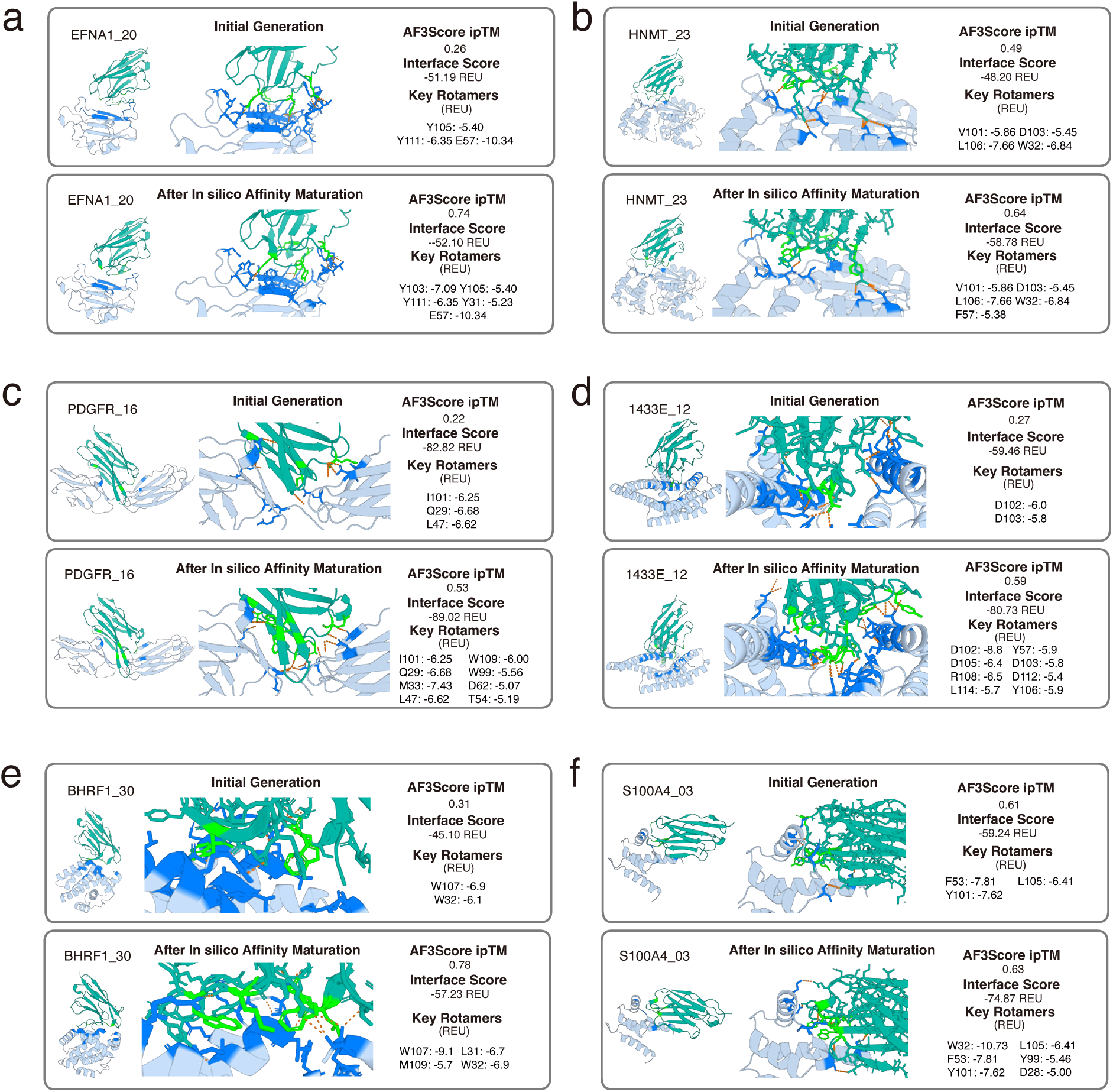
Affinity maturation and structural characterization of designed VHHs. (a–f) Optimization of lead candidates for each target. (Note: CCL2 is omitted here as it is presented in the main figure). Each panel displays a comparative view between the initial generative model output (top) and the variant after affinity maturation (bottom). **Structural details:** Left panels illustrate the global fold and magnified interface interactions. Key residues on the designed binders are highlighted in *green*, with target-binding residues shown in *marine*. Hydrogen bonds are represented by *orange dashed lines*. **Computational metrics:** The corresponding AF3Score confidence scores and Rosetta interface energy scores are provided to the right of each structure. The interface decomposition highlights critical residues achieving total interaction energy *<* −5 Rosetta Energy Units (REU).

## Appendix E Supplementary Tables

**Table S4:**
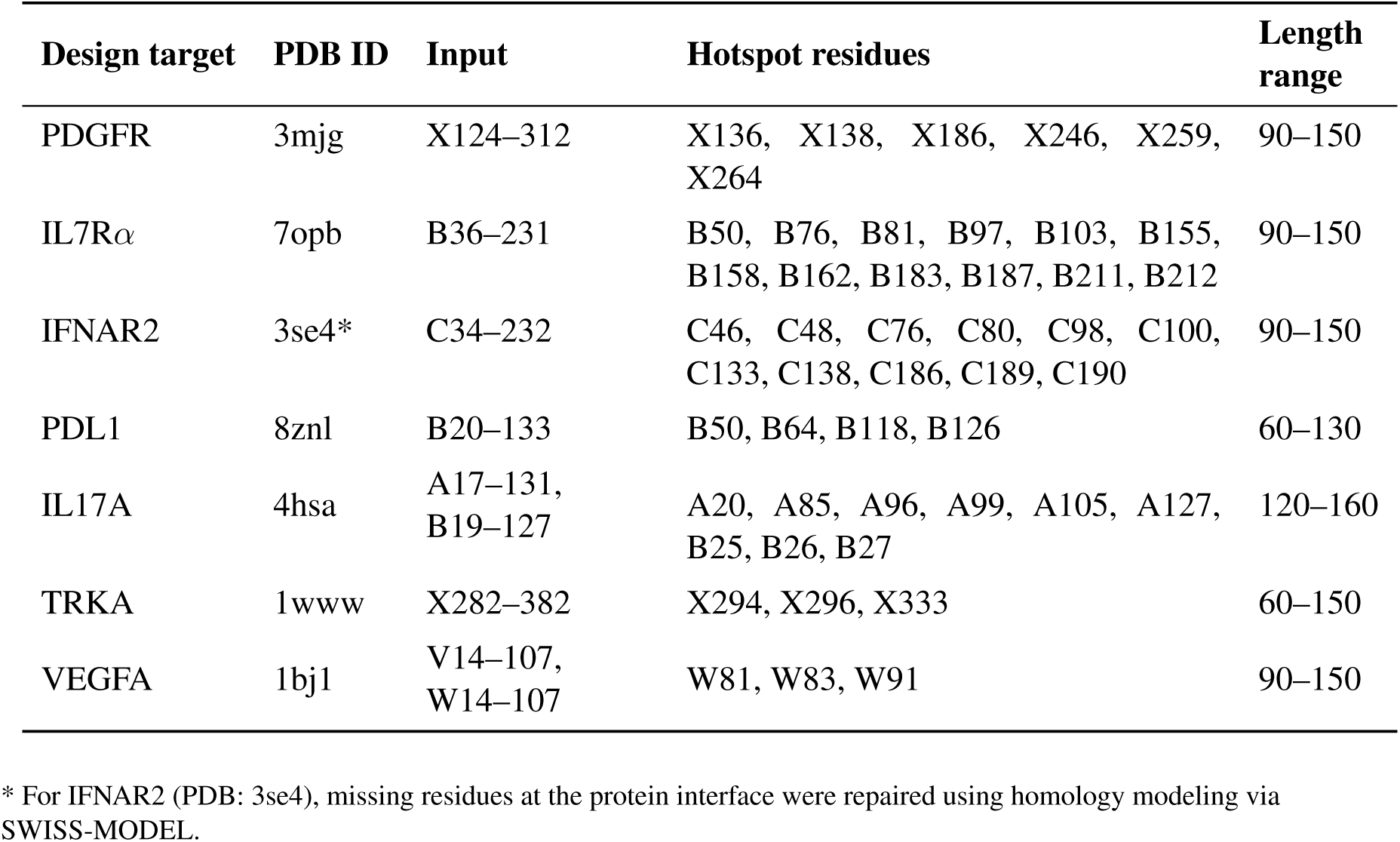
Binder design task specifications for *in silico* benchmarking and experimental testing.

**Table S5:**
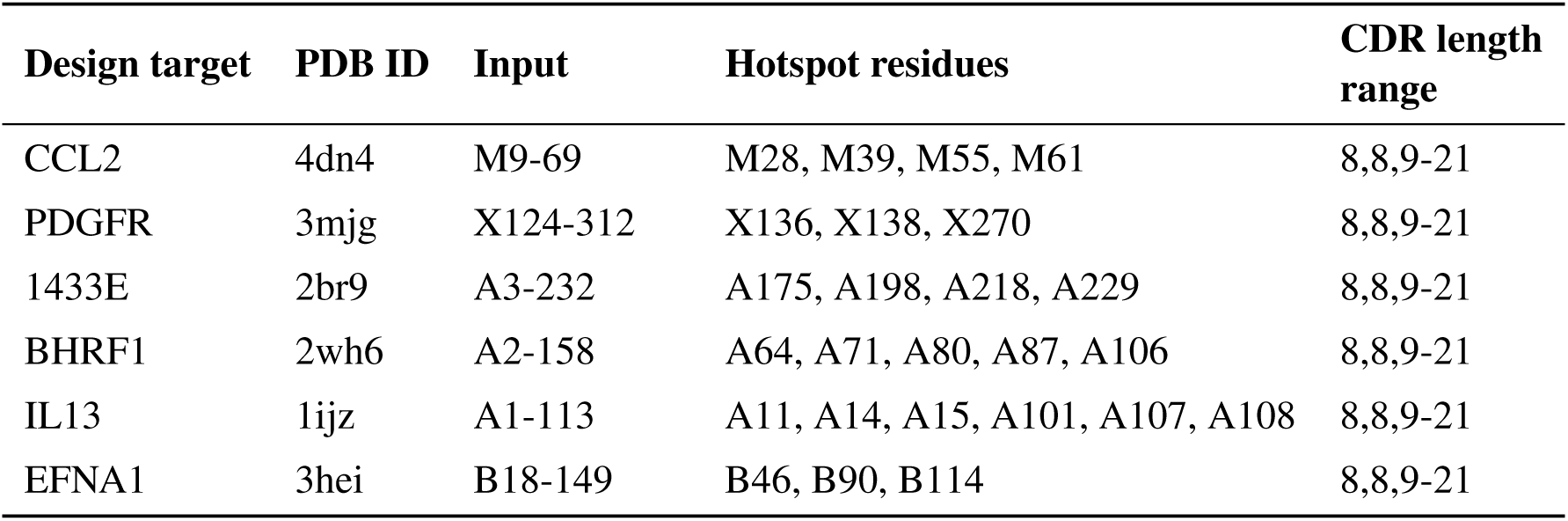
VHH design problem specifications for *in silico* benchmarking and experimental testing.

**Table S6:**
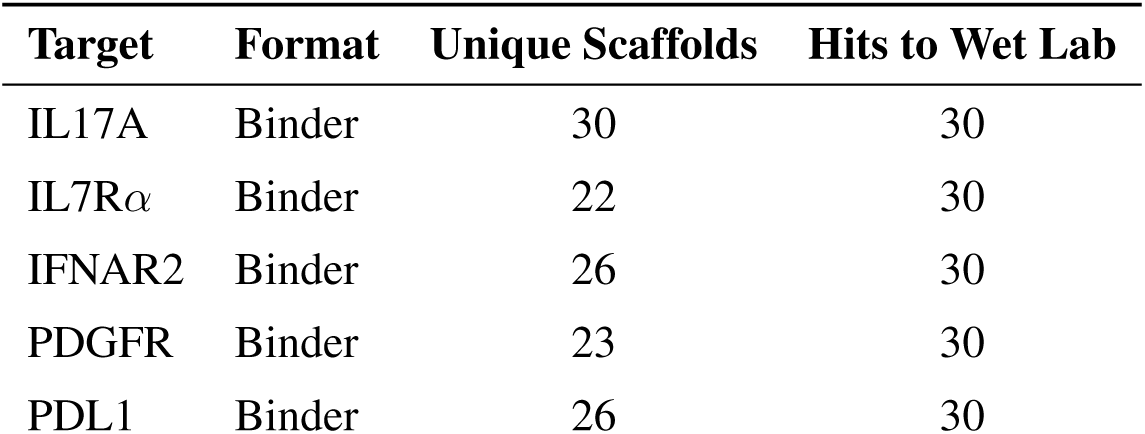
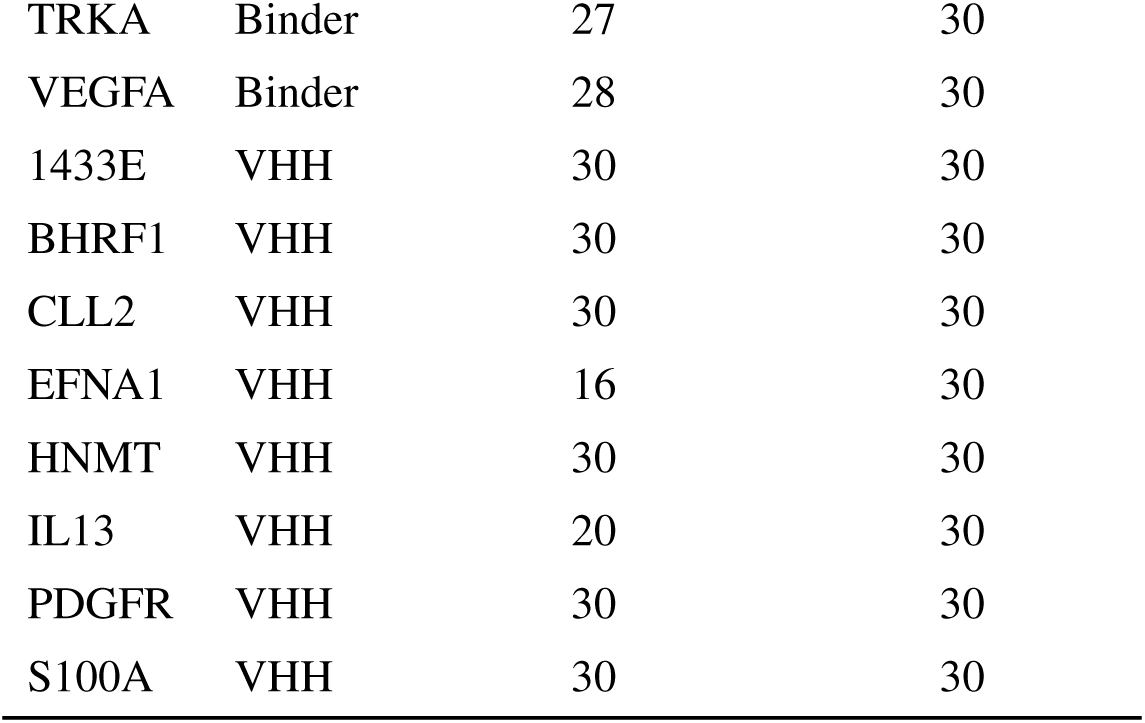
Design summary for protein and unique scaffolds information.

**Table S7:**
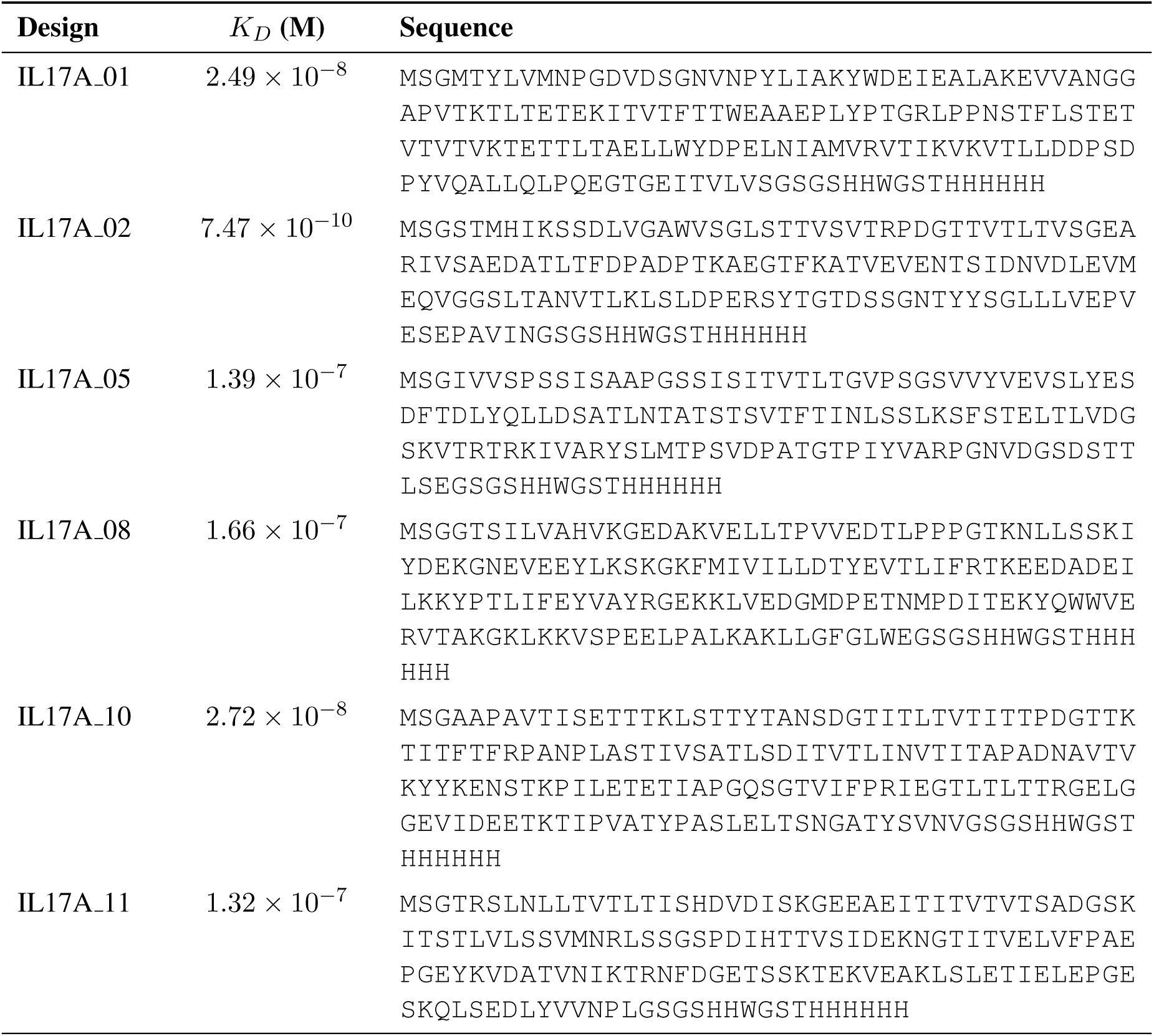
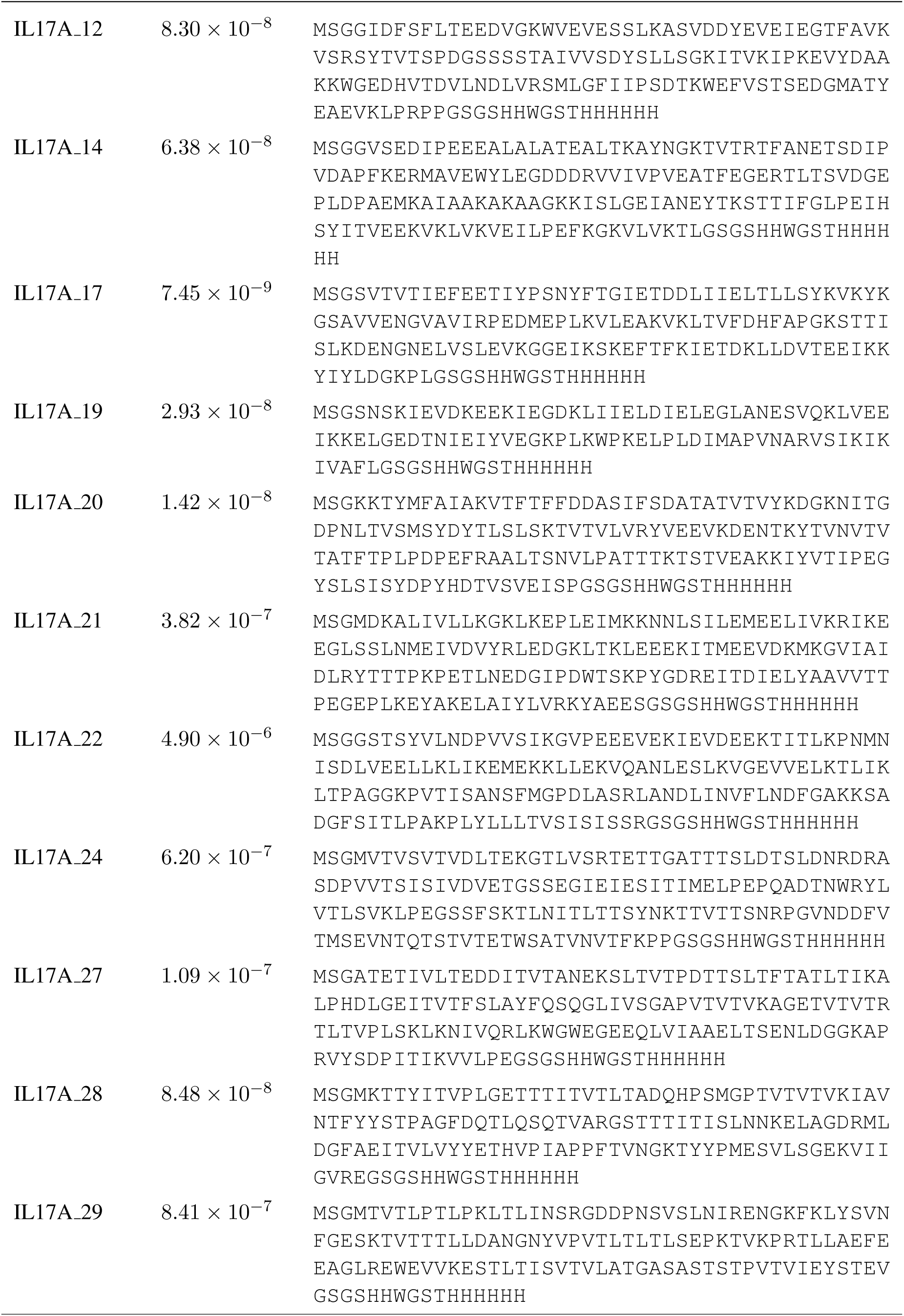
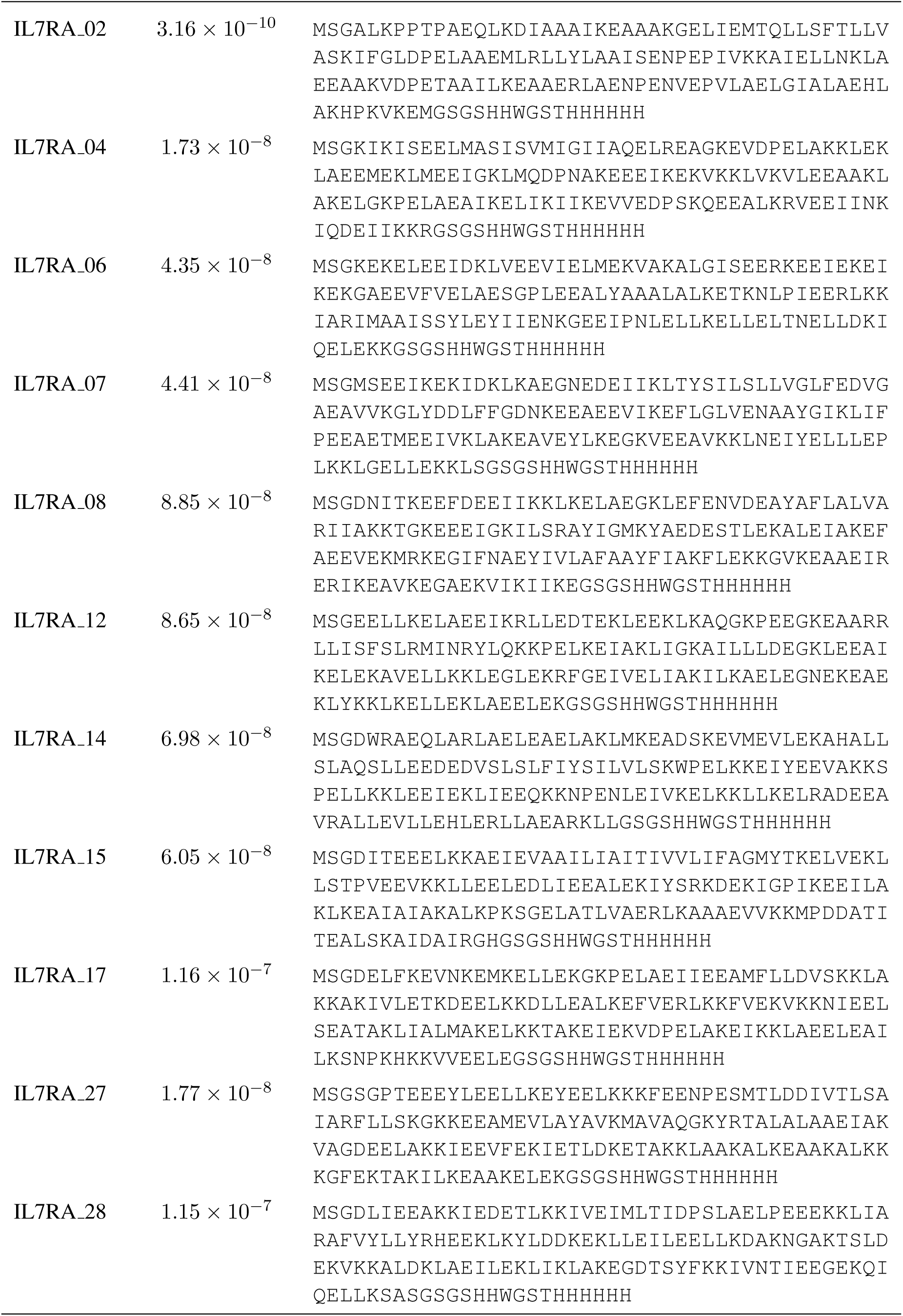
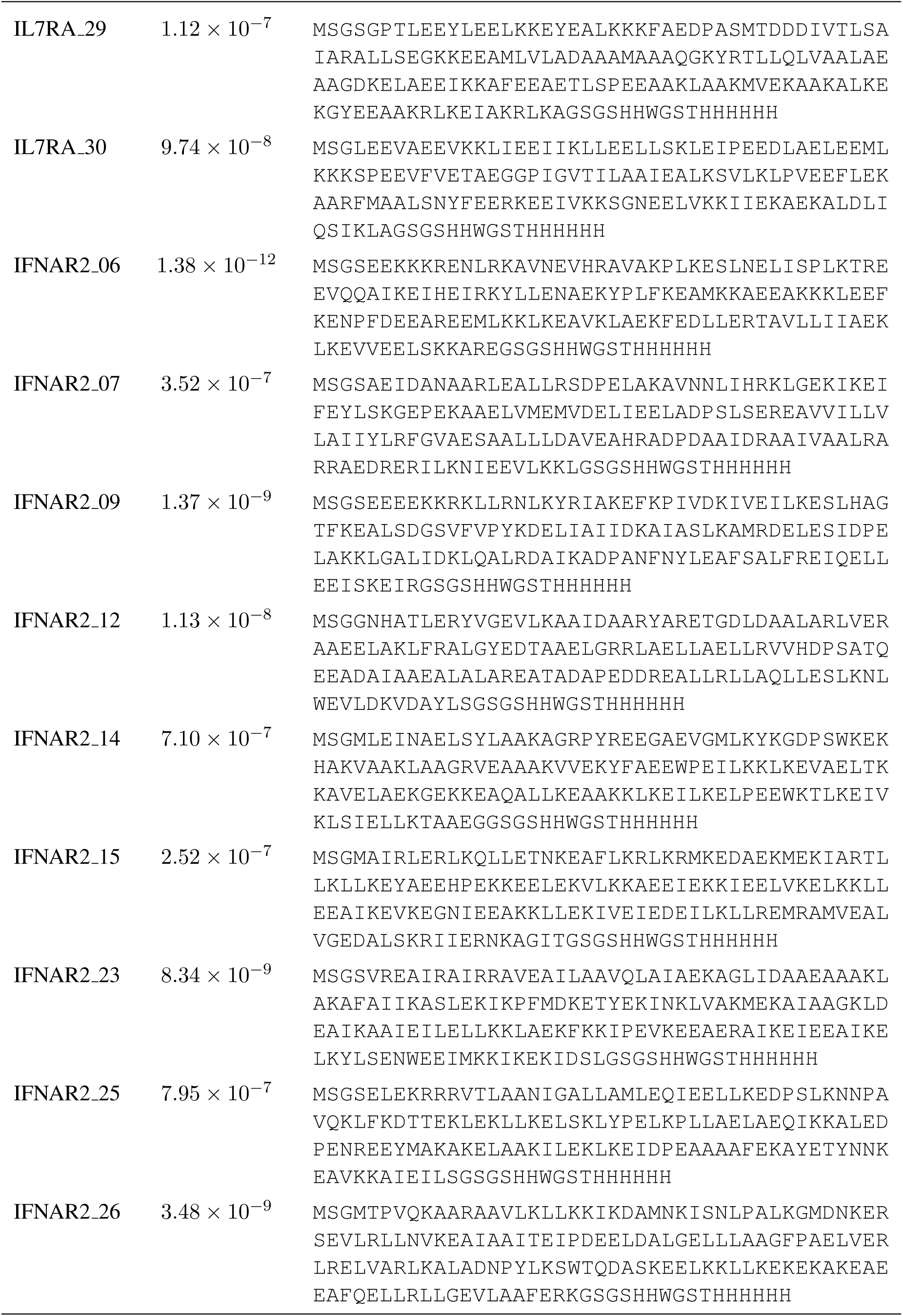
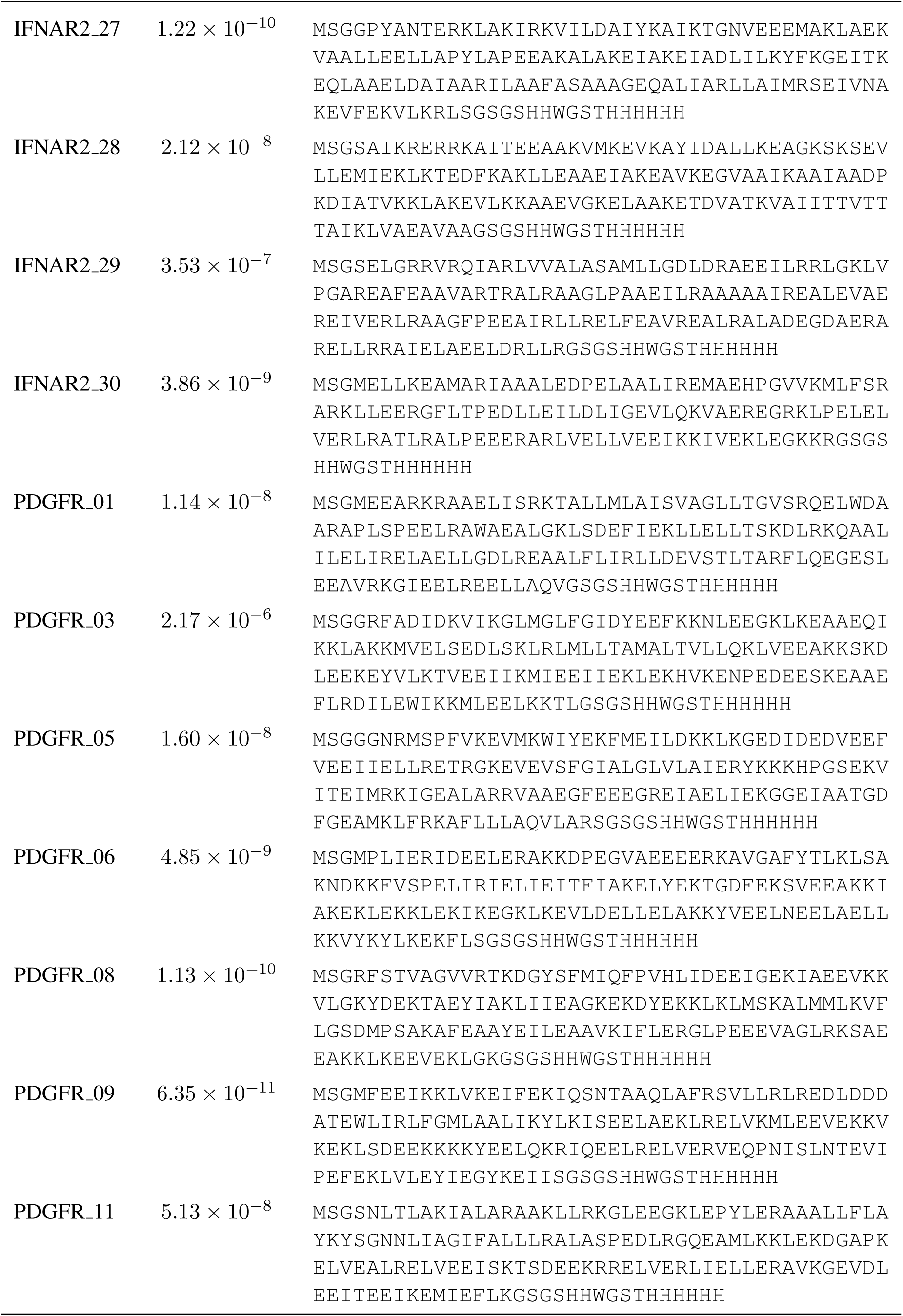
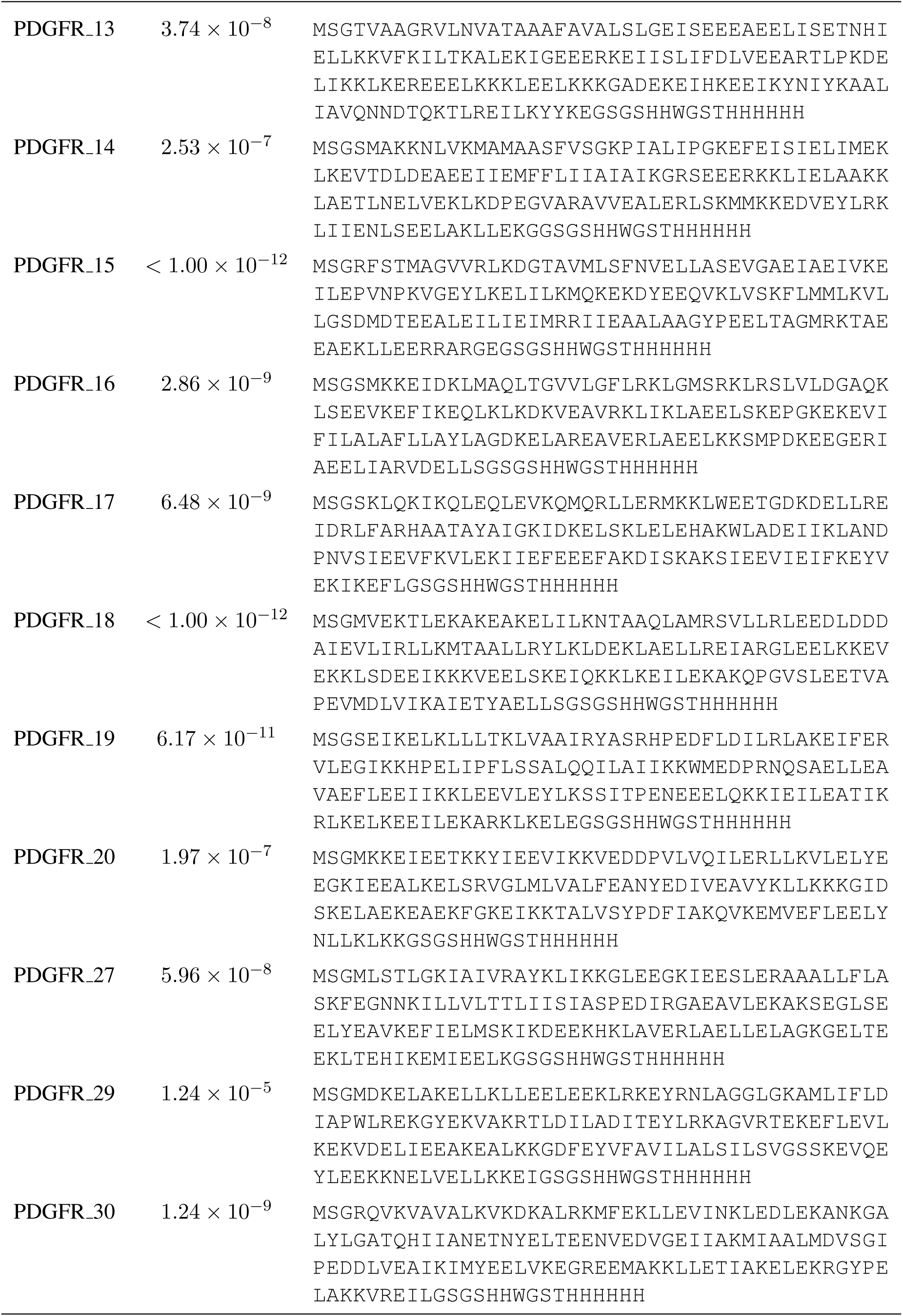
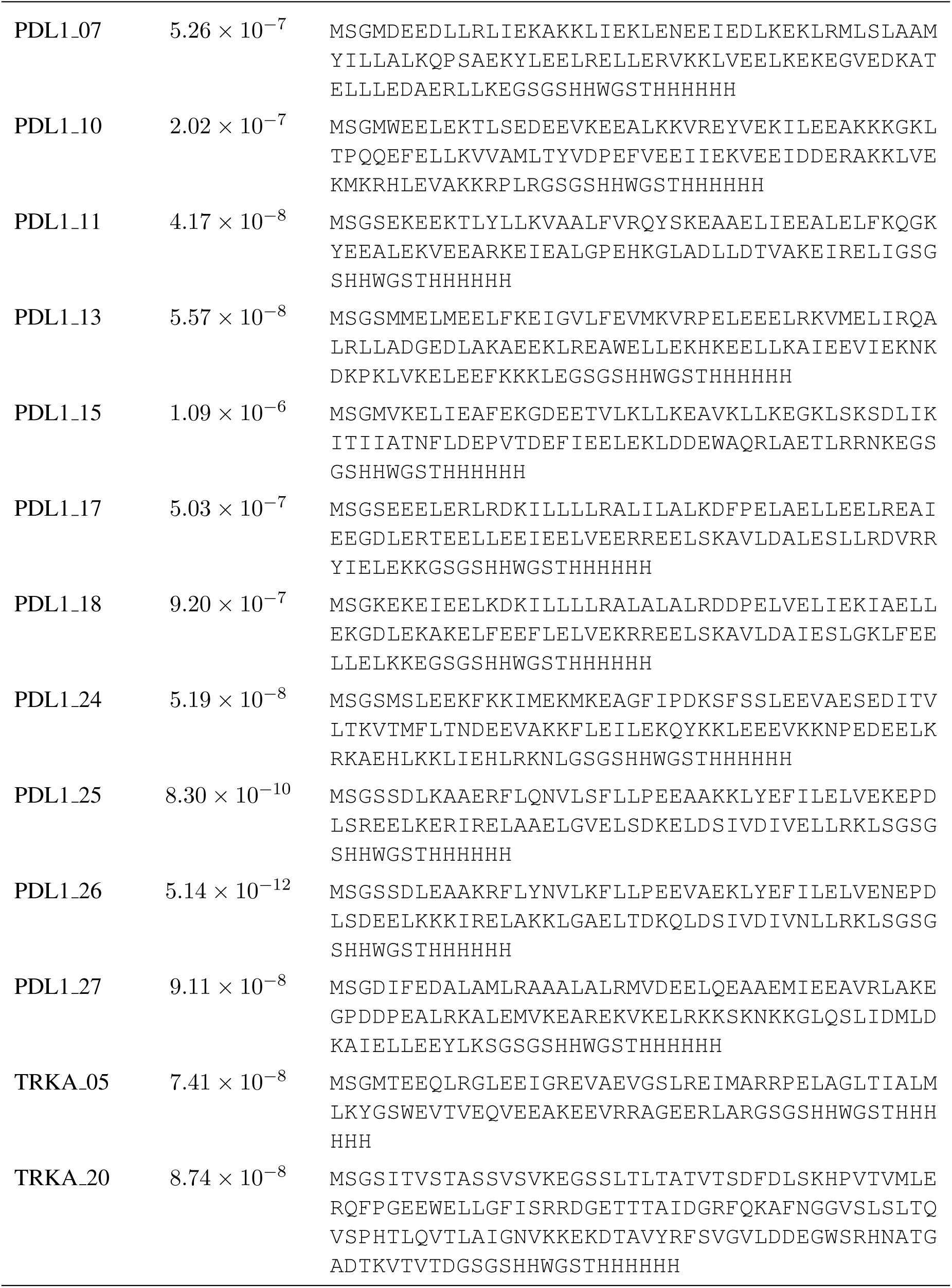
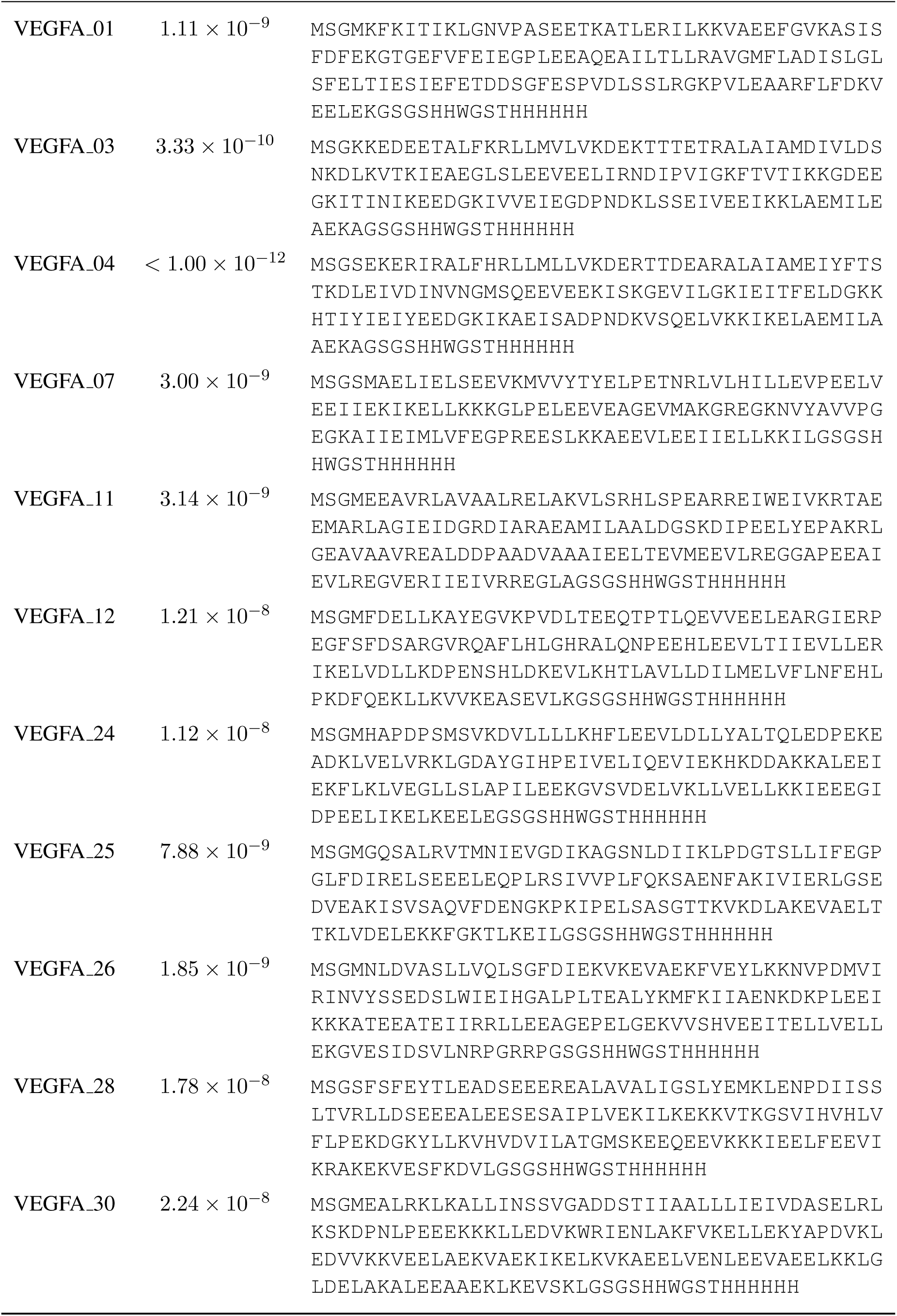
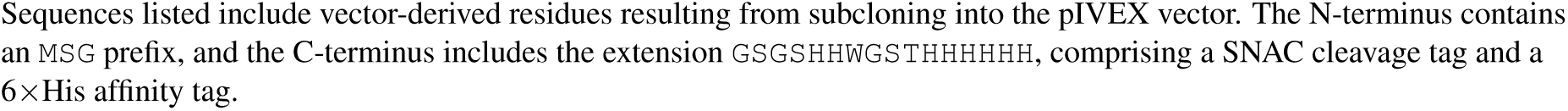
Sequences and binding affinities of all binders per target.

**Table S8:**
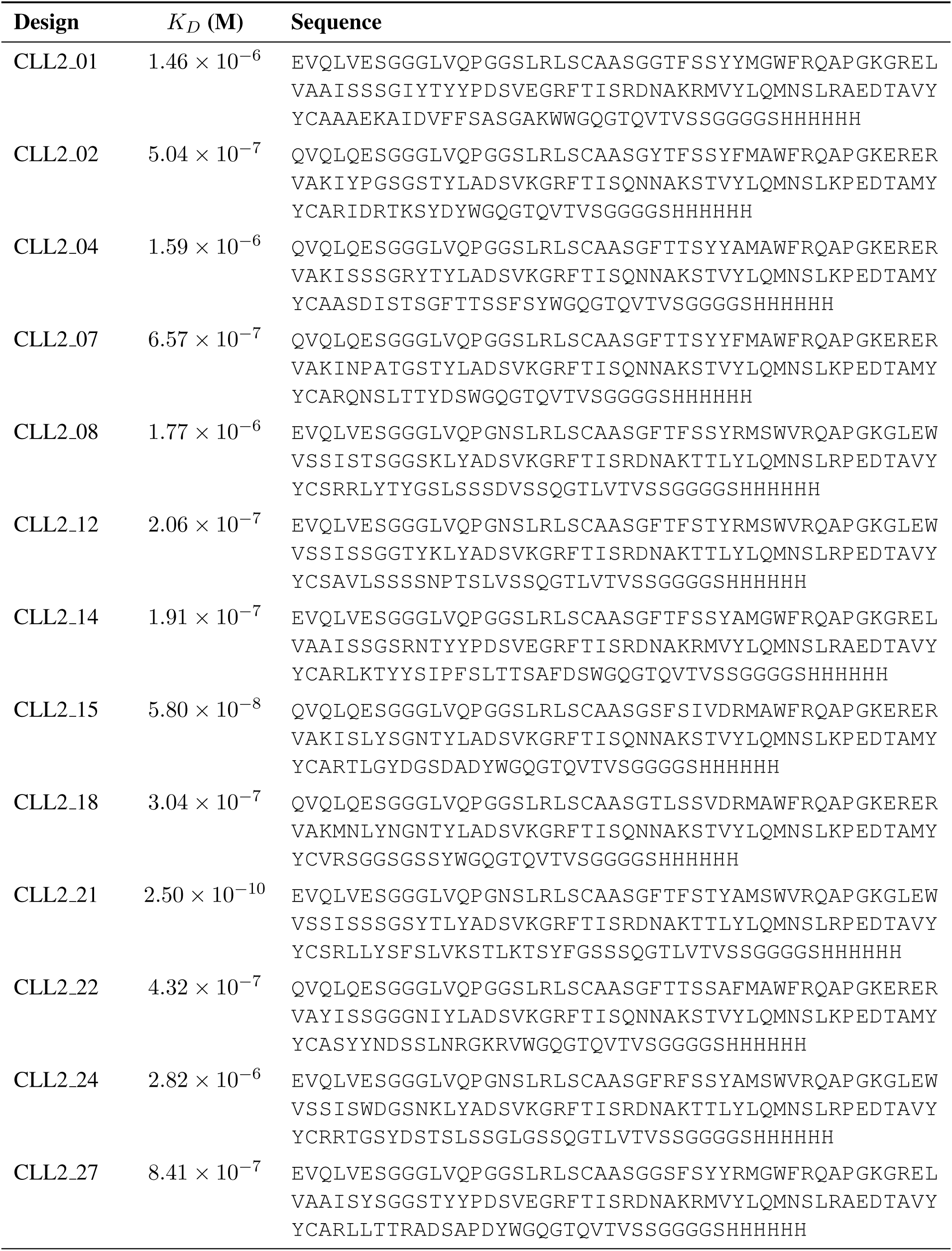
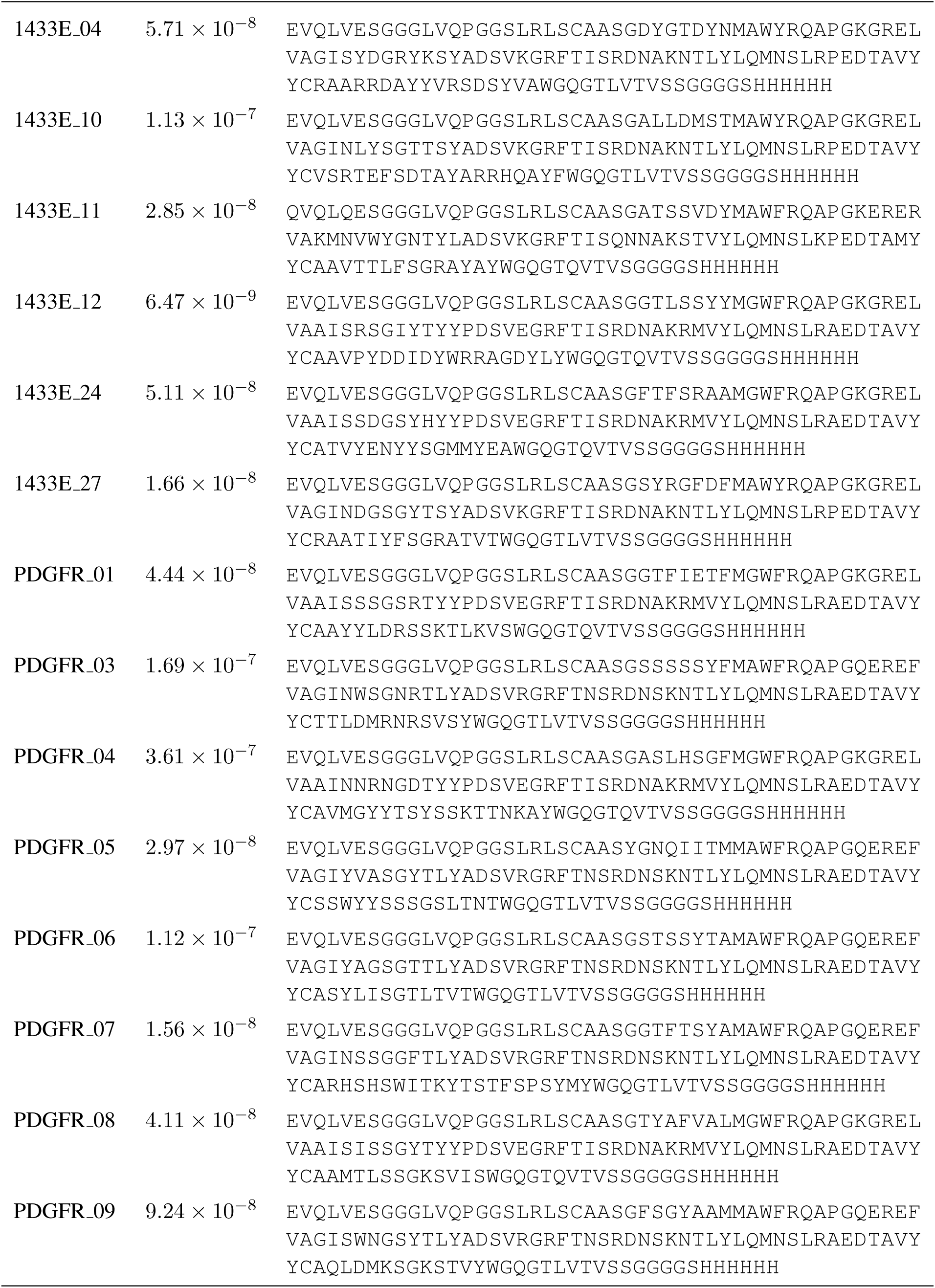
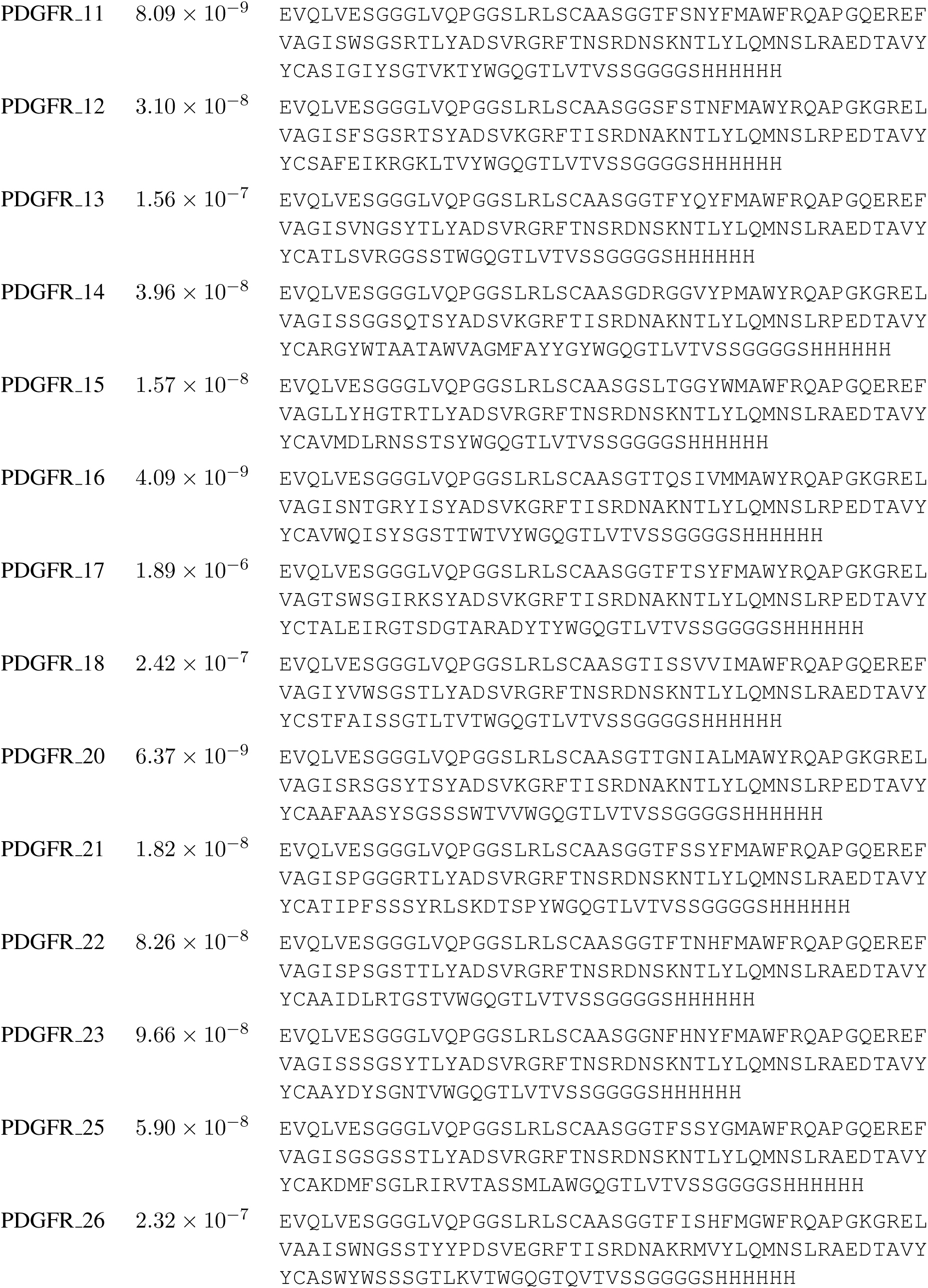
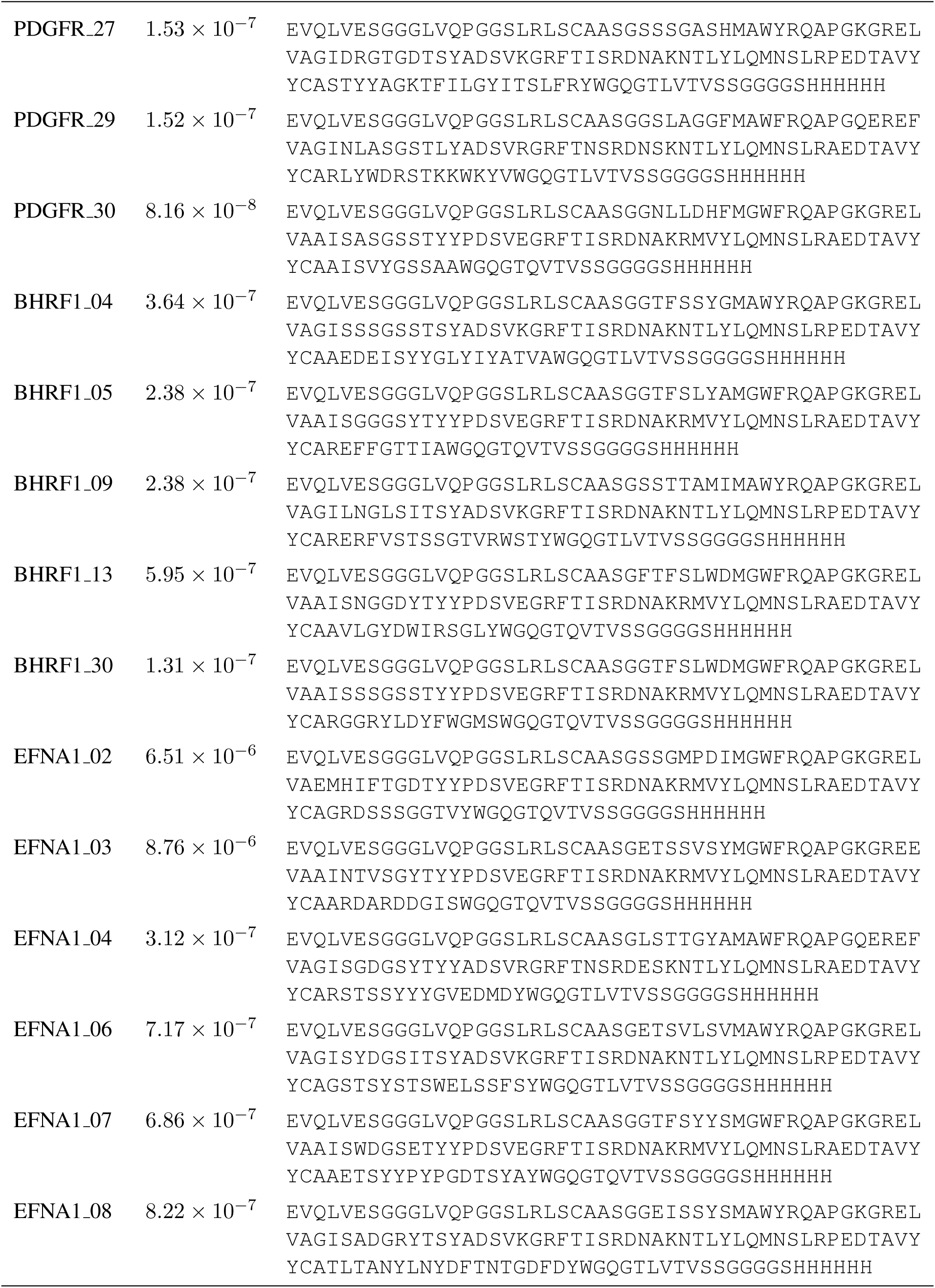
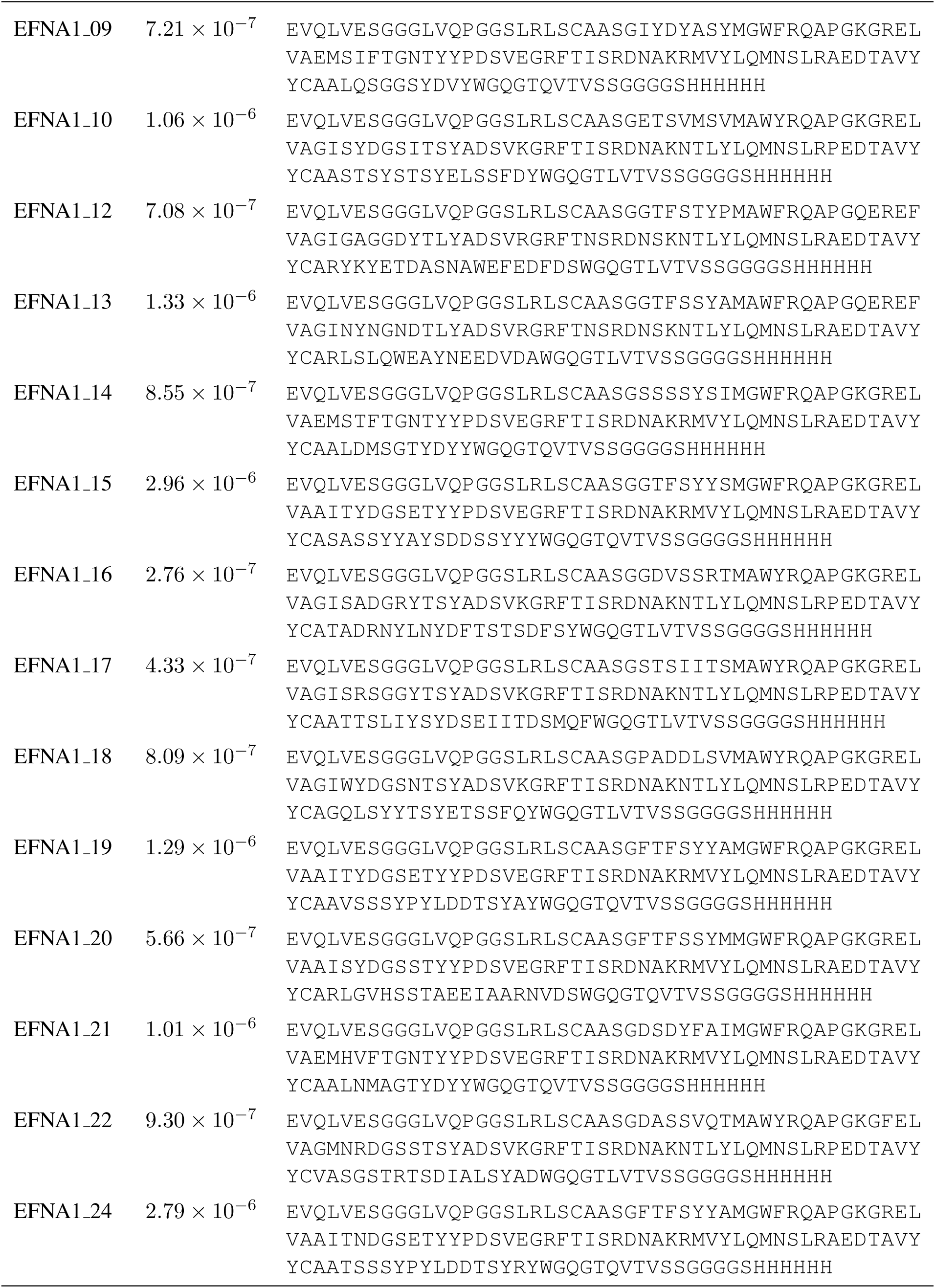
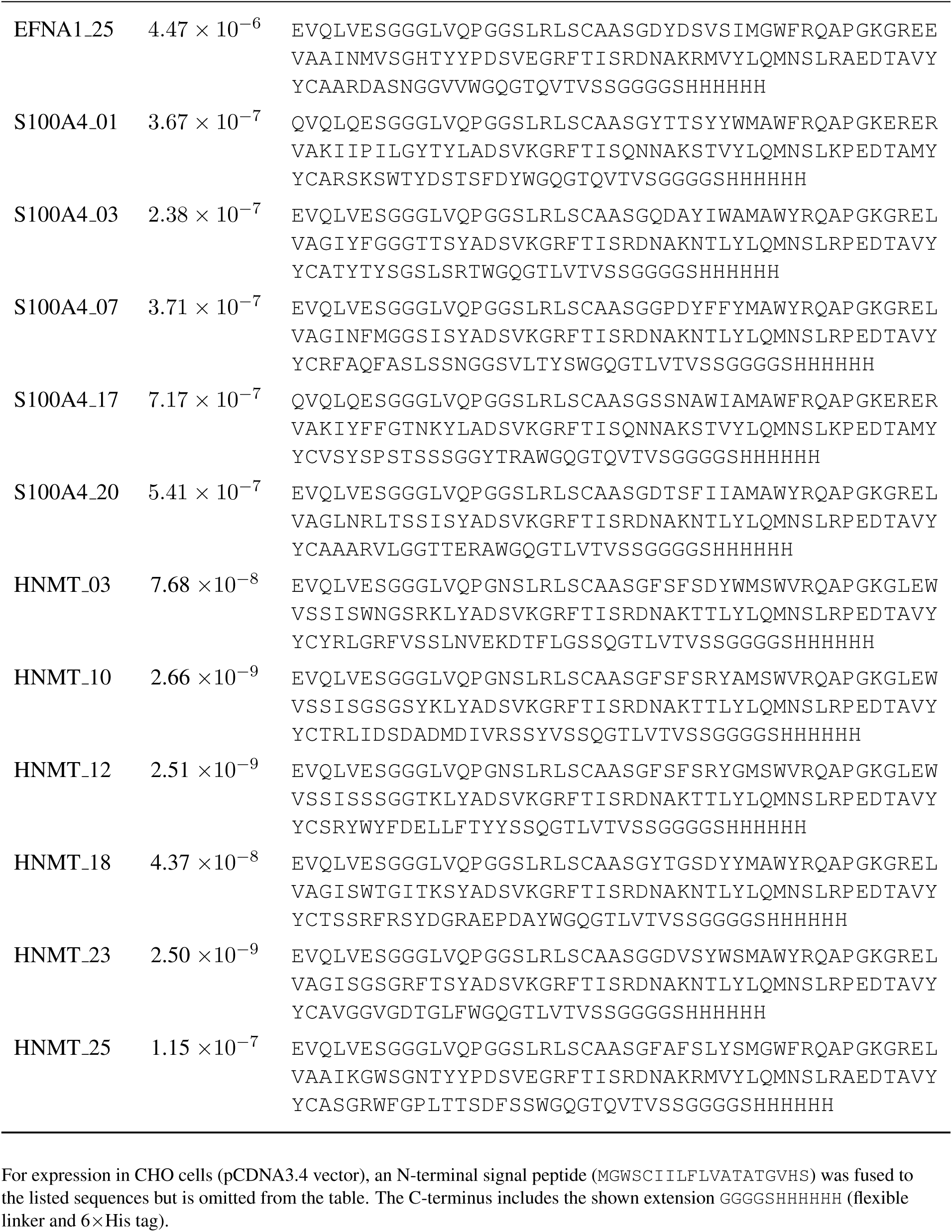
Sequences and binding affinities of all VHHs per target. Design *K_D_* (M) Sequence.

## References

[1] Longxing Cao, et al. “Design of protein-binding proteins from the target structure alone”. In: Nature 605.7910 (2022), pp. 551–560.

[2] Georges Köhler and Cesar Milstein. “Continuous cultures of fused cells secreting antibody of predefined specificity”. In: Nature 256.5517 (1975), pp. 495–497.

[3] George P Smith. “Filamentous fusion phage: novel expression vectors that display cloned antigens on the virion surface”. In: Science 228.4705 (1985), pp. 1315–1317.

[4] Greg Winter, et al. “Making antibodies by phage display technology”. In: Annual review of immunology 12.1 (1994), pp. 433–455.

[5] Frances H Arnold. “Directed evolution: bringing new chemistry to life”. In: Angewandte Chemie (International Ed. in English) 57.16 (2017), p. 4143.

[6] Frances H Arnold. “Design by directed evolution”. In: Accounts of chemical research 31.3 (1998), pp. 125–131.

[7] Ivan Anishchenko, et al. “De novo protein design by deep network hallucination”. In: Nature 600.7889 (2021), pp. 547–552.

[8] Jue Wang, et al. “Scaffolding protein functional sites using deep learning”. In: Science 377.6604 (2022), pp. 387–394.

[9] Milong Ren, et al. “Accurate and robust protein sequence design with CarbonDesign”. In: Nature Machine Intelligence 6.5 (2024), pp. 536–547.

[10] Joseph L Watson, et al. “De novo design of protein structure and function with RFdiffusion”. In: Nature (2023), pp. 1–3.

[11] Nathaniel R Bennett, et al. “Atomically accurate de novo design of antibodies with RFdiffusion”. In: Nature (2025), pp. 1–11.

[12] Martin Pacesa, et al. “One-shot design of functional protein binders with BindCraft”. In: Nature 646.8084 (2025), pp. 483–492.

[13] John B Ingraham, et al. “Illuminating protein space with a programmable generative model”. In: Nature 623.7989 (2023), pp. 1070–1078.

[14] Pablo Gainza, et al. “De novo design of protein interactions with learned surface fingerprints”. In: Nature 617.7959 (2023), pp. 176–184.

[15] Luis S Mille-Fragoso, et al. “Efficient generation of epitope-targeted de novo antibodies with Germinal”. In: bioRxiv (2025).

[16] Yaron Lipman, et al. “Flow matching for generative modeling”. In: International Conference on Learning Representations (2023).

[17] Ricky T. Q. Chen and Yaron Lipman. “Flow Matching on General Geometries”. In: International Conference on Learning Representations (2024).

[18] Jason Yim, et al. “Fast protein backbone generation with SE(3) flow matching”. In: arXiv preprint arXiv:2310.05297 (2023).

[19] Avishek Joey Bose, et al. “Se (3)-stochastic flow matching for protein backbone generation”. In: arXiv preprint arXiv:2310.02391 (2023).

[20] Yu Liu et al. “Af3score: A score-only adaptation of alphafold3 for biomolecular structure evaluation”. In: Journal of Chemical Information and Modeling 65.15 (2025), pp. 8207–8214.

[21] John Jumper, et al. “Highly accurate protein structure prediction with AlphaFold”. In: Nature 596.7873 (2021), pp. 583–589.

[22] Josh Abramson, et al. “Accurate structure prediction of biomolecular interactions with AlphaFold 3”. In: Nature 630.8016 (2024), pp. 493–500.

[23] Yeqing Lin, et al. “Out of many, one: Designing and scaffolding proteins at the scale of the structural universe with genie 2”. In: arXiv preprint arXiv:2405.15489 (2024).

[24] Tomas Geffner, et al. “Proteina: Scaling flow-based protein structure generative models”. In: arXiv preprint arXiv:2503.00710 (2025).

[25] Zhuoqi Zheng, et al. “MotifBench: A standardized protein design benchmark for motif-scaffolding problems”. In: arXiv preprint arXiv:2502.12479 (2025).

[26] Tianyu Lu, et al. “Conditional protein structure generation with protpardelle-1c”. In: bioRxiv (2025).

[27] Hannes Stark, et al. “Boltzgen: Toward universal binder design”. In: bioRxiv (2025), pp. 2025–11.

[28] Nathaniel R Bennett, et al. “Improving de novo protein binder design with deep learning”. In: Nature Communications 14.1 (2023), p. 2625.

[29] Justas Dauparas, et al. “Robust deep learning–based protein sequence design using ProteinMPNN”. In: Science 378.6615 (2022), pp. 49–56.

[30] Frédéric A Dreyer, et al. “Inverse folding for antibody sequence design using deep learning”. In: arXiv preprint arXiv:2310.19513 (2023).

[31] Rebecca F Alford et al. “The Rosetta all-atom energy function for macromolecular modeling and design”. In: Journal of chemical theory and computation 13.6 (2017), pp. 3031–3048.

[32] Matthias Glögl, et al. “Target-conditioned diffusion generates potent TNFR superfamily antagonists and agonists”. In: Science 386.6726 (2024), pp. 1154–1161.

[33] Jin Sub Lee and Philip M Kim. “FlowPacker: protein side-chain packing with torsional flow matching”. In: Bioinformatics 41.3 (2025), btaf010.

[34] Ian Sillitoe, et al. “CATH: increased structural coverage of functional space”. In: Nucleic acids research 49.D1 (2021), pp. D266–D273.

[35] Helen M Berman, et al. “The protein data bank”. In: Nucleic acids research 28.1 (2000), pp. 235–242.

[36] Martin Steinegger and Johannes Soding. “MMseqs2 enables sensitive protein sequence searching for the analysis of massive data sets”. In: Nature biotechnology 35.11 (2017), pp. 1026–1028.

[37] James Dunbar, et al. “SAbDab: the structural antibody database”. In: Nucleic acids research 42.D1 (2014), pp. D1140–D1146.

