## Supplementary Information for "High-Affinity Protein Binder Design via Flow Matching and In Silico Maturation"

**D Supplementary Figures** **25**

**E Supplementary Tables** **44**

### Appendix A Supplementary Methods

#### A.1 Notation

We denote the number of residues in a protein structure by $N$, and each residue is represented by a *frame* $T = (x, r) \in \mathrm{SE}(3)$, where $x \in \mathbb{R}^3$ denotes a translation vector and $r \in \mathrm{SO}(3)$ a rotation matrix. Our model generates backbone atoms, i.e. $\Omega = \{N, C, C_\alpha, O\}$, and we denote by $z \in \mathbb{R}^3$ the global coordinates of a single atom. For binder design tasks, we denote the target backbone atoms and the binder backbone atoms by $\mathcal{B}$ and $\mathcal{T}$ respectively. The set of atom pairs between the target and the binder whose backbone atoms are separated by a distance less than 10 Å is denoted by $\mathcal{P}$ which is used in loss computation.

We establish the following notation for operators employed in the subsequent algorithms. Capitalized operator names are exclusively reserved for modules containing learnable parameters. For instance, LINEAR signifies a parameterized linear transformation, defined by a weight matrix $W$ and a bias vector $b$. The term LAYERNORM specifically denotes layer normalization applied across the channel dimensions. Conversely, lowercase names are utilized for functions that operate without learnable parameters (e.g., `sigmoid`, `softmax`) and standard non-parameterized mathematical operations (e.g., `sin`, `cos`, and `exp`).

#### A.2 Datasets

##### A.2.1 Data Curation

We curated a unified structural dataset comprising three complementary data sources: monomeric protein structures, protein–protein interaction (PPI) complexes, and antibody-like binders, including VHH and scFv.

Monomeric protein structures were obtained from two sources: protein domains curated in the CATH database [1] and single-chain structures from the RCSB Protein Data Bank (PDB) [2]. We filtered the data based on the following criteria:

- non canonical amino acid ratio less than $20\%$.
- Sequence length is between 25 and 512.
- Coil percent is less than 0.8
- Radius of gyration is within the lowest $96\%$ of the distribution.

Sequence redundancy was reduced by clustering all sequences using MMseqs2 [3]. After filtering and deduplication, the monomer dataset comprised approximately 60k clusters, spanning a broad range of lengths and folds. This dataset was used primarily for unconditional structure learning, enabling the model to learn general protein structures.

We curated a protein–protein interaction (PPI) dataset consisting of both chain–chain interactions (CCI) and domain–domain interactions (DDI). We filter the data based on the following criteria:

- number of contact pairs larger than 80 where contact pair is defined as interchain residues within the distance of 10 Å.
- non canonical amino acid ratio less than $20\%$.
- structure similarity between target and binder is less than 0.6.
- binder radius of gyration divided by length less than $k$, where $k = 0.15$ if binder size is larger than 100, $k = 0.16$ if binder size is between 70 and 100, $k = 0.18$ if binder size is less than 70.
- binder main chain SASA ration is less than 17.
- Sequence length is less than 570.
- Binder sequence length is less than 256 and target sequence length is larger than 30.

After filtering and redundancy removal, the combined CCI + DDI dataset contained approximately 18k interaction pairs. This dataset was used to train the model to capture general interface geometry and inter-chain spatial organization, which are essential for downstream binder generation.

**Table S1: Input features to the model.** $N$ **is the number of residues.**

| Feature & Shape | Description |
|---|---|
| `aatype` $[N]$ | The input amino acid sequence. |
| `residue_index` $[N]$ | The index into the original amino acid sequence. |
| `chain_index` $[N]$ | 0 for the target chain and 1 for the binder chain. |
| `diffuse_mask` $[N]$ | 0 for the diffused regions to be generated and 1 for the conditioned regions (e.g., target or motifs). |
| `hotspot_mask` $[N]$ | Indicating whether a residue is a hotspot. |
| `noise_level` $[N]$ | Representing the amount of noise injected at the current timestep, ranging from 0 to 1. |
| `noised_trans` $[N, 3]$ | Noised CA coordinates. |

To specialize the model for antibody-like binder design, we curated a dataset of antibody, VHH and TCR structures from SAbDab [4]. The dataset comprised approximately 6.9k antibody structures, 1.2k VHH structures, and 350 TCR structures, clustered into 2.6k, 500 and 200 clusters, respectively.

###### A.2.2 Input Features

In Table S1, we list the input features to the model. These features will be embedded into the single embedding network and pair embedding network.

##### A.3 Input Embeddings

Initial embedding details are presented in Algorithm 1 and Algorithm 2.

**Algorithm 1: Embeddings for initial single representations**

**def** `SingleEmbedder`($\{\mathbf{f}_i^{aatype}\}, \{\mathbf{f}_i^{residue\_index}\}, \{\mathbf{f}_i^{chain\_index}\}, \{\mathbf{f}_i^{diffuse\_mask}\}, \{\mathbf{f}_i^{hotspot\_mask}\}, \{\mathbf{f}_i^{noise\_level}\}$) **:**

$\mathbf{r}_i = \text{ResidueIndexEmbedder}(\mathbf{f}_i^{\text{residue\_index}})$

$\mathbf{n}_i = \text{NoiseLevelEmbedder}(\mathbf{f}_i^{\text{noise\_level}})$

$\mathbf{a}_i = \text{Linear}(\text{one\_hot}(\mathbf{f}_i^{\text{aatype}}))$

$\mathbf{o}_i = \text{concat}(\mathbf{r}_i, \mathbf{n}_i, \mathbf{a}_i, \mathbf{f}_i^{\text{chain\_index}}, \mathbf{f}_i^{\text{diffuse\_mask}}, \mathbf{f}_i^{\text{hotspot\_mask}})$

$\mathbf{s}_i = \text{Linear}(\mathbf{o}_i)$ $\qquad \mathbf{s}_i \in \mathbb{R}^{c_s},\ c_s = 256$

return $\{\mathbf{s}_\mathbf{i}\}$

**Algorithm 2: Embeddings for initial pair representations**

**def** `PairEmbedder`($\{\mathbf{s}_i\}, \{\mathbf{f}_i^{residue\_index}\}, \{\mathbf{f}_i^{chain\_index}\}, \{\mathbf{f}_i^{diffuse\_mask}\}, \{\mathbf{f}_i^{noised\_trans}\}$) **:**

$\mathbf{a}_i = \text{Linear}(\mathbf{s}_i)$

$\mathbf{n}_{ij} = \text{concat}(\mathbf{a}_i, \mathbf{a}_j)$

$\mathbf{r}_{ij} = \text{relpos}\left(\{\mathbf{f}_i^{\text{residue\_index}}\}\right)$

$\{\mathbf{D}_{ij}\} = \text{distogram}\left(\{\mathbf{f}_i^{\text{noised\_trans}}\}\right)$

$\mathbf{m}_{ij} = \text{concat}(\mathbf{f}_i^{\text{diffuse\_mask}}, \mathbf{f}_j^{\text{diffuse\_mask}})$

$\mathbf{e}_{ij} = \text{concat}(\mathbf{n}_{ij}, \mathbf{r}_{ij}, \mathbf{D}_{ij}, \mathbf{m}_{ij})$

$\mathbf{z}_{ij} = \text{Linear}(\mathbf{e}_{ij})$ $\qquad \mathbf{z}_{ij} \in \mathbb{R}^{c_z \times c_z},\ c_z = 128$

return $\{\mathbf{z}_{ij}\}$

**Algorithm 3: Embeddings for residue index**

**def** `ResidueIndexEmbedder` $(\{\mathbf{f}_i^{\textit{residue\_index}}\}, \textit{embed\_size} = 128, \textit{max\_len} = 2056)$ **:**

$k = \text{arange}\left(\frac{\text{embed\_size}}{2}\right)$

$\mathbf{p}_i = \sin(\mathbf{f}_i^{\text{residue\_index}} \times \frac{\pi}{(\text{max\_len})^{(2k/\text{embed\_size})}})$

$\mathbf{q}_i = \cos(\mathbf{f}_i^{\text{residue\_index}} \times \frac{\pi}{(\text{max\_len})^{(2k/\text{embed\_size})}})$

$\mathbf{r}_i = \text{concat}(\mathbf{p}_i, \mathbf{q}_i)$

return $\{\mathbf{r}_i\}$

**Algorithm 4: Embeddings for noise level**

**def** `NoiseLevelEmbedder` $(\{\mathbf{f}_i^{\textit{noise\_level}}\}, \textit{embed\_size} = 128, \textit{max\_positions} = 2056)$ **:**

$t_i = \mathbf{f}_i^{\text{noise\_level}} \times \text{max\_positions}$

$\text{half\_dim} = \left\lfloor \frac{\text{embed\_size}}{2} \right\rfloor$

$\text{scale} = \frac{\log(\text{max\_positions})}{\text{half\_dim} - 1}$

$\text{freq} = \exp(-\text{arange}(\text{half\_dim}) \times \text{scale})$

$\mathbf{p}_i = t_i \times \text{freq}$

$\mathbf{n}_i = \text{concat}(\sin(\mathbf{p}_i), \cos(\mathbf{p}_i))$

return $\{\mathbf{n}_i\}$

**Algorithm 5: Relative position encoding**

**def** `relpos` $(\{\mathbf{f}_i^{\textit{residue\_index}}\})$ **:**

$\mathbf{d}_{ij} = \mathbf{f}_i^{\text{residue\_index}} - \mathbf{f}_j^{\text{residue\_index}}$

$\mathbf{r}_{ij} = \text{ResidueIndexEmbedder}(\mathbf{d}_{ij})$

$\mathbf{r}_{ij} = \text{Linear}(\mathbf{r}_{ij})$

return $\{\mathbf{r}_{ij}\}$

**Algorithm 6: Compute distogram from atomic coordinates**

**def** `distogram(`$\{\mathbf{p}_i\}, \mathbf{d}_{\min}, \mathbf{d}_{\max}, K$`)`**:**

  $\mathbf{d}_{ij} \leftarrow \|\mathbf{p}_i - \mathbf{p}_j\|_2$

  $\mathbf{b}_k = \mathbf{d}_{\min} + (k-1)\frac{\mathbf{d}_{\max} - \mathbf{d}_{\min}}{K-1}$ for $k = 1 \dots K$

  **for** $i, j = 1 \dots N$ **do**

    **for** $k = 1 \dots K$ **do**

      $\mathbf{D}_{ij,k} \leftarrow 1[\,\mathbf{b}_k < \mathbf{d}_{ij} \leq \mathbf{b}_{k+1}\,]$

  return $\{\mathbf{D}_{ij}\}$

#### A.4 Model Architecture

Our model follows a hierarchical architecture that integrates both sequence-level and structure-level reasoning. Specifically, we employ two key components that have become standard in protein structure modeling: the **Pairformer** and the **Invariant Point Attention (IPA)** module.

The Pairformer module is used to jointly model residue-wise and pairwise representations through interleaved attention and transition layers. It allows effective information exchange between the one-dimensional sequence features and the two-dimensional pairwise geometric features, enabling the network to reason about both local and long-range residue interactions. We adopt the standard Pairformer design as introduced in AlphaFold-style architectures, without modification. We use 4 Pairformer blocks in our model.

Following AlphaFold [5], we use the Invariant Point Attention module to transform pairwise relational information into 3D coordinate updates. The IPA performs attention over residue embeddings in SE(3)-invariant space, ensuring that the resulting structural representations are equivariant to rotations and translations. This design allows the model to learn spatial relationships in a physically meaningful and symmetry-preserving manner. Our model includes 6 IPA layers.

Our model consists of 23.2M parameters. Together, these modules enable our model to integrate sequence context, pairwise geometry, and spatial reasoning in a unified framework for protein structure generation and refinement.

##### A.5 Loss Functions

**Flow matching loss in** $\mathrm{SE}(3)$**.** Following the flow-matching framework, the model learns to predict the continuous velocity field that transforms a noisy structure toward its clean counterpart. Each residue is represented in SE(3) space by a translation vector $x \in \mathbb{R}^3$ and a rotation matrix $r \in \mathrm{SO}(3)$.

Given a clean structure $\mathbf{T}_1$ and a noisy structure $\mathbf{T}_t$ sampled along the flow trajectory, the target velocity is defined as the time derivative of the flow:

$$\mathbf{v}_t = \frac{d\mathbf{T}_t}{dt}. \tag{1}$$

The SE(3) flow-matching loss minimizes the discrepancy between predicted and ground-truth velocities on both translational and rotational components:

$$\mathcal{L}_{\text{flow}} = \mathbb{E}_{t,p_0,p_1}\left[\sum_{i=1}^{N}\left(\left\|\hat{\mathbf{v}}_{t,x}^{(i)}(\mathbf{T}_t,t) - \mathbf{v}_{t,x}^{(i)}\right\|_{\mathbb{R}^3}^2 + \left\|\hat{\mathbf{v}}_{t,r}^{(i)}(\mathbf{T}_t,t) - \mathbf{v}_{t,r}^{(i)}\right\|_{\mathrm{SO(3)}}^2\right)\right],$$

where $p_0$ and $p_1$ denote noise distribution and data distribution respectively, $p_0 = \mathcal{N}(0, I_3) \otimes \mathcal{U}(\mathrm{SO}(3))$, $t \sim \mathcal{U}([0,1])$.

Following FrameFlow [6], the model predicts the clean frames $\mathbf{T}_1$ given the corrupted frames $\mathbf{T}_t$ at time $t$, and the velocity can be approximated as

$$\hat{\mathbf{v}}_{t,x}^{(i)} = \frac{\hat{\mathbf{x}}_1^{(i)} - \mathbf{x}_t^{(i)}}{1-t} \tag{2}$$

$$\hat{\mathbf{v}}_{t,r}^{(i)} = \frac{\log_{\mathbf{r}_t^{(i)}}(\hat{\mathbf{r}}_1^{(i)})}{1-t}, \tag{3}$$

where $\log_{\mathbf{r}_t^{(i)}}$ is the logarithmic map on $\mathrm{SO}(3)$. As a result, the objective can be reparameterized as

$$\mathcal{L}_{\text{flow}} = \mathbb{E}_{t,\,p_0,\,p_1}\left[\frac{1}{(1-t)^2}\sum_{i=1}^{N}\Big(\|\hat{\mathbf{x}}_1^{(i)}(\mathbf{T}_t,t) - \mathbf{x}_1^{(i)}\|^2_{\mathbb{R}^3}\right.$$
$$\left. + \|\log_{\mathbf{r}_t^{(i)}}\big(\hat{\mathbf{r}}_1^{(i)}(\mathbf{T}_t,t)\big) - \log_{\mathbf{r}_t^{(i)}}\big(\mathbf{r}_1^{(i)}\big)\|^2_{\text{SO}(3)}\Big)\right]. \tag{4}$$

**Interface contact loss.** We introduce interface contact loss to improve the learning of target to binder interaction. Since we observed that some sampled results positioned binders too far from the target, leading to weak contacts, we designed this interface contact loss to improve the quality of interfaces. This loss is defined on residue pairs across binder and target. We examine on interchain residue pairs within 10 Å in native structures, and compare the predicted pairwise distances of these pairs against true pairwise distances. Let $\mathcal{P} = \big\{(i,j) \mid z_i \in \mathcal{B}, z_j \in \mathcal{T}, \|z_i - z_j\|_2 \leq 10\,\text{Å}\big\}$, where $z_i$ and $\hat{z}_i$ are the ground-truth and predicted coordinates of backbone atoms respectively. Then the interface contact loss can be written as

$$\mathcal{L}_{\text{interface}} = \frac{1}{|\mathcal{P}|}\sum_{(i,j)\in\mathcal{P}} \|\|\hat{z}_i - \hat{z}_j\|_2 - \|z_i - z_j\|_2\|_2^2. \tag{5}$$

**Backbone atom loss.** The backbone atom coordinates are reconstructed from the predicted translation and rotation vectors, and we compute the mean squared error (MSE) between the predicted and ground-truth coordinates of the backbone atoms. The backbone atom loss is defined as

$$\mathcal{L}_{\text{bb}} = \frac{1}{|\mathcal{B}|}\sum_{i\in\mathcal{B}} \|\hat{z}_i - z_i\|_2^2. \tag{6}$$

**Binder pairwise distance loss.** By minimizing binder pairwise distance loss, the model is trained to accurately predict the spatial relationship (the pairwise distances) between residues in the binder, which is crucial for generating physically realistic proteins. This loss is defined as

**Algorithm 7: SE(3) flow matching interpolation**

**def** `Interpolant` $(\mathbf{T}_0, \mathbf{T}_1, t)$**:**

  **for** $i = 1, \ldots, N$ **do**

    `// Translations`

    $\mathbf{x}_t^{(i)} \leftarrow (1-t)\mathbf{x}_0^{(i)} + t\mathbf{x}_1^{(i)}$

    `// Rotations`

    $\mathbf{r}_t^{(i)} \leftarrow \exp_{\mathbf{r}_0^{(i)}}((1-e^{-ct}) \log_{\mathbf{r}_0^{(i)}}(\mathbf{r}_1^{(i)}))$

    $\mathbf{T}_t^{(i)} \leftarrow (\mathbf{x}_t^{(i)}, \mathbf{r}_t^{(i)})$

  **return** $\mathbf{T}_t$

**Algorithm 8: PPIFlow training**

**def** *Train***:**

  **while** *not converged* **do**

    $\mathbf{T}_1 \sim p_1$

    $\mathbf{T}^{(0)} \leftarrow$ `SampleInit`$(N)$

    $t \sim \mathcal{U}([0, 1])$

    $\mathbf{T}_t \leftarrow$ `Interpolant`$(\mathbf{T}_0, \mathbf{T}_1, t)$

    **if** $\text{Uniform}(0, 1) < 0.5$ **then**

      `// Train step without self-conditioning`

      $\hat{\mathbf{T}}_1^{\text{sc}} \leftarrow \mathbf{0}$

    **else**

      `// Train step with self-conditioning`

      $\hat{\mathbf{T}}_1^{\text{sc}} \leftarrow$ `PPIFlow`$(\mathbf{T}_t, t, \mathbf{0})$

      $\hat{\mathbf{T}}_1^{\text{sc}} \leftarrow$ `StopGradient`$(\hat{\mathbf{T}}_1^{\text{sc}})$

    $\hat{\mathbf{T}}_1 \leftarrow$ `PPIFlow`$(\mathbf{T}_t, t, \hat{\mathbf{T}}_1^{\text{sc}})$

    $\theta = \text{optimizer}(\theta, \mathcal{L}, \mathbf{T}_1, \hat{\mathbf{T}}_1)$

$$\mathcal{L}_{\text{dist}} = \frac{1}{|\mathcal{B}|^2} \sum_{i,j \in \mathcal{B}} \| \|\hat{z}_i - \hat{z}_j\|_2 - \|z_i - z_j\|_2 \|_2^2. \tag{7}$$

**Algorithm 9: PPIFlow inference**

**def** `SampleInit` $(N)$:
  // Random initial structure for $N$ residues
  **for** $i = 1, \ldots, N$ **do**
    $\mathbf{x}_0^{(i)} \sim \mathcal{N}(0, I_3)^N$
    $\mathbf{r}_0^{(i)} \sim \mathcal{U}(\mathrm{SO}(3))^N$
    $\mathbf{T}_0^{(i)} \leftarrow (\mathbf{x}_0^{(i)}, \mathbf{r}_0^{(i)})$
  **return** $\mathbf{T}_0$

**def** `ReverseStep` $(\mathbf{T}_t, \hat{\mathbf{T}}_1, c = 10)$:
  // One step of reverse flow
  **for** $i = 1, \ldots, N$ **do**
    $(\mathbf{x}_t^{(i)}, \mathbf{r}_t^{(i)}) \leftarrow \mathbf{T}_t^{(i)}$
    $(\hat{\mathbf{x}}_1^{(i)}, \hat{\mathbf{r}}_1^{(i)}) \leftarrow \hat{\mathbf{T}}_1^{(i)}$
    // Update translations
    $\mathbf{x}_{t+1/k}^{(i)} \leftarrow \frac{\hat{\mathbf{x}}_1^{(i)} - \mathbf{x}_t^{(i)}}{1-t} \times \frac{1}{k}$
    // Update rotations
    $\mathbf{r}_{t+1/k}^{(i)} \leftarrow \exp_{\mathbf{r}_t^{(i)}}(c \times \log_{\mathbf{r}_t^{(i)}}(\hat{\mathbf{r}}_1^{(i)}))$
    $\mathbf{T}_{t+1/k}^{(i)} \leftarrow (\mathbf{x}_{t+1/k}^{(i)}, \mathbf{r}_{t+1/k}^{(i)})$
  **return** $\mathbf{T}_{t+1/k}$

**def** `Sample` $(N, k = 100)$:
  // Generation of $N$-residue backbone structure
  $\mathbf{T}_0 \leftarrow$ `SampleInit` $(N)$
  $\hat{\mathbf{T}}_1^{\mathrm{sc}} \leftarrow \mathbf{0}$     // Initialize self-conditioning
  **for** $t = 0, 1/k, \ldots, (k-1)/k$ **do**
    $\hat{\mathbf{T}}_1 \leftarrow$ `PPIFlow` $(\mathbf{T}_t, t, \hat{\mathbf{T}}_1^{\mathrm{sc}})$
    $\mathbf{T}_{t+1/k} \leftarrow$ `ReverseStep` $(\mathbf{T}_t, \hat{\mathbf{T}}_1)$
    $\hat{\mathbf{T}}_1^{\mathrm{sc}} \leftarrow \hat{\mathbf{T}}_1$
  **return** $\mathbf{T}_1$

**Table S2: Binder design target protein**

| Protein | Company | Description | Cat | Accession No. | Protein structure |
|---|---|---|---|---|---|
| TrkA | ACRO | Human TrkA/NTRK1 (33-417) Protein, Mouse IgG2a Fc Tag | TRA-H5259 | P04629-2 | TrkA(ALA33-Gly417)-mFc(glu98-lys330) |
| IFNAR2 | ACRO | Human IFN-alpha/beta R2 Protein, Fc Tag | IF2-H5255 | P48551-2 | IFNAR2(Ile27-Lys243)-Fc(Pro100-Lys330) |
| IL17A | KACTUS | Biotinylated Human IL-17A/CTLA-8 Protein | ILA-HM418B | Q16552-1 | IL-17A(Gly24-Ala155)-His-Avi |
| IL7RA | ACRO | Human IL-7 R alpha/CD127 Protein, Fc Tag | ILA-H525a | NP-054862.1 | IL-7RA(Glu21-Asp239)-Fc(Pro100-Lys330) |
| PD-L1 | ACRO | Human PD-L1/B7-H1 Protein (MALS verified) | PD1-H5258 | NP-054862.1 | PD-L1(Phe19-Arg238)-Fc(Pro100-Lys330) |
| PDGFR | ACRO | Human PDGF R beta/CD140b Protein, Fc Tag | PDB-H5259 | P09619-1 | PDGFRB(Leu33-Phe530)-Fc(Pro100-Lys330) |
| VEGF165 | ACRO | Human VEGF165 Protein, premium grade | VE5-H4210 | P15692-4 | VEGF165(Ala27-Arg191) |

**Table S3: VHH design target protein**

| Protein | Company | Description | Cat No. | Accession | Protein structure |
|---|---|---|---|---|---|
| EFNA1 | SinoBio | Recombinant Human Ephrin-A1 (hFc Tag) | 10882-H02H | NP-004419.2 | (Met1-Ser182)-hFc |
| PDGFR | SinoBio | Recombinant Human PDGFRB (ECD), HPLC-verified | 10514-H02H1 | NP-002600.1 | (Met1-Phe530)-hFc |
| 1433E | SinoBio | Recombinant Human 14-3-3 epsilon | 10842-HNCE | NP-006752.1 | Gly-Pro-(Met1-Gln255) |
| IL-13 | SinoBio | Recombinant Human IL-13 (hFc Tag), HPLC-verified | 10369-H01H | AAK53823.1 | hFc-(Gly21-Asn132) |
| CCL2 | KACTUS | Biotinylated Human MCP-1/CCL2 | MCP-HM401B | P13500 | (Gln24-Thr99)-His-Avi |
| BHRF1 | Oritop | BHRF1 | H202511 1402 | P03182 | His-SUMO-(Met1-Ser165) |
| S100A4 | SinoBio | Recombinant Human S100A4 Protein (hFc Tag) | 10185-H01H | NP-002952.1 | hFc-(Met1-Lys101) |
| HNMT | SinoBio | Recombinant Human HNMT Protein (GST Tag) | 13096-H09E | AAH20677.1 | GST-(Met1-Ala292) |

### Appendix D Supplementary Figures

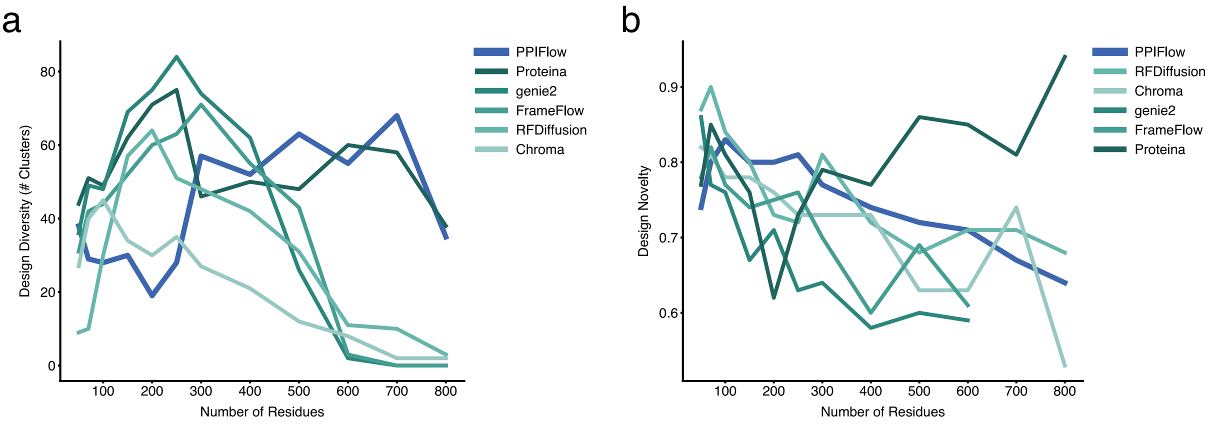

**Figure S1: Benchmarking monomer generation performance across varying lengths.**

**(a)** Evaluation of structural diversity in monomer generation tasks for sequences ranging from 50 to 800 amino acids (100 designs per length). PPIFlow is compared against Proteina, Genie2, FrameFlow, RFDiffusion, and Chroma. **(b)** Structural novelty analysis quantified by the maximum TM-score of successful designs against the PDB databases.

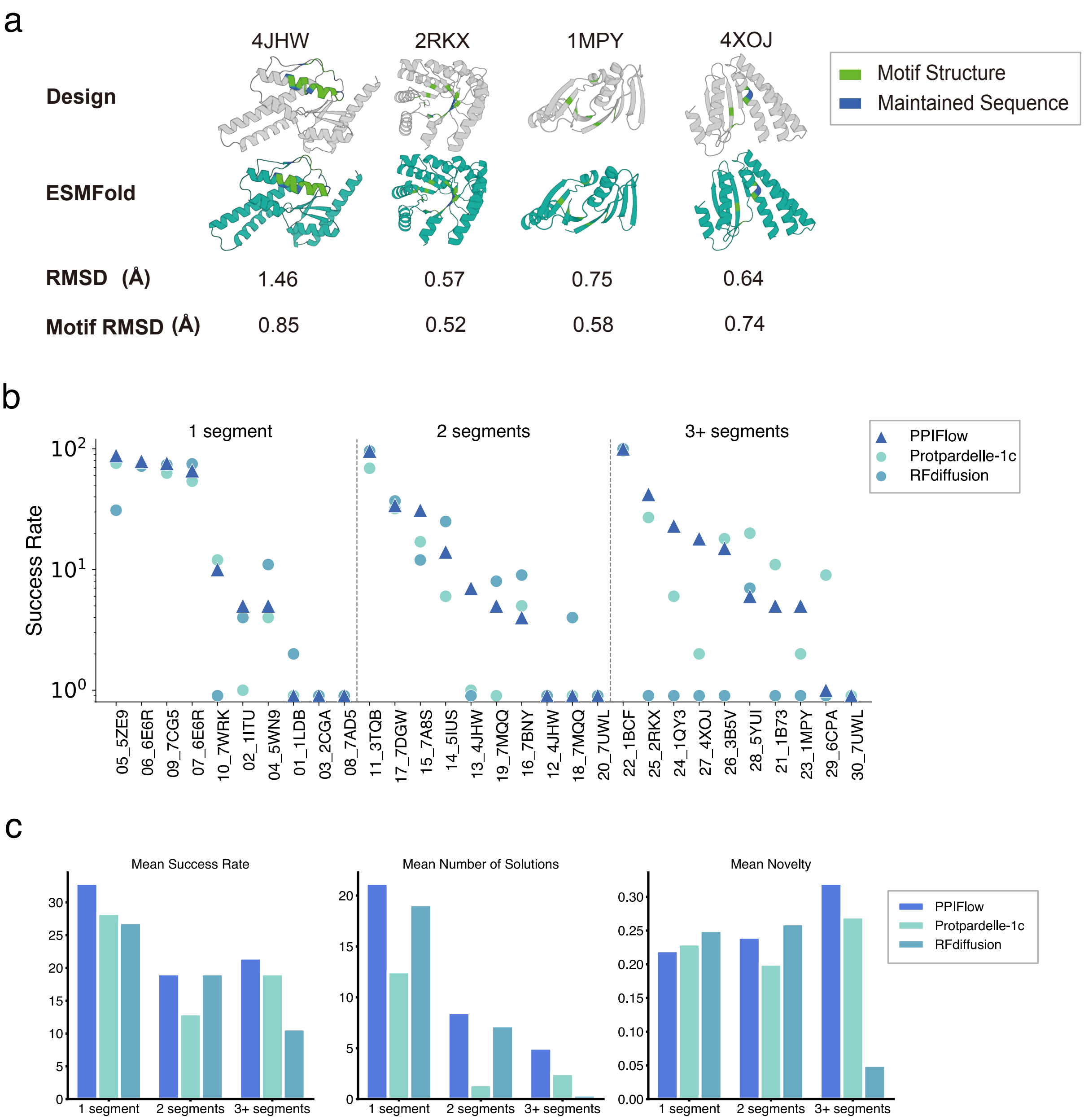

**Figure S2: Assessment of motif scaffolding performance and structural fidelity.**

**(a)** Performance of PPIFlow on four representative scaffolding tasks (PDB IDs: 4JHW, 2RKX, 1MPY, and 4XOJ). Panels show the alignment between PPIFlow-generated backbones and ESMFold-refolded structures, annotated with global C$\alpha$ RMSD and motif-specific C$\alpha$ RMSD values. **(b)** Comparison of design success rates on the MotifBench dataset between PPIFlow, Protpardelle-1c, and RFDiffusion. **(c)** Comprehensive comparison of the three models across mean success rate, solution diversity, and structural novelty.

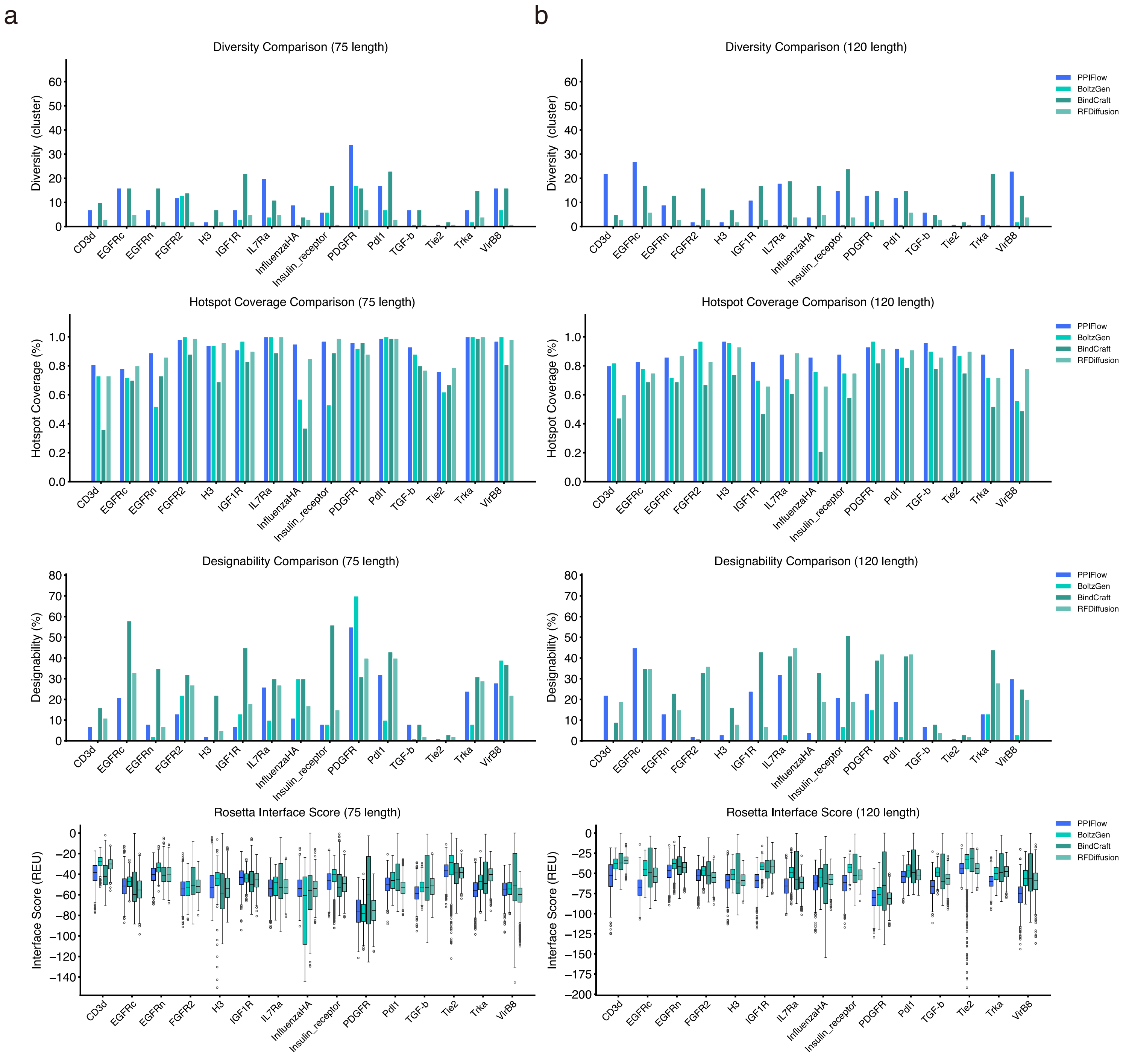

a
b
Diversity Comparison (75 length)
Diversity Comparison (120 length)
Diversity (cluster)
PPIFlow
BoltzGen
BindCraft
RFDiffusion
CD3d
EGFRc
EGFRn
FGFR2
H3
IGF1R
IL7Ra
InfluenzaHA
Insulin_receptor
PDGFR
Pdl1
TGF-b
Tie2
Trka
VirB8
Hotspot Coverage Comparison (75 length)
Hotspot Coverage Comparison (120 length)
Hotspot Coverage (%)
Designability Comparison (75 length)
Designability Comparison (120 length)
Designability (%)
Rosetta Interface Score (75 length)
Rosetta Interface Score (120 length)
Interface Score (REU)

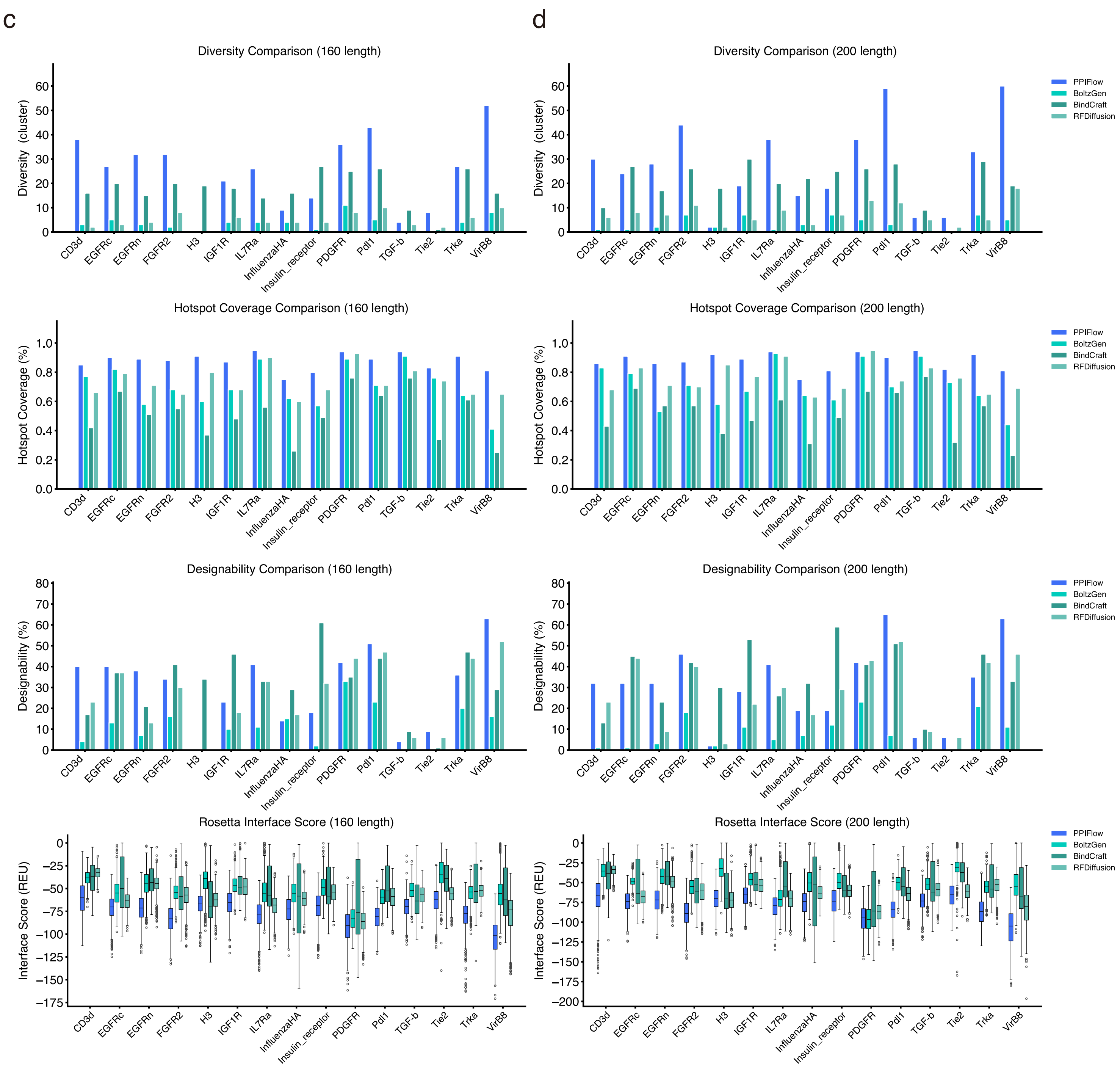
c
d
Diversity Comparison (160 length)
Diversity Comparison (200 length)
Diversity (cluster)
Hotspot Coverage Comparison (160 length)
Hotspot Coverage Comparison (200 length)
Hotspot Coverage (%)
Designability Comparison (160 length)
Designability Comparison (200 length)
Designability (%)
Rosetta Interface Score (160 length)
Rosetta Interface Score (200 length)
Interface Score (REU)
PPIFlow
BoltzGen
BindCraft
RFDiffusion
CD3d
EGFRc
EGFRn
FGFR2
H3
IGF1R
IL7Ra
InfluenzaHA
Insulin_receptor
PDGFR
Pdl1
TGF-b
Tie2
Trka
VirB8

**Figure S3: Comparative analysis of de novo binder design efficiency.**

Results are grouped by target sequence length: **(a)** 75 residues, **(b)** 120 residues, **(c)** 160 residues, and **(d)** 200 residues. For each group, four hierarchical performance metrics are evaluated (from top to bottom): *(i) Structural diversity*, quantifying the conformational breadth of the generated binder ensemble; *(ii) Hotspot coverage*, measuring the spatial conservation of critical target-interface residues; *(iii) Designability*, defined by the structural self-consistency (scRMSD) between generative backbones and forward-folded structures; *(iv) Binding energetics*, represented by Rosetta interface scores (REU). All subfigures share a common color scheme representing the four evaluated algorithms.

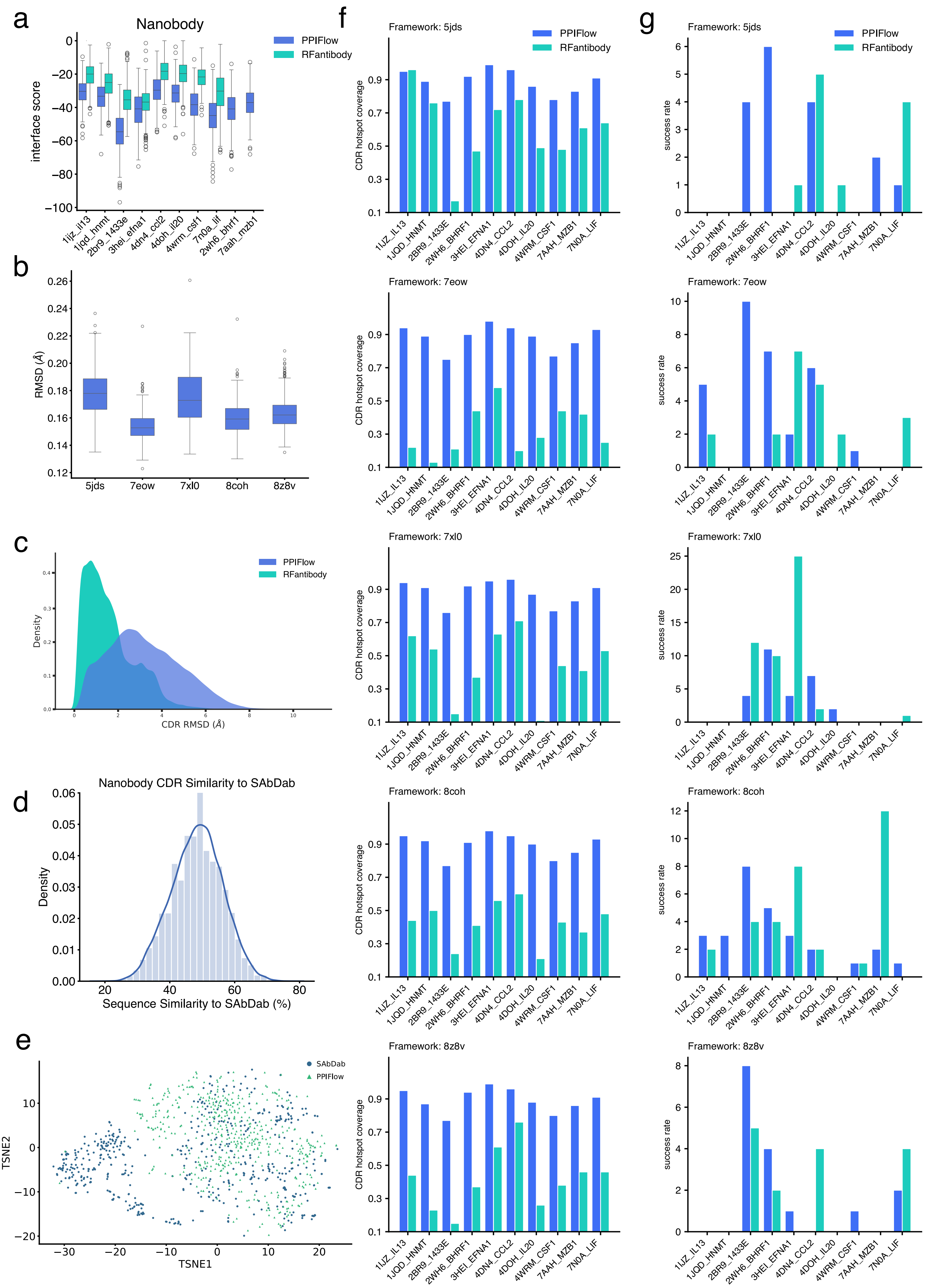
a
Nanobody
interface score
PPIFlow
RFantibody
1ijz_il13
1jqd_hnmt
2br9_1433e
3hei_efna1
4dn4_ccl2
4doh_il20
4wrm_csf1
7n0a_lif
2wh6_bhrf1
7aah_mzb1
b
RMSD (Å)
5jds
7eow
7xl0
8coh
8z8v
c
Density
CDR RMSD (Å)
PPIFlow
RFantibody
d
Nanobody CDR Similarity to SAbDab
Density
Sequence Similarity to SAbDab (%)
e
TSNE1
TSNE2
SAbDab
PPIFlow
f
Framework: 5jds
Framework: 7eow
Framework: 7xl0
Framework: 8coh
Framework: 8z8v
CDR hotspot coverage
PPIFlow
RFantibody
1IJZ_IL13
1JQD_HNMT
2BR9_1433E
2WH6_BHRF1
3HEI_EFNA1
4DN4_CCL2
4DOH_IL20
4WRM_CSF1
7AAH_MZB1
7N0A_LIF
g
Framework: 5jds
Framework: 7eow
Framework: 7xl0
Framework: 8coh
Framework: 8z8v
success rate
PPIFlow
RFantibody

**Figure S4: Benchmarking nanobody (VHH) design precision and diversity.**

**(a)** Distribution of Rosetta interface scores comparing PPIFlow and RFAntibody across nanobody design benchmarks. Notably, RFAntibody failed to generate valid binder candidates for the BHRF1 and MZB1 targets. **(b)** Structural recovery of framework regions (FRs), quantified by C$\alpha$ RMSD (Å) across multiple nanobody template PDB IDs, highlighting the backbone stability of the designs. **(c)** Conformational fidelity of CDR3 loops, calculated using the PyMOL alignment algorithm. Structures were pre-aligned via framework regions, followed by CDR3 RMSD calculation without further rotation to assess loop positioning accuracy. **(d, e)** Sequence and structural landscape analysis. (d) Sequence diversity of designed CDR3 loops relative to the SAbDab database (training set); (e) Structural clustering of the generated ensembles, illustrating the breadth of the sampled conformational space compared to known antibody repertoires. **(f, g)** Comprehensive evaluation of design quality across framework templates, reporting (f) CDR hotspot coverage efficiency and (g) design success rates across the diverse target panel.

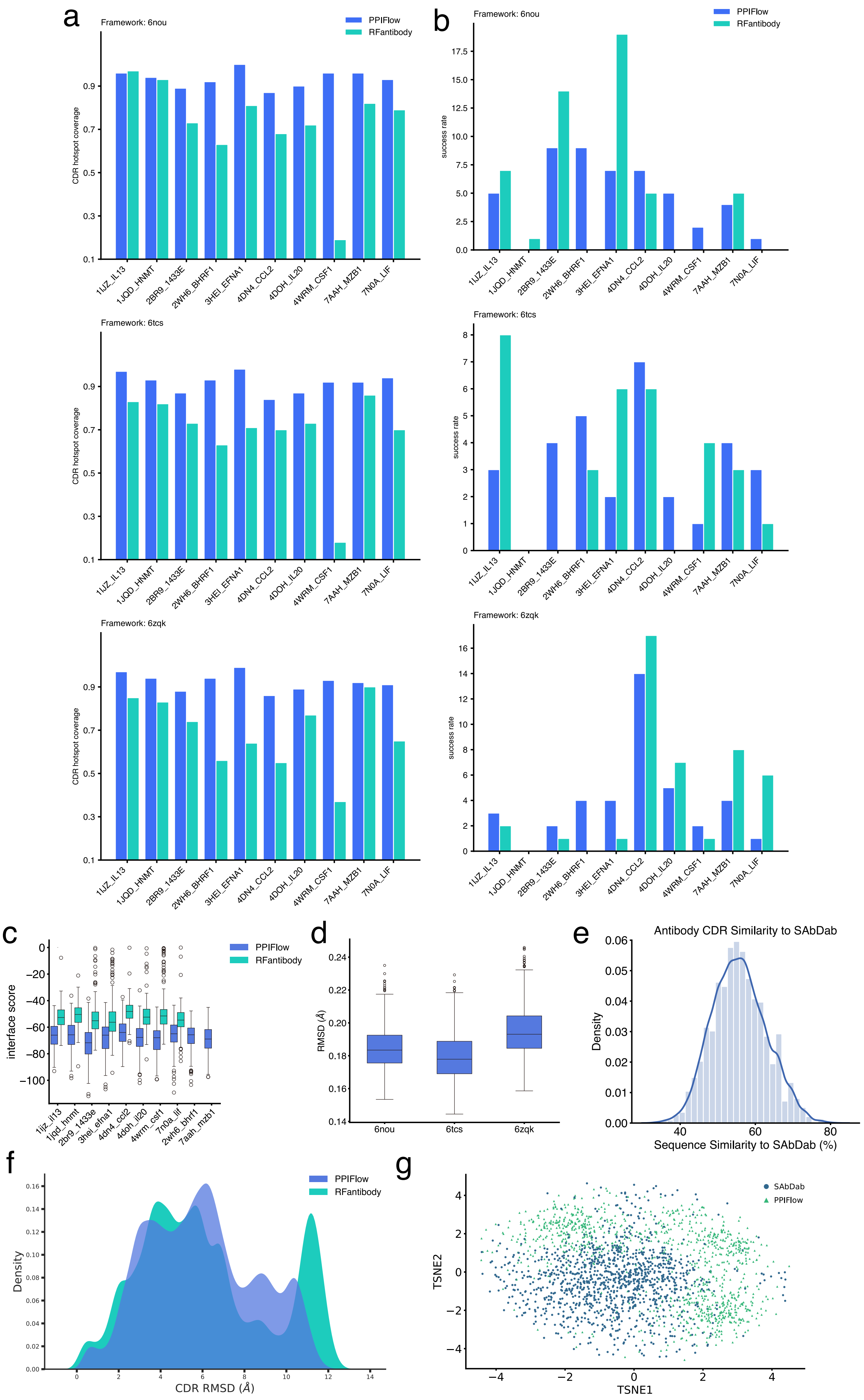
a
b
Framework: 6nou
Framework: 6tcs
Framework: 6zqk
PPIFlow
RFantibody
CDR hotspot coverage
success rate
1IJZ_IL13
1JQD_HNMT
2BR9_1433E
2WH6_BHRF1
3HEI_EFNA1
4DN4_CCL2
4DOH_IL20
4WRM_CSF1
7AAH_MZB1
7N0A_LIF
c
interface score
1ijz_il13
1jqd_hnmt
2br9_1433e
3hei_efna1
4dn4_ccl2
4doh_il20
4wrm_csf1
7n0a_lif
2wh6_bhrf1
7aah_mzb1
d
RMSD (Å)
6nou
6tcs
6zqk
e
Antibody CDR Similarity to SAbDab
Density
Sequence Similarity to SAbDab (%)
f
Density
CDR RMSD (Å)
g
SAbDab
PPIFlow
TSNE1
TSNE2

**Figure S5: Benchmarking scFv design precision.**

**(a, b)** Detailed performance metrics for antibody design tasks across a library of diverse framework templates. Panels illustrate the efficiency of PPIFlow in maintaining CDR hotspot coverage and achieving high success rates for various antibody frameworks; **(c)** Distribution of Rosetta interface scores comparing PPIFlow and RFAntibody across scFv design benchmarks. Notably, RFAntibody failed to generate valid binder candidates for the BHRF1 and MZB1 targets. **(d)** Structural recovery of framework regions (FRs), quantified by C$\alpha$ RMSD (Å) across multiple template PDB IDs, highlighting the backbone stability of the designs. **(e)** Sequence diversity of designed CDR3 loops relative to the SAbDab database (training set); **(f)** Conformational fidelity of CDR3 loops, calculated using the PyMOL alignment algorithm. Structures were pre-aligned via framework regions, followed by CDR3 RMSD calculation without further rotation to assess loop positioning accuracy. **(g)** Structural clustering of the generated ensembles, illustrating the breadth of the sampled conformational space compared to known antibody repertoires.

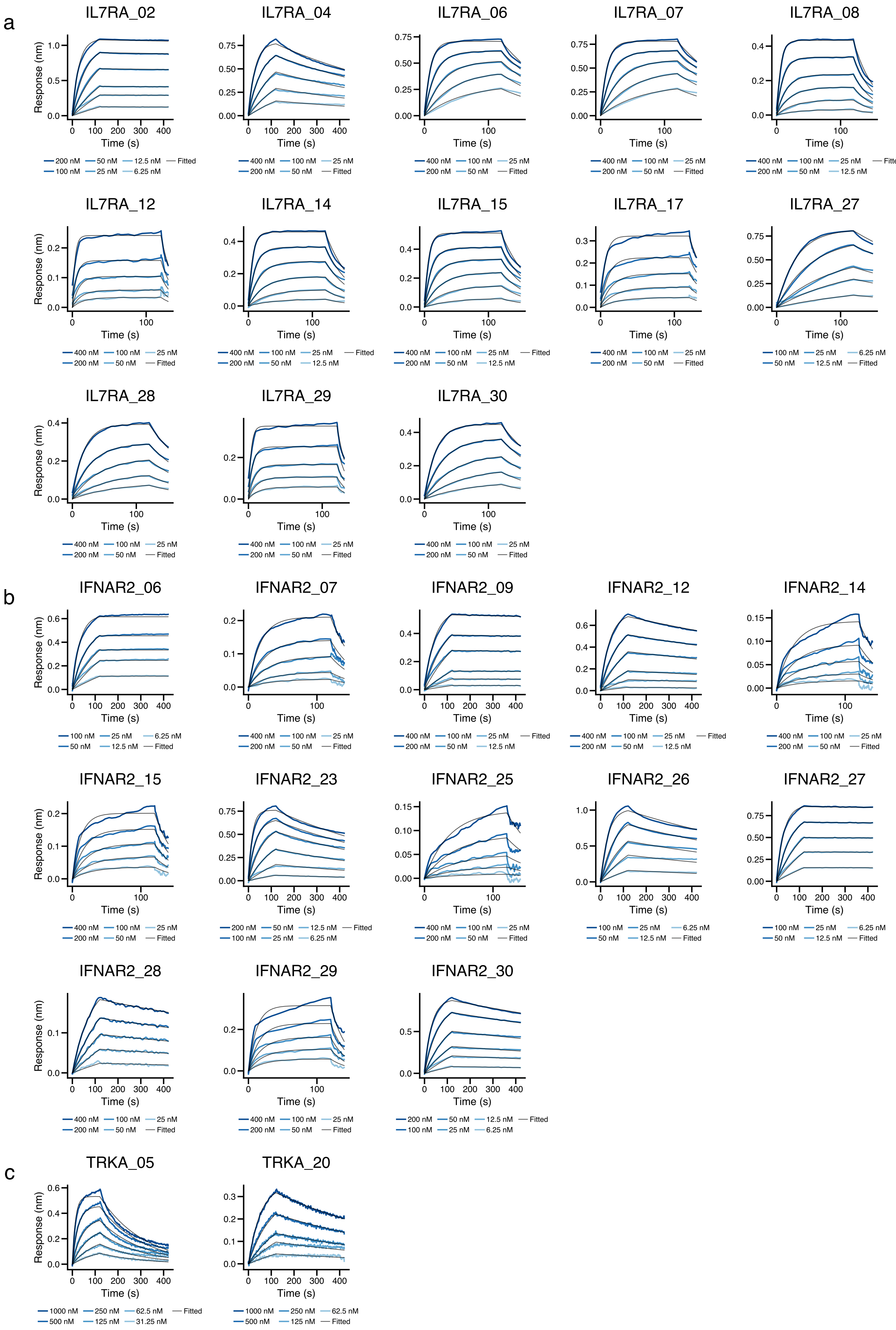

a
IL7RA_02
IL7RA_04
IL7RA_06
IL7RA_07
IL7RA_08
IL7RA_12
IL7RA_14
IL7RA_15
IL7RA_17
IL7RA_27
IL7RA_28
IL7RA_29
IL7RA_30
b
IFNAR2_06
IFNAR2_07
IFNAR2_09
IFNAR2_12
IFNAR2_14
IFNAR2_15
IFNAR2_23
IFNAR2_25
IFNAR2_26
IFNAR2_27
IFNAR2_28
IFNAR2_29
IFNAR2_30
c
TRKA_05
TRKA_20
Response (nm)
Time (s)
1000 nM
500 nM
400 nM
250 nM
200 nM
125 nM
100 nM
62.5 nM
50 nM
31.25 nM
25 nM
12.5 nM
6.25 nM
Fitted

d

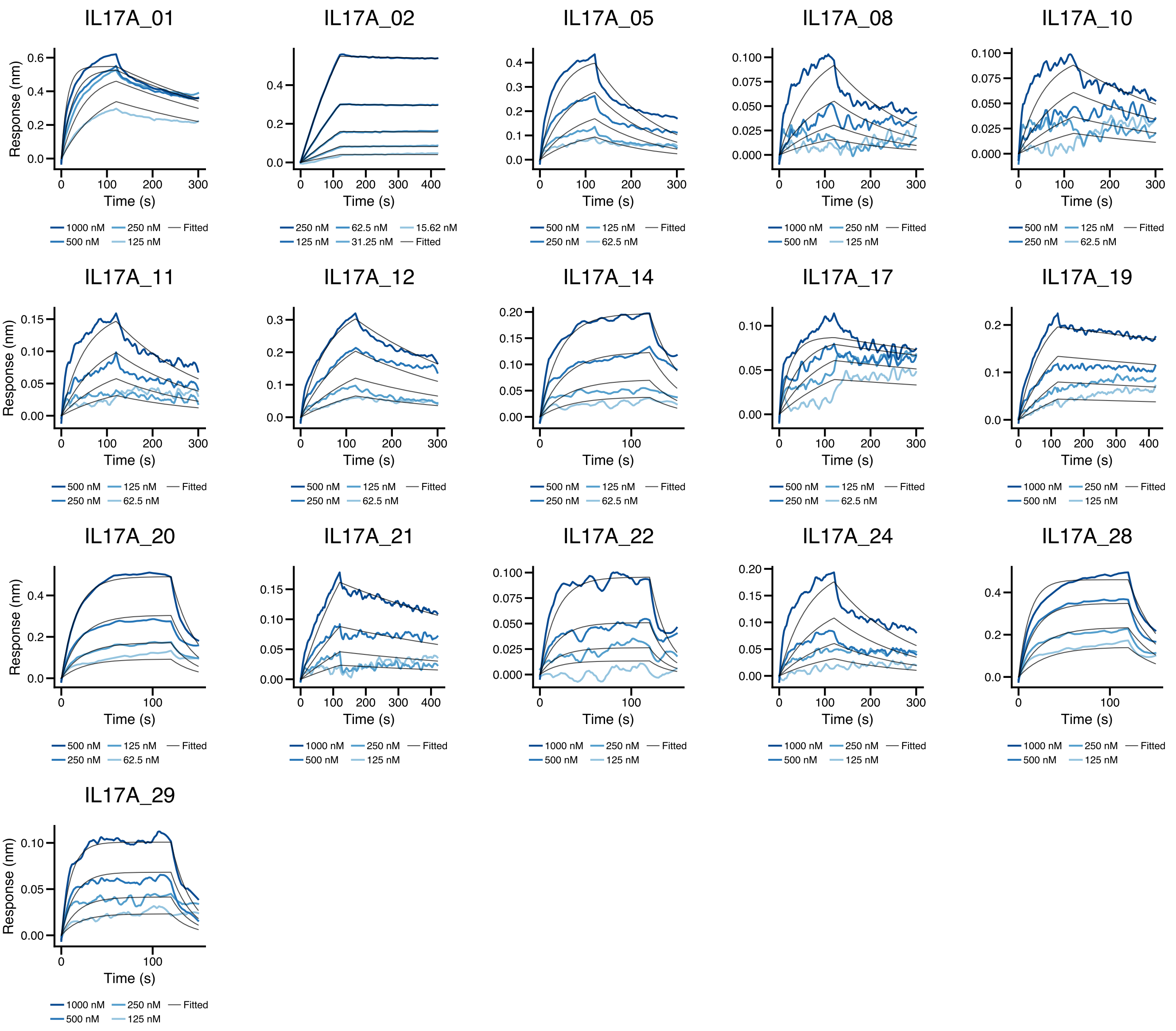
IL17A_01
Response (nm)
Time (s)
1000 nM 250 nM Fitted
500 nM 125 nM
IL17A_02
Time (s)
250 nM 62.5 nM 15.62 nM
125 nM 31.25 nM Fitted
IL17A_05
Time (s)
500 nM 125 nM Fitted
250 nM 62.5 nM
IL17A_08
Time (s)
1000 nM 250 nM Fitted
500 nM 125 nM
IL17A_10
Time (s)
500 nM 125 nM Fitted
250 nM 62.5 nM
IL17A_11
Response (nm)
Time (s)
500 nM 125 nM Fitted
250 nM 62.5 nM
IL17A_12
Time (s)
500 nM 125 nM Fitted
250 nM 62.5 nM
IL17A_14
Time (s)
500 nM 125 nM Fitted
250 nM 62.5 nM
IL17A_17
Time (s)
500 nM 125 nM Fitted
250 nM 62.5 nM
IL17A_19
Time (s)
1000 nM 250 nM Fitted
500 nM 125 nM
IL17A_20
Response (nm)
Time (s)
500 nM 125 nM Fitted
250 nM 62.5 nM
IL17A_21
Time (s)
1000 nM 250 nM Fitted
500 nM 125 nM
IL17A_22
Time (s)
1000 nM 250 nM Fitted
500 nM 125 nM
IL17A_24
Time (s)
1000 nM 250 nM Fitted
500 nM 125 nM
IL17A_28
Time (s)
1000 nM 250 nM Fitted
500 nM 125 nM
IL17A_29
Response (nm)
Time (s)
1000 nM 250 nM Fitted
500 nM 125 nM

e

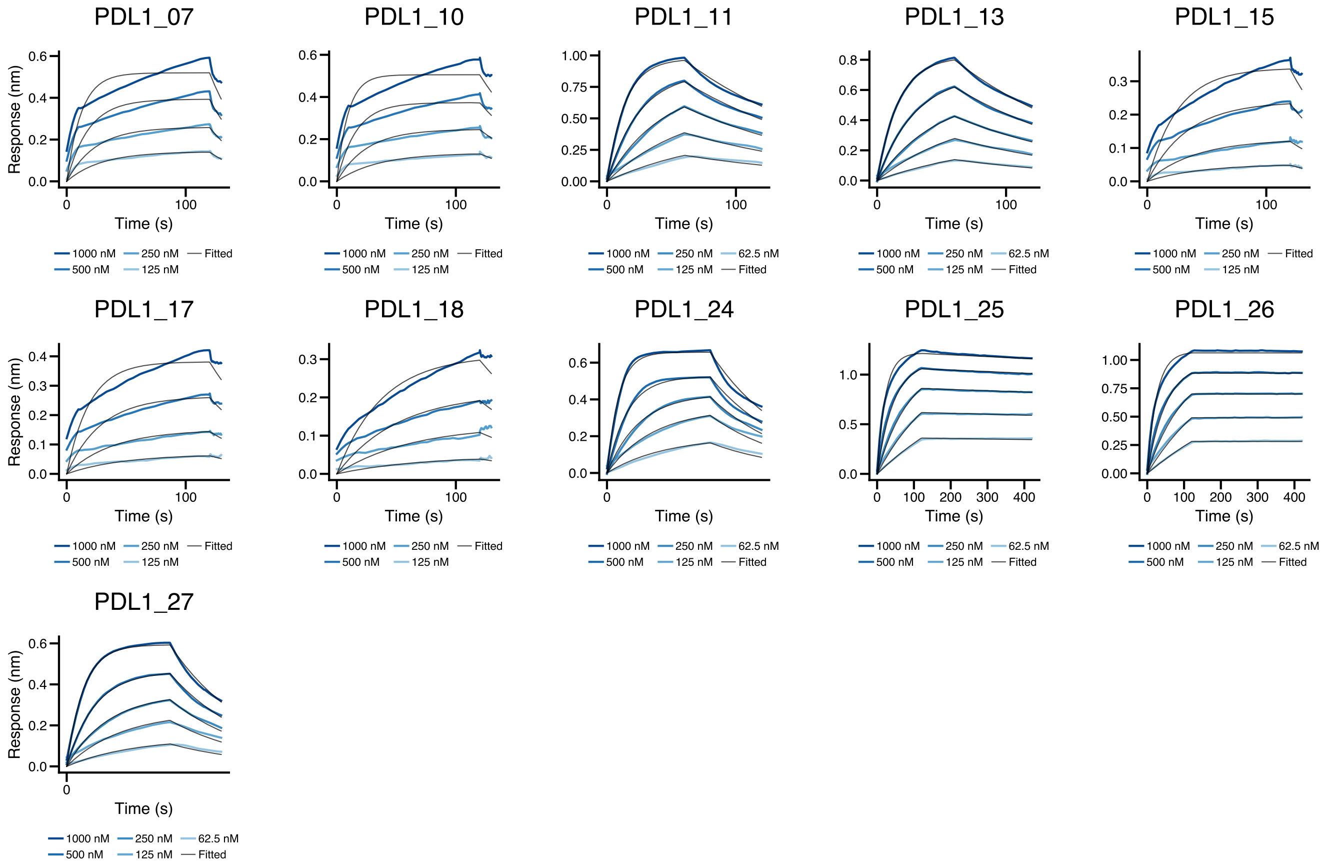
PDL1_07
Response (nm)
Time (s)
1000 nM 250 nM Fitted
500 nM 125 nM
PDL1_10
Time (s)
1000 nM 250 nM Fitted
500 nM 125 nM
PDL1_11
Time (s)
1000 nM 250 nM 62.5 nM
500 nM 125 nM Fitted
PDL1_13
Time (s)
1000 nM 250 nM 62.5 nM
500 nM 125 nM Fitted
PDL1_15
Time (s)
1000 nM 250 nM Fitted
500 nM 125 nM
PDL1_17
Response (nm)
Time (s)
1000 nM 250 nM Fitted
500 nM 125 nM
PDL1_18
Time (s)
1000 nM 250 nM Fitted
500 nM 125 nM
PDL1_24
Time (s)
1000 nM 250 nM 62.5 nM
500 nM 125 nM Fitted
PDL1_25
Time (s)
1000 nM 250 nM 62.5 nM
500 nM 125 nM Fitted
PDL1_26
Time (s)
1000 nM 250 nM 62.5 nM
500 nM 125 nM Fitted
PDL1_27
Response (nm)
Time (s)
1000 nM 250 nM 62.5 nM
500 nM 125 nM Fitted

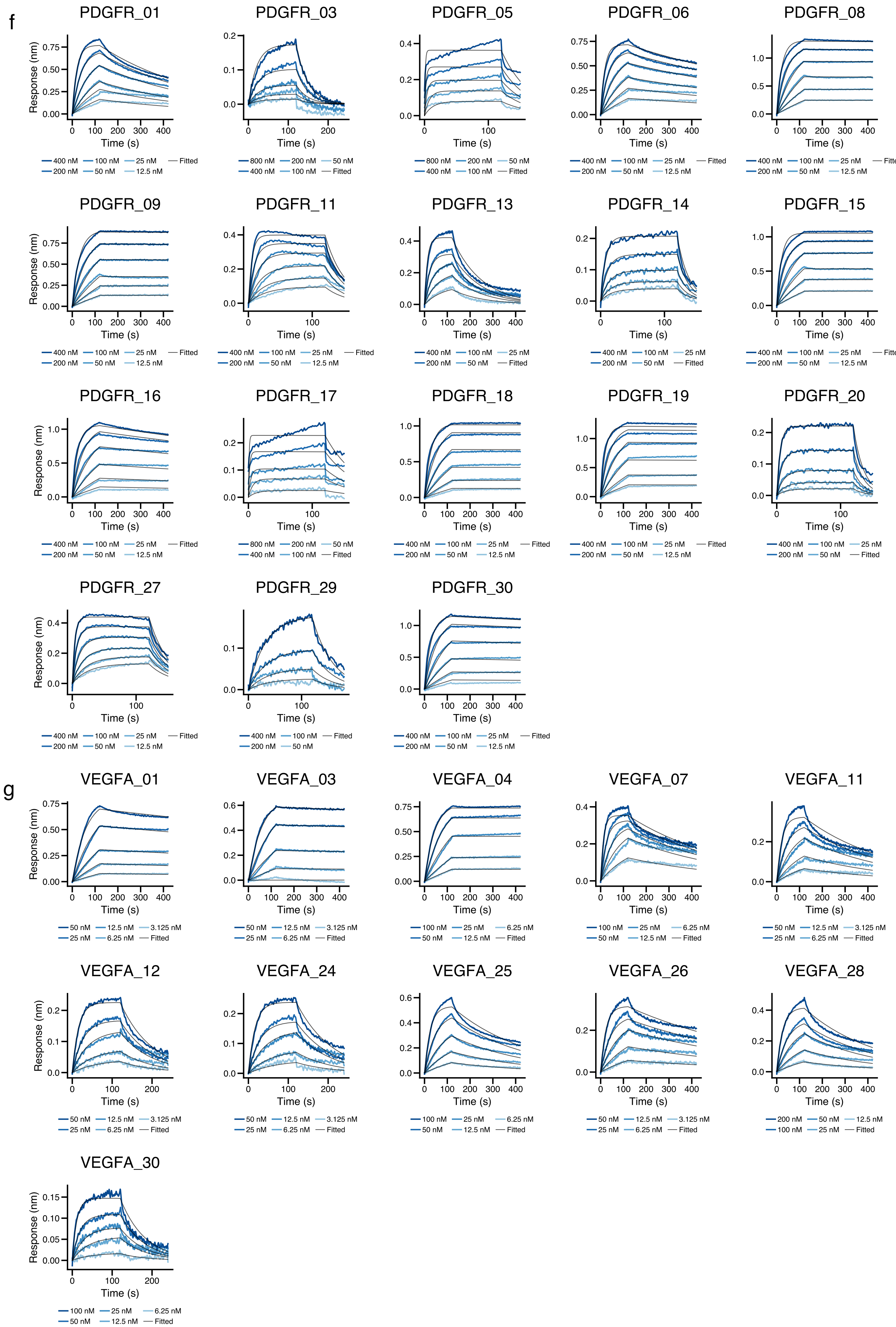
f
PDGFR_01
Response (nm)
Time (s)
400 nM 100 nM 25 nM Fitted
200 nM 50 nM 12.5 nM
PDGFR_03
Time (s)
800 nM 200 nM 50 nM
400 nM 100 nM Fitted
PDGFR_05
Time (s)
800 nM 200 nM 50 nM
400 nM 100 nM Fitted
PDGFR_06
Time (s)
400 nM 100 nM 25 nM Fitted
200 nM 50 nM 12.5 nM
PDGFR_08
Time (s)
400 nM 100 nM 25 nM Fitted
200 nM 50 nM 12.5 nM
PDGFR_09
Response (nm)
Time (s)
400 nM 100 nM 25 nM Fitted
200 nM 50 nM 12.5 nM
PDGFR_11
Time (s)
400 nM 100 nM 25 nM Fitted
200 nM 50 nM 12.5 nM
PDGFR_13
Time (s)
400 nM 100 nM 25 nM
200 nM 50 nM Fitted
PDGFR_14
Time (s)
400 nM 100 nM 25 nM
200 nM 50 nM Fitted
PDGFR_15
Time (s)
400 nM 100 nM 25 nM Fitted
200 nM 50 nM 12.5 nM
PDGFR_16
Response (nm)
Time (s)
400 nM 100 nM 25 nM Fitted
200 nM 50 nM 12.5 nM
PDGFR_17
Time (s)
800 nM 200 nM 50 nM
400 nM 100 nM Fitted
PDGFR_18
Time (s)
400 nM 100 nM 25 nM Fitted
200 nM 50 nM 12.5 nM
PDGFR_19
Time (s)
400 nM 100 nM 25 nM Fitted
200 nM 50 nM 12.5 nM
PDGFR_20
Time (s)
400 nM 100 nM 25 nM
200 nM 50 nM Fitted
PDGFR_27
Response (nm)
Time (s)
400 nM 100 nM 25 nM Fitted
200 nM 50 nM 12.5 nM
PDGFR_29
Time (s)
400 nM 100 nM Fitted
200 nM 50 nM
PDGFR_30
Time (s)
400 nM 100 nM 25 nM Fitted
200 nM 50 nM 12.5 nM
g
VEGFA_01
Response (nm)
Time (s)
50 nM 12.5 nM 3.125 nM
25 nM 6.25 nM Fitted
VEGFA_03
Time (s)
50 nM 12.5 nM 3.125 nM
25 nM 6.25 nM Fitted
VEGFA_04
Time (s)
100 nM 25 nM 6.25 nM
50 nM 12.5 nM Fitted
VEGFA_07
Time (s)
100 nM 25 nM 6.25 nM
50 nM 12.5 nM Fitted
VEGFA_11
Time (s)
50 nM 12.5 nM 3.125 nM
25 nM 6.25 nM Fitted
VEGFA_12
Response (nm)
Time (s)
50 nM 12.5 nM 3.125 nM
25 nM 6.25 nM Fitted
VEGFA_24
Time (s)
50 nM 12.5 nM 3.125 nM
25 nM 6.25 nM Fitted
VEGFA_25
Time (s)
100 nM 25 nM 6.25 nM
50 nM 12.5 nM Fitted
VEGFA_26
Time (s)
50 nM 12.5 nM 3.125 nM
25 nM 6.25 nM Fitted
VEGFA_28
Time (s)
200 nM 50 nM 12.5 nM
100 nM 25 nM Fitted
VEGFA_30
Response (nm)
Time (s)
100 nM 25 nM 6.25 nM
50 nM 12.5 nM Fitted

**Figure S6: Bio-layer interferometry (BLI) kinetic analysis of de novo designed binders.**

Multi-cycle BLI sensorgrams illustrating the binding kinetics of synthetic binders against seven therapeutic targets: **(a)** IL7RA, **(b)** IFNAR2, **(c)** TrkA, **(d)** IL17A, **(e)** PDL1, **(f)** PDGFR, and **(g)** VEGFA. Raw data (colored lines) were globally fitted with a 1:1 binding model (black lines). Molar concentrations for each titration step are annotated below the respective sensorgrams.

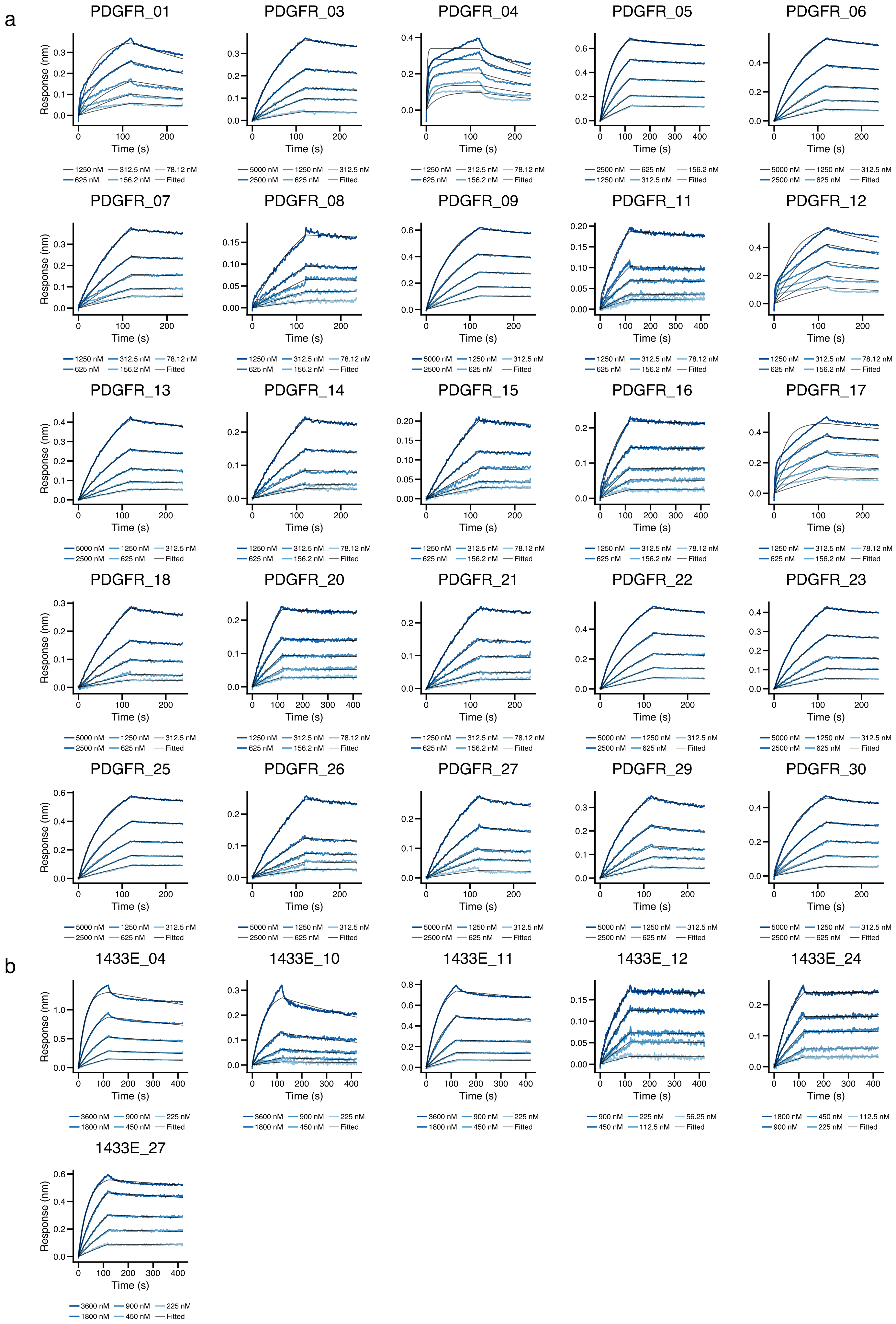
a
PDGFR_01
PDGFR_03
PDGFR_04
PDGFR_05
PDGFR_06
PDGFR_07
PDGFR_08
PDGFR_09
PDGFR_11
PDGFR_12
PDGFR_13
PDGFR_14
PDGFR_15
PDGFR_16
PDGFR_17
PDGFR_18
PDGFR_20
PDGFR_21
PDGFR_22
PDGFR_23
PDGFR_25
PDGFR_26
PDGFR_27
PDGFR_29
PDGFR_30
b
1433E_04
1433E_10
1433E_11
1433E_12
1433E_24
1433E_27
Response (nm)
Time (s)
1250 nM 312.5 nM 78.12 nM
625 nM 156.2 nM Fitted
5000 nM 1250 nM 312.5 nM
2500 nM 625 nM Fitted
2500 nM 625 nM 156.2 nM
1250 nM 312.5 nM Fitted
3600 nM 900 nM 225 nM
1800 nM 450 nM Fitted
900 nM 225 nM 56.25 nM
450 nM 112.5 nM Fitted
1800 nM 450 nM 112.5 nM
900 nM 225 nM Fitted

c

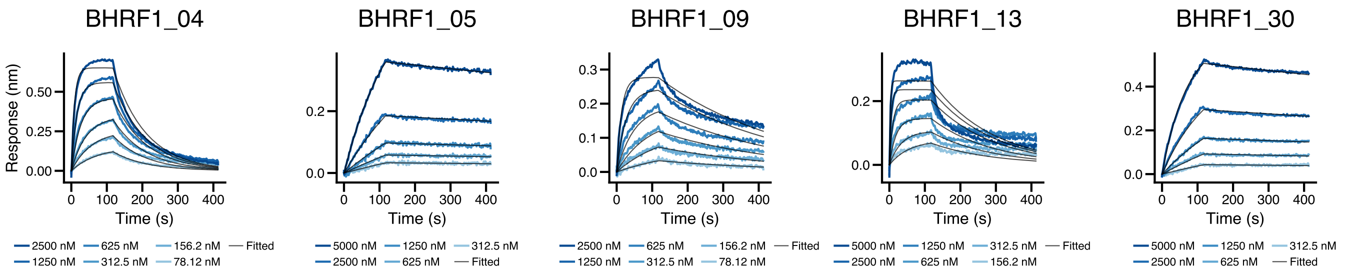

BHRF1_04
BHRF1_05
BHRF1_09
BHRF1_13
BHRF1_30
Response (nm)
Time (s)
2500 nM 625 nM 156.2 nM Fitted
1250 nM 312.5 nM 78.12 nM
5000 nM 1250 nM 312.5 nM
2500 nM 625 nM Fitted
2500 nM 625 nM 156.2 nM Fitted
1250 nM 312.5 nM 78.12 nM
5000 nM 1250 nM 312.5 nM Fitted
2500 nM 625 nM 156.2 nM
5000 nM 1250 nM 312.5 nM
2500 nM 625 nM Fitted

d

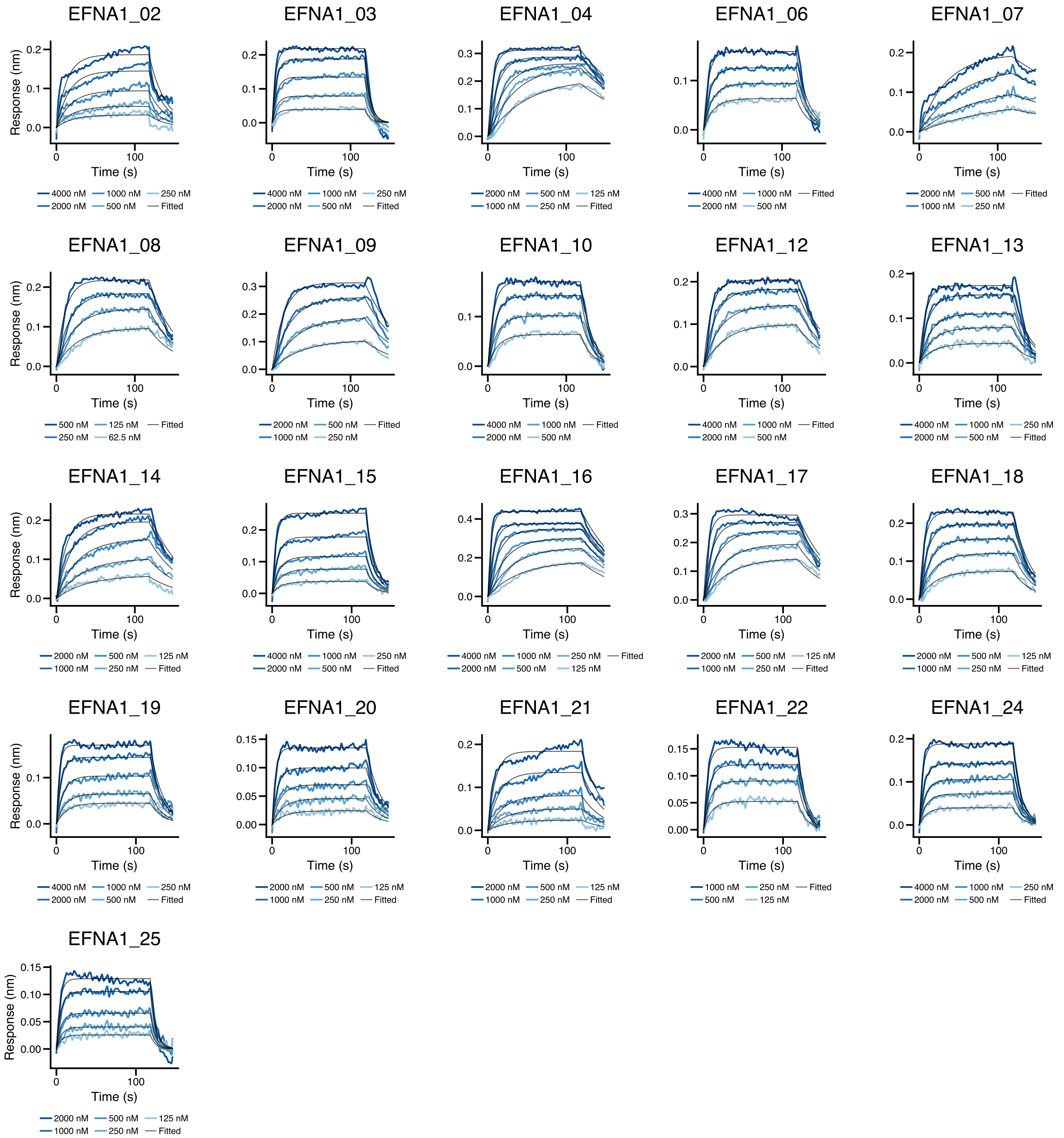

EFNA1_02
EFNA1_03
EFNA1_04
EFNA1_06
EFNA1_07
EFNA1_08
EFNA1_09
EFNA1_10
EFNA1_12
EFNA1_13
EFNA1_14
EFNA1_15
EFNA1_16
EFNA1_17
EFNA1_18
EFNA1_19
EFNA1_20
EFNA1_21
EFNA1_22
EFNA1_24
EFNA1_25
Response (nm)
Time (s)
4000 nM 1000 nM 250 nM
2000 nM 500 nM Fitted
2000 nM 500 nM 125 nM
1000 nM 250 nM Fitted
4000 nM 1000 nM Fitted
2000 nM 500 nM
2000 nM 500 nM Fitted
1000 nM 250 nM
500 nM 125 nM Fitted
250 nM 62.5 nM
1000 nM 250 nM Fitted
500 nM 125 nM

e

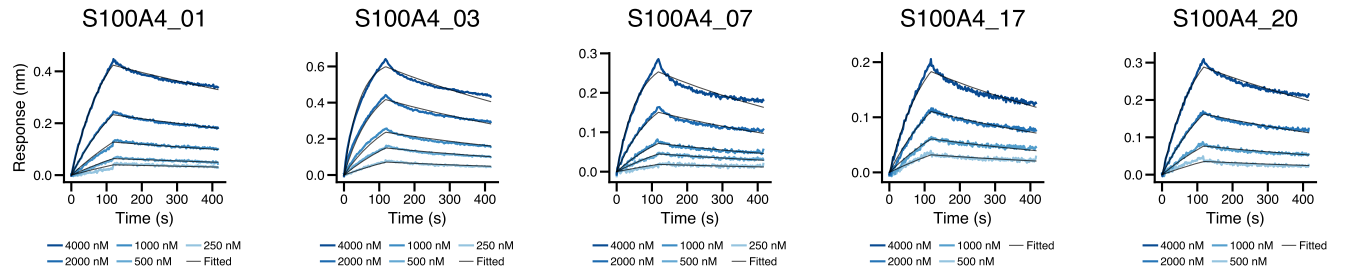

S100A4_01
S100A4_03
S100A4_07
S100A4_17
S100A4_20
Response (nm)
Time (s)
4000 nM 1000 nM 250 nM
2000 nM 500 nM Fitted
4000 nM 1000 nM Fitted
2000 nM 500 nM

f

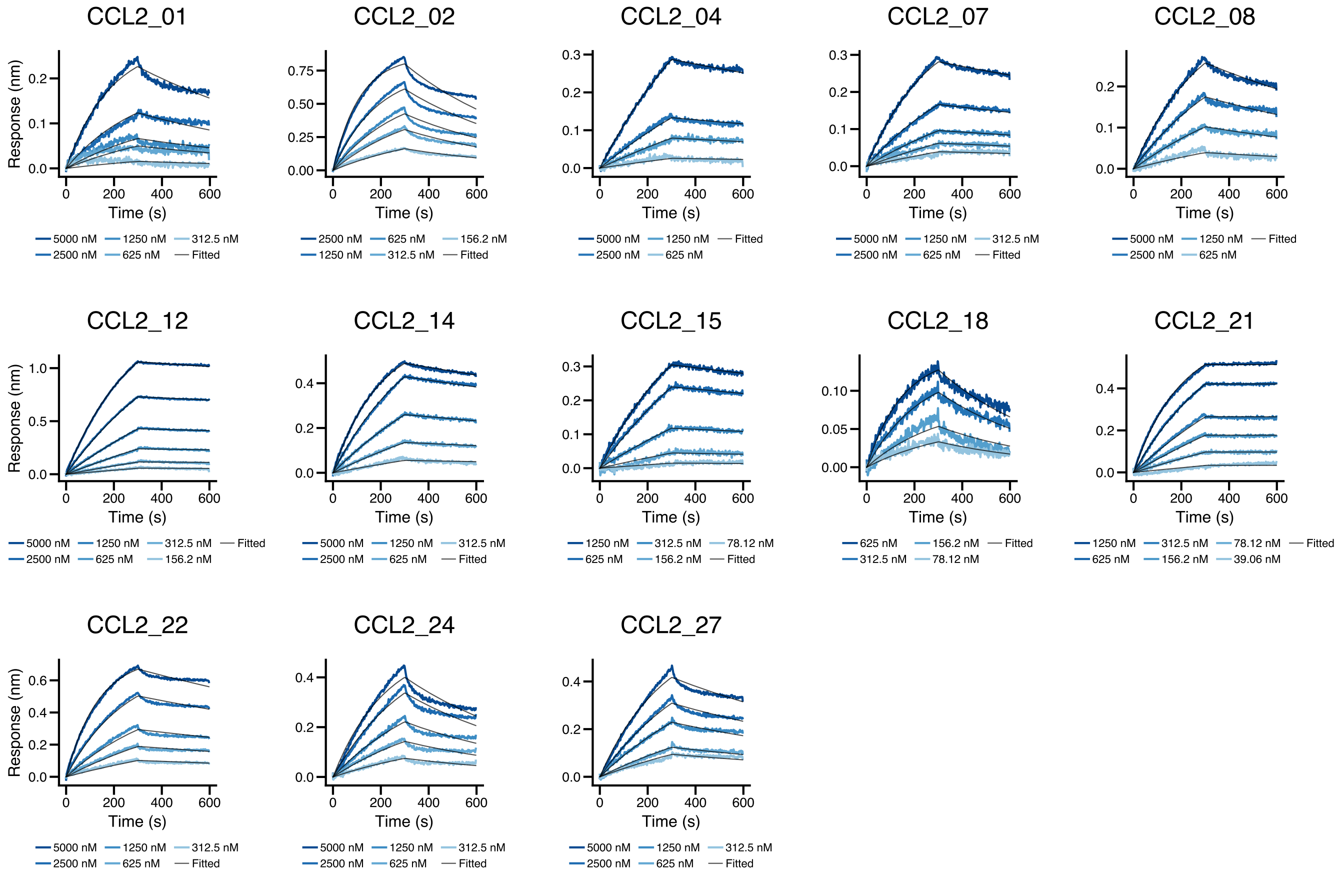
CCL2_01
CCL2_02
CCL2_04
CCL2_07
CCL2_08
CCL2_12
CCL2_14
CCL2_15
CCL2_18
CCL2_21
CCL2_22
CCL2_24
CCL2_27
Response (nm)
Time (s)
5000 nM 1250 nM 312.5 nM
2500 nM 625 nM Fitted
2500 nM 625 nM 156.2 nM
1250 nM 312.5 nM Fitted
5000 nM 1250 nM Fitted
2500 nM 625 nM
5000 nM 1250 nM 312.5 nM Fitted
2500 nM 625 nM 156.2 nM
1250 nM 312.5 nM 78.12 nM
625 nM 156.2 nM Fitted
625 nM 156.2 nM Fitted
312.5 nM 78.12 nM
1250 nM 312.5 nM 78.12 nM Fitted
625 nM 156.2 nM 39.06 nM

g

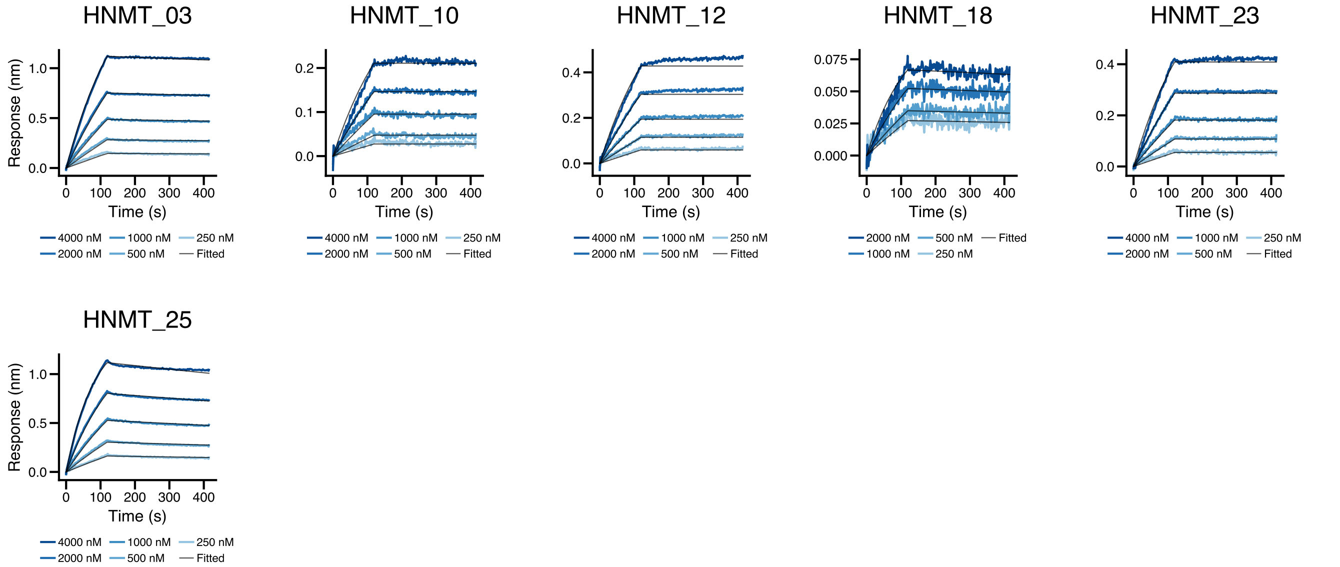
HNMT_03
HNMT_10
HNMT_12
HNMT_18
HNMT_23
HNMT_25
Response (nm)
Time (s)
4000 nM 1000 nM 250 nM
2000 nM 500 nM Fitted
2000 nM 500 nM Fitted
1000 nM 250 nM

**Figure S7: Kinetic characterization of designed VHH domains via BLI.**

Representative BLI sensorgrams showing the binding affinity of designed VHH candidates against: **(a)** PDGFR, **(b)** 1433E, **(c)** BHRF1, **(d)** EFNA1, **(e)** S100A4, **(f)** CCL2 and **(g)** HNMT. Multi-point kinetic titrations were performed to evaluate binding specificity. The data confirm high-affinity interactions and diverse dissociation profiles across the tested VHH-target complexes.

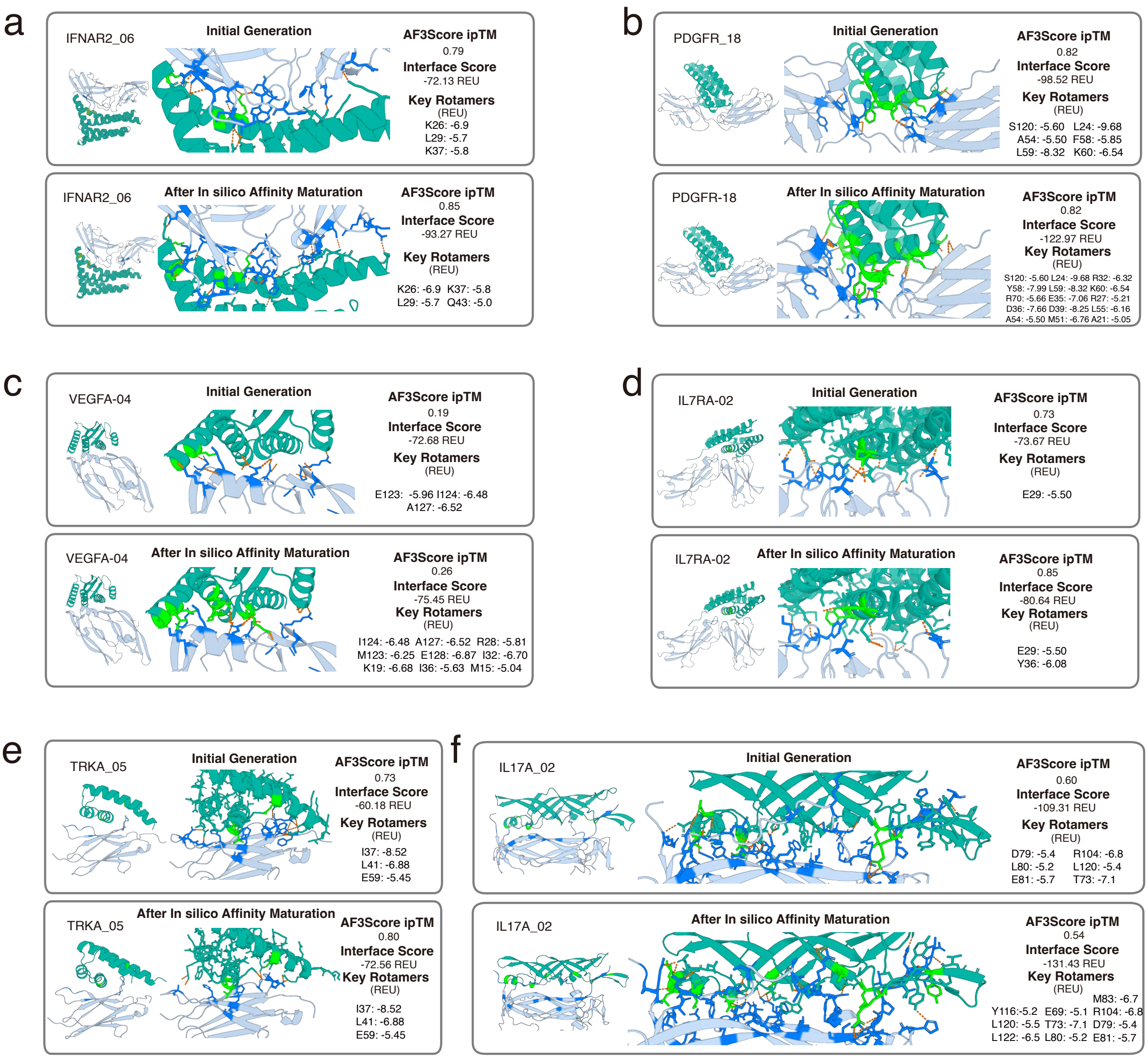

**Figure S8: Affinity maturation and structural characterization of designed binders.**

**(a–f)** Optimization of lead candidates for each target. (Note: PD-L1 is omitted here as it is presented in the main figure). Each panel displays a comparative view between the initial generative model output (top) and the variant after affinity maturation (bottom). **Structural details:** Left panels illustrate the global fold and magnified interface interactions. Key residues on the designed binders are highlighted in *green*, with target-binding residues shown in *marine*. Hydrogen bonds are represented by *orange dashed lines*. **Computational metrics:** The corresponding AF3Score confidence scores and Rosetta interface energy scores are provided to the right of each structure. The interface decomposition highlights critical residues achieving total interaction energy $< -5$ Rosetta Energy Units (REU).

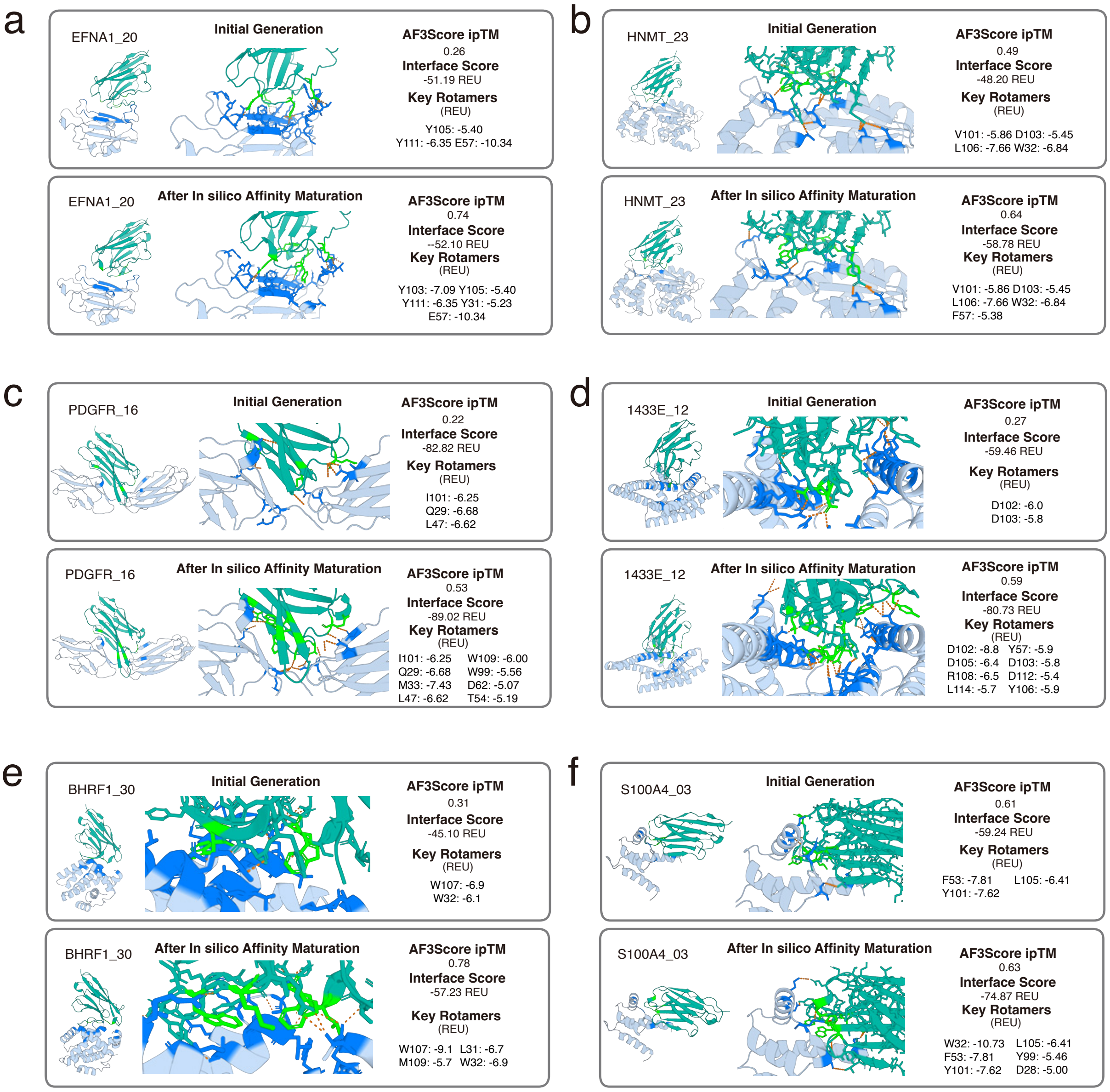

**Figure S9: Affinity maturation and structural characterization of designed VHHs.**

**(a–f)** Optimization of lead candidates for each target. (Note: CCL2 is omitted here as it is presented in the main figure). Each panel displays a comparative view between the initial generative model output (top) and the variant after affinity maturation (bottom). **Structural details:** Left panels illustrate the global fold and magnified interface interactions. Key residues on the designed binders are highlighted in *green*, with target-binding residues shown in *marine*. Hydrogen bonds are represented by *orange dashed lines*. **Computational metrics:** The corresponding AF3Score confidence scores and Rosetta interface energy scores are provided to the right of each structure. The interface decomposition highlights critical residues achieving total interaction energy $< -5$ Rosetta Energy Units (REU).

#### Appendix E Supplementary Tables

**Table S4: Binder design task specifications for *in silico* benchmarking and experimental testing.**

| Design target | PDB ID | Input | Hotspot residues | Length range |
|---|---|---|---|---|
| PDGFR | 3mjg | X124–312 | X136, X138, X186, X246, X259, X264 | 90–150 |
| IL7R$\alpha$ | 7opb | B36–231 | B50, B76, B81, B97, B103, B155, B158, B162, B183, B187, B211, B212 | 90–150 |
| IFNAR2 | 3se4* | C34–232 | C46, C48, C76, C80, C98, C100, C133, C138, C186, C189, C190 | 90–150 |
| PDL1 | 8znl | B20–133 | B50, B64, B118, B126 | 60–130 |
| IL17A | 4hsa | A17–131, B19–127 | A20, A85, A96, A99, A105, A127, B25, B26, B27 | 120–160 |
| TRKA | 1www | X282–382 | X294, X296, X333 | 60–150 |
| VEGFA | 1bj1 | V14–107, W14–107 | W81, W83, W91 | 90–150 |

* For IFNAR2 (PDB: 3se4), missing residues at the protein interface were repaired using homology modeling via SWISS-MODEL.

**Table S5: VHH design problem specifications for *in silico* benchmarking and experimental testing.**

| Design target | PDB ID | Input | Hotspot residues | CDR length range |
|---|---|---|---|---|
| CCL2 | 4dn4 | M9-69 | M28, M39, M55, M61 | 8,8,9-21 |
| PDGFR | 3mjg | X124-312 | X136, X138, X270 | 8,8,9-21 |
| 1433E | 2br9 | A3-232 | A175, A198, A218, A229 | 8,8,9-21 |
| BHRF1 | 2wh6 | A2-158 | A64, A71, A80, A87, A106 | 8,8,9-21 |
| IL13 | 1ijz | A1-113 | A11, A14, A15, A101, A107, A108 | 8,8,9-21 |
| EFNA1 | 3hei | B18-149 | B46, B90, B114 | 8,8,9-21 |

**Table S6: Design summary for protein and unique scaffolds information.**

| Target | Format | Unique Scaffolds | Hits to Wet Lab |
|---|---|---|---|
| IL17A | Binder | 30 | 30 |
| IL7R$\alpha$ | Binder | 22 | 30 |
| IFNAR2 | Binder | 26 | 30 |
| PDGFR | Binder | 23 | 30 |
| PDL1 | Binder | 26 | 30 |

| Target | Format | Unique Scaffolds | Hits to Wet Lab |
|---|---|---|---|
| TRKA | Binder | 27 | 30 |
| VEGFA | Binder | 28 | 30 |
| 1433E | VHH | 30 | 30 |
| BHRF1 | VHH | 30 | 30 |
| CLL2 | VHH | 30 | 30 |
| EFNA1 | VHH | 16 | 30 |
| HNMT | VHH | 30 | 30 |
| IL13 | VHH | 20 | 30 |
| PDGFR | VHH | 30 | 30 |
| S100A | VHH | 30 | 30 |

**Table S7: Sequences and binding affinities of all binders per target.**

| Design | $K_D$ (M) | Sequence |
|---|---|---|
| IL17A_01 | $2.49 \times 10^{-8}$ | MSGMTYLVMNPGDVDSGNVNPYLIAKYWDEIEALAKEVVANGG APVTKTLTETEKITVTFTTWEAAEPLYPTGRLPPNSTFLSTET VTVTVKTETTLTAELLWYDPELNIAMVRVTIKVKVTLLDDPSD PYVQALLQLPQEGTGEITVLVSGSGSHHWGSTHHHHHH |
| IL17A_02 | $7.47 \times 10^{-10}$ | MSGSTMHIKSSDLVGAWVSGLSTTVSVTRPDGTTVTLTVSGEA RIVSAEDATLTFDPADPTKAEGTFKATVEVENTSIDNVDLEVM EQVGGSLTANVTLKLSLDPERSYTGTDSSGNTYYSGLLLVEPV ESEPAVINGSGSHHWGSTHHHHHH |
| IL17A_05 | $1.39 \times 10^{-7}$ | MSGIVVSPSSISAAPGSSISITVTLTGVPSGSVVYVEVSLYES DFTDLYQLLDSATLNTATSTSVTFTINLSSLKSFSTELTLVDG SKVTRTRKIVARYSLMTPSVDPATGTPIYVARPGNVDGSDSTT LSEGSGSHHWGSTHHHHHH |
| IL17A_08 | $1.66 \times 10^{-7}$ | MSGGTSILVAHVKGEDAKVELLTPVVEDTLPPPGTKNLLSSKI YDEKGNEVEEYLKSKGKFMIVILLDTYEVTLIFRTKEEDADEI LKKYPTLIFEYVAYRGEKKLVEDGMDPETNMPDITEKYQWWVE RVTAKGKLKKVSPEELPALKAKLLGFGLWEGSGSHHWGSTHHH HHH |
| IL17A_10 | $2.72 \times 10^{-8}$ | MSGAAPAVTISETTTKLSTTYTANSDGTITLTVTITTPDGTTK TITFTFRPANPLASTIVSATLSDITVTLINVTITAPADNAVTV KYYKENSTKPILETETIAPGQSGTVIFPRIEGTLTLTTRGELG GEVIDEETKTIPVATYPASLELTSNGATYSVNVGSGSHHWGST HHHHHH |
| IL17A_11 | $1.32 \times 10^{-7}$ | MSGTRSLNLLTVTLTISHDVDISKGEEAEITITVTVTSADGSK ITSTLVLSSVMNRLSSGSPDIHTTVSIDEKNGTITVELVFPAE PGEYKVDATVNIKTRNFDGETSSKTEKVEAKLSLETIELEPGE SKQLSEDLYVVNPLGSGSHHWGSTHHHHHH |

Continued on next page...

Table S7 (Continued)

| **Design** | $K_D$ **(M)** | **Sequence** |
|---|---|---|
| IL17A_12 | $8.30 \times 10^{-8}$ | MSGGIDFSFLTEEDVGKWVEVESSLKASVDDYEVEIEGTFAVKVSRSYTVTSPDGSSSSTAIVVSDYSLLSGKITVKIPKEVYDAAKKWGEDHVTDVLNDLVRSMLGFIIPSDTKWEFVSTSEDGMATYEAEVKLPRPPGSGSHHWGSTHHHHHH |
| IL17A_14 | $6.38 \times 10^{-8}$ | MSGGVSEDIPEEEALALATEALTKAYNGKTVTRTFANETSDIPVDAPFKERMAVEWYLEGDDDRVVIVPVEATFEGERTLTSVDGEPLDPAEMKAIAAKAKAAGKKISLGEIANEYTKSTTIFGLPEIHSYITVEEKVKLVKVEILPEFKGKVLVKTLGSGSHHWGSTHHHHHH |
| IL17A_17 | $7.45 \times 10^{-9}$ | MSGSVTVTIEFEETIYPSNYFTGIETDDLIIELTLLSYKVKYKGSAVVENGVAVIRPEDMEPLKVLEAKVKLTVFDHFAPGKSTTISLKDENGNELVSLEVKGGEIKSKEFTFKIETDKLLDVTEEIKKYIYLDGKPLGSGSHHWGSTHHHHHH |
| IL17A_19 | $2.93 \times 10^{-8}$ | MSGSNSKIEVDKEEKIEGDKLIIELDIELEGLANESVQKLVEEIKKELGEDTNIEIYVEGKPLKWPKELPLDIMAPVNARVSIKIKIVAFLGSGSHHWGSTHHHHHH |
| IL17A_20 | $1.42 \times 10^{-8}$ | MSGKKTYMFAIAKVTFTFFDDASIFSDATATVTVYKDGKNITGDPNLTVSMSYDYTLSLSKTVTVLVRYVEEVKDENTKYTVNVTVTATFTPLPDPEFRAALTSNVLPATTTKTSTVEAKKIYVTIPEGYSLSISYDPYHDTVSVEISPGSGSHHWGSTHHHHHH |
| IL17A_21 | $3.82 \times 10^{-7}$ | MSGMDKALIVLLKGKLKEPLEIMKKNNLSILEMEELIVKRIKEEGLSSLNMEIVDVYRLEDGKLTKLEEEKITMEEVDKMKGVIAIDLRYTTTPKPETLNEDGIPDWTSKPYGDREITDIELYAAVVTTPEGEPLKEYAKELAIYLVRKYAEESGSGSHHWGSTHHHHHH |
| IL17A_22 | $4.90 \times 10^{-6}$ | MSGGSTSYVLNDPVVSIKGVPEEEVEKIEVDEEKTITLKPNMNISDLVEELLKLIKEMEKKLLEKVQANLESLKVGEVVELKTLIKLTPAGGKPVTISANSFMGPDLASRLANDLINVFLNDFGAKKSADGFSITLPAKPLYLLLTVSISISSRGSGSHHWGSTHHHHHH |
| IL17A_24 | $6.20 \times 10^{-7}$ | MSGMVTVSVTVDLTEKGTLVSRTETTGATTTSLDTSLDNRDRASDPVVTSISIVDVETGSSEGIEIESITIMELPEPQADTNWRYLVTLSVKLPEGSSFSKTLNITLTTSYNKTTVTTSNRPGVNDDFVTMSEVNTQTSTVTETWSATVNVTFKPPGSGSHHWGSTHHHHHH |
| IL17A_27 | $1.09 \times 10^{-7}$ | MSGATETIVLTEDDITVTANEKSLTVTPDTTSLTFTATLTIKALPHDLGEITVTFSLAYFQSQGLIVSGAPVTVTVKAGETVTVTRTLTVPLSKLKNIVQRLKWGWEGEEQLVIAAELTSENLDGGKAPRVYSDPITIKVVLPEGSGSHHWGSTHHHHHH |
| IL17A_28 | $8.48 \times 10^{-8}$ | MSGMKTTYITVPLGETTTITVTLTADQHPSMGPTVTVTVKIAVNTFYYSTPAGFDQTLQSQTVARGSTTTITISLNNKELAGDRMLDGFAEITVLVYYETHVPIAPPFTVNGKTYYPMESVLSGEKVIIGVREGSGSHHWGSTHHHHHH |
| IL17A_29 | $8.41 \times 10^{-7}$ | MSGMTVTLPTLPKLTLINSRGDDPNSVSLNIRENGKFKLYSVNFGESKTVTTTLLDANGNYVPVTLTLTLSEPKTVKPRTLLAEFEEAGLREWEVVKESTLTISVTVLATGASASTSTPVTVIEYSTEVGSGSHHWGSTHHHHHH |

Continued on next page...

Table S7 (Continued)

| Design | $K_D$ (M) | Sequence |
|---|---|---|
| IL7RA_02 | $3.16 \times 10^{-10}$ | MSGALKPPTPAEQLKDIAAAIKEAAAKGELIEMTQLLSFTLLVASKIFGLDPELAAEMLRLLYLAAISENPEPIVKKAIELLNKLAEEAAKVDPETAAILKEAAERLAENPENVEPVLAELGIALAEHLAKHPKVKEMGSGSHHWGSTHHHHHH |
| IL7RA_04 | $1.73 \times 10^{-8}$ | MSGKIKISEELMASISVMIGIIAQELREAGKEVDPELAKKLEKLAEEMEKLMEEIGKLMQDPNAKEEEIKEKVKKLVKVLEEAAKLAKELGKPELAEAIKELIKIIKEVVEDPSKQEEALKRVEEIINKIQDEIIKKRGSGSHHWGSTHHHHHH |
| IL7RA_06 | $4.35 \times 10^{-8}$ | MSGKEKELEEIDKLVEEVIELMEKVAKALGISEERKEEIEKEIKEKGAEEVFVELAESGPLEEALYAAALALKETKNLPIEERLKKIARIMAAISSYLEYIIENKGEEIPNLELLKELLELTNELLDKIQELEKKGSGSHHWGSTHHHHHH |
| IL7RA_07 | $4.41 \times 10^{-8}$ | MSGMSEEIKEKIDKLKAEGNEDEIIKLTYSILSLLVGLFEDVGAEAVVKGLYDDLFFGDNKEEAEEVIKEFLGLVENAAYGIKLIFPEEAETMEEIVKLAKEAVEYLKEGKVEEAVKKLNEIYELLLEPLKKLGELLEKKLSGSGSHHWGSTHHHHHH |
| IL7RA_08 | $8.85 \times 10^{-8}$ | MSGDNITKEEFDEEIIKKLKELAEGKLEFENVDEAYAFLALVARIIAKKTGKEEEIGKILSRAYIGMKYAEDESTLEKALEIAKEFAEEVEKMRKEGIFNAEYIVLAFAAYFIAKFLEKKGVKEAAEIRERIKEAVKEGAEKVIKIIKEGSGSHHWGSTHHHHHH |
| IL7RA_12 | $8.65 \times 10^{-8}$ | MSGEELLKELAEEIKRLLEDTEKLEEKLKAQGKPEEGKEAARRLLISFSLRMINRYLQKKPELKEIAKLIGKAILLLDEGKLEEAIKELEKAVELLKKLEGLEKRFGEIVELIAKILKAELEGNEKEAEKLYKKLKELLEKLAEELEKGSGSHHWGSTHHHHHH |
| IL7RA_14 | $6.98 \times 10^{-8}$ | MSGDWRAEQLARLAELEAELAKLMKEADSKEVMEVLEKAHALLSLAQSLLEEDEDVSLSLFIYSILVLSKWPELKKEIYEEVAKKSPELLKKLEEIEKLIEEQKKNPENLEIVKELKKLLKELRADEEAVRALLEVLLEHLERLLAEARKLLGSGSHHWGSTHHHHHH |
| IL7RA_15 | $6.05 \times 10^{-8}$ | MSGDITEEELKKAEIEVAAILIAITIVVLIFAGMYTKELVEKLLSTPVEEVKKLLEELEDLIEEALEKIYSRKDEKIGPIKEEILAKLKEAIAIAKALKPKSGELATLVAERLKAAAEVVKKMPDDATITEALSKAIDAIRGHGSGSHHWGSTHHHHHH |
| IL7RA_17 | $1.16 \times 10^{-7}$ | MSGDELFKEVNKEMKELLEKGKPELAEIIEEAMFLLDVSKKLAKKAKIVLETKDEELKKDLLEALKEFVERLKKFVEKVKKNIEELSEATAKLIALMAKELKKTAKEIEKVDPELAKEIKKLAEELEAILKSNPKHKKVVEELEGSGSHHWGSTHHHHHH |
| IL7RA_27 | $1.77 \times 10^{-8}$ | MSGSGPTEEEYLEELLKEYEELKKKFEENPESMTLDDIVTLSAIARFLLSKGKKEEAMEVLAYAVKMAVAQGKYRTALALAAEIAKVAGDEELAKKIEEVFEKIETLDKETAKKLAAKALKEAAKALKKKGFEKTAKILKEAAKELEKGSGSHHWGSTHHHHHH |
| IL7RA_28 | $1.15 \times 10^{-7}$ | MSGDLIEEAKKIEDETLKKIVEIMLTIDPSLAELPEEEKKLIARAFVYLLYRHEEKLKYLDDKEKLLEILEELLKDAKNGAKTSLDEKVKKALDKLAEILEKLIKLAKEGDTSYFKKIVNTIEEGEKQIQELLKSASGSGSHHWGSTHHHHHH |

Continued on next page...

Table S7 (Continued)

| Design | $K_D$ (M) | Sequence |
|---|---|---|
| IL7RA_29 | $1.12 \times 10^{-7}$ | MSGSGPTLEEYLEELKKEYEALKKKFAEDPASMTDDDIVTLSA IARALLSEGKKEEAMLVLADAAAMAAAQGKYRTLLQLVAALAE AAGDKELAEEIKKAFEEAETLSPEEAAKLAAKMVEKAAKALKE KGYEEAAKRLKEIAKRLKAGSGSHHWGSTHHHHHH |
| IL7RA_30 | $9.74 \times 10^{-8}$ | MSGLEEVAEEVKKLIEEIIKLLEELLSKLEIPEEDLAELEEML KKKSPEEVFVETAEGGPIGVTILAAIEALKSVLKLPVEEFLEK AARFMAALSNYFEERKEEIVKKSGNEELVKKIIEKAEKALDLI QSIKLAGSGSHHWGSTHHHHHH |
| IFNAR2_06 | $1.38 \times 10^{-12}$ | MSGSEEKKKRENLRKAVNEVHRAVAKPLKESLNELISPLKTRE EVQQAIKEIHEIRKYLLENAEKYPLFKEAMKKAEEAKKKLEEF KENPFDEEAREEMLKKLKEAVKLAEKFEDLLERTAVLLIIAEK LKEVVEELSKKAREGSGSHHWGSTHHHHHH |
| IFNAR2_07 | $3.52 \times 10^{-7}$ | MSGSAEIDANAARLEALLRSDPELAKAVNNLIHRKLGEKIKEI FEYLSKGEPEKAAELVMEMVDELIEELADPSLSEREAVVILLV LAIIYLRFGVAESAALLLDAVEAHRADPDAAIDRAAIVAALRA RRAEDRERILKNIEEVLKKLGSGSHHWGSTHHHHHH |
| IFNAR2_09 | $1.37 \times 10^{-9}$ | MSGSEEEEKKRKLLRNLKYRIAKEFKPIVDKIVEILKESLHAG TFKEALSDGSVFVPYKDELIAIIDKAIASLKAMRDELESIDPE LAKKLGALIDKLQALRDAIKADPANFNYLEAFSALFREIQELL EEISKEIRGSGSHHWGSTHHHHHH |
| IFNAR2_12 | $1.13 \times 10^{-8}$ | MSGGNHATLERYVGEVLKAAIDAARYARETGDLDAALARLVER AAEELAKLFRALGYEDTAAELGRRLAELLAELLRVVHDPSATQ EEADAIAAEALALAREATADAPEDDREALLRLLAQLLESLKNL WEVLDKVDAYLSGSGSHHWGSTHHHHHH |
| IFNAR2_14 | $7.10 \times 10^{-7}$ | MSGMLEINAELSYLAAKAGRPYREEGAEVGMLKYKGDPSWKEK HAKVAAKLAAGRVEAAAKVVEKYFAEEWPEILKKLKEVAELTK KAVELAEKGEKKEAQALLKEAAKKLKEILKELPEEWKTLKEIV KLSIELLKTAAEGGSGSHHWGSTHHHHHH |
| IFNAR2_15 | $2.52 \times 10^{-7}$ | MSGMAIRLERLKQLLETNKEAFLKRLKRMKEDAEKMEKIARTL LKLLKEYAEEHPEKKEELEKVLKKAEEIEKKIEELVKELKKLL EEAIKEVKEGNIEEAKKLLEKIVEIEDEILKLLREMRAMVEAL VGEDALSKRIIERNKAGITGSGSHHWGSTHHHHHH |
| IFNAR2_23 | $8.34 \times 10^{-9}$ | MSGSVREAIRAIRRAVEAILAAVQLAIAEKAGLIDAAEAAAKL AKAFAIIKASLEKIKPFMDKETYEKINKLVAKMEKAIAAGKLD EAIKAAIEILELLKKLAEKFKKIPEVKEEAERAIKEIEEAIKE LKYLSENWEEIMKKIKEKIDSLGSGSHHWGSTHHHHHH |
| IFNAR2_25 | $7.95 \times 10^{-7}$ | MSGSELEKRRRVTLAANIGALLAMLEQIEELLKEDPSLKNNPA VQKLFKDTTEKLEKLLKELSKLYPELKPLLAELAEQIKKALED PENREEYMAKAKELAAKILEKLKEIDPEAAAAFEKAYETYNNK EAVKKAIEILSGSGSHHWGSTHHHHHH |
| IFNAR2_26 | $3.48 \times 10^{-9}$ | MSGMTPVQKAARAAVLKLLKKIKDAMNKISNLPALKGMDNKER SEVLRLLNVKEAIAAITEIPDEELDALGELLLAAGFPAELVER LRELVARLKALADNPYLKSWTQDASKEELKKLLKEKEKAKEAE EAFQELLRLLGEVLAAFERKGSGSHHWGSTHHHHHH |

Continued on next page...

Table S7 (Continued)

| Design | $K_D$ (M) | Sequence |
|---|---|---|
| IFNAR2_27 | $1.22 \times 10^{-10}$ | MSGGPYANTERKLAKIRKVILDAIYKAIKTGNVEEEMAKLAEK VAALLEELLAPYLAPEEAKALAKEIAKEIADLILKYFKGEITK EQLAAELDAIAARILAAFASAAAGEQALIARLLAIMRSEIVNA KEVFEKVLKRLSGSGSHHWGSTHHHHHH |
| IFNAR2_28 | $2.12 \times 10^{-8}$ | MSGSAIKRERRKAITEEAAKVMKEVKAYIDALLKEAGKSKSEV LLEMIEKLKTEDFKAKLLEAAEIAKEAVKEGVAAIKAAIAADP KDIATVKKLAKEVLKKAAEVGKELAAKETDVATKVAIITTVTT TAIKLVAEAVAAGSGSHHWGSTHHHHHH |
| IFNAR2_29 | $3.53 \times 10^{-7}$ | MSGSELGRRVRQIARLVVALASAMLLGDLDRAEEILRRLGKLV PGAREAFEAAVARTRALRAAGLPAAEILRAAAAAIREALEVAE REIVERLRAAGFPEEAIRLLRELFEAVREALRALADEGDAERA RELLRRAIELAEELDRLLRGSGSHHWGSTHHHHHH |
| IFNAR2_30 | $3.86 \times 10^{-9}$ | MSGMELLKEAMARIAAALEDPELAALIREMAEHPGVVKMLFSR ARKLLEERGFLTPEDLLEILDLIGEVLQKVAEREGRKLPELEL VERLRATLRALPEEERARLVELLVEEIKKIVEKLEGKKRGSGS HHWGSTHHHHHH |
| PDGFR_01 | $1.14 \times 10^{-8}$ | MSGMEEARKRAAELISRKTALLMLAISVAGLLTGVSRQELWDA ARAPLSPEELRAWAEALGKLSDEFIEKLLELLTSKDLRKQAAL ILELIRELAELLGDLREAALFLIRLLDEVSTLTARFLQEGESL EEAVRKGIEELREELLAQVGSGSHHWGSTHHHHHH |
| PDGFR_03 | $2.17 \times 10^{-6}$ | MSGGRFADIDKVIKGLMGLFGIDYEEFKKNLEEGKLKEAAEQI KKLAKKMVELSEDLSKLRLMLLTAMALTVLLQKLVEEAKKSKD LEEKEYVLKTVEEIIKMIEEIIEKLEKHVKENPEDEESKEAAE FLRDILEWIKKMLEELKKTLGSGSHHWGSTHHHHHH |
| PDGFR_05 | $1.60 \times 10^{-8}$ | MSGGGNRMSPFVKEVMKWIYEKFMEILDKKLKGEDIDEDVEEF VEEIIELLRETRGKEVEVSFGIALGLVLAIERYKKKHPGSEKV ITEIMRKIGEALARRVAAEGFEEEGREIAELIEKGGEIAATGD FGEAMKLFRKAFLLLAQVLARSGSGSHHWGSTHHHHHH |
| PDGFR_06 | $4.85 \times 10^{-9}$ | MSGMPLIERIDEELERAKKDPEGVAEEEERKAVGAFYTLKLSA KNDKKFVSPELIRIELIEITFIAKELYEKTGDFEKSVEEAKKI AKEKLEKKLEKIKEGKLKEVLDELLELAKKYVEELNEELAELL KKVYKYLKEKFLSGSGSHHWGSTHHHHHH |
| PDGFR_08 | $1.13 \times 10^{-10}$ | MSGRFSTVAGVVRTKDGYSFMIQFPVHLIDEEIGEKIAEEVKK VLGKYDEKTAEYIAKLIIEAGKEKDYEKKLKLMSKALMMLKVF LGSDMPSAKAFEAAYEILEAAVKIFLERGLPEEEVAGLRKSAE EAKKLKEEVEKLGKGSGSHHWGSTHHHHHH |
| PDGFR_09 | $6.35 \times 10^{-11}$ | MSGMFEEIKKLVKEIFEKIQSNTAAQLAFRSVLLRLREDLDDD ATEWLIRLFGMLAALIKYLKISEELAEKLRELVKMLEEVEKKV KEKLSDEEKKKKYEELQKRIQEELRELVERVEQPNISLNTEVI PEFEKLVLEYIEGYKEIISGSGSHHWGSTHHHHHH |
| PDGFR_11 | $5.13 \times 10^{-8}$ | MSGSNLTLAKIALARAAKLLRKGLEEGKLEPYLERAAALLFLA YKYSGNNLIAGIFALLLRALASPEDLRGQEAMLKKLEKDGAPK ELVEALRELVEEISKTSDEEKRRELVERLIELLERAVKGEVDL EEITEEIKEMIEFLKGSGSHHWGSTHHHHHH |

Continued on next page...

Table S7 (Continued)

| Design | $K_D$ (M) | Sequence |
|---|---|---|
| PDGFR_13 | $3.74 \times 10^{-8}$ | MSGTVAAGRVLNVATAAAFAVALSLGEISEEEAEELISETNHI ELLKKVFKILTKALEKIGEEERKEIISLIFDLVEEARTLPKDE LIKKLKEREEELKKKLEELKKKGADEKEIHKEEIKYNIYKAAL IAVQNNDTQKTLREILKYYKEGSGSHHWGSTHHHHHH |
| PDGFR_14 | $2.53 \times 10^{-7}$ | MSGSMAKKNLVKMAMAASFVSGKPIALIPGKEFEISIELIMEK LKEVTDLDEAEEIIEMFFLIIAIAIKGRSEEERKKLIELAAKK LAETLNELVEKLKDPEGVARAVVEALERLSKMMKKEDVEYLRK LIIENLSEELAKLLEKGGSGSHHWGSTHHHHHH |
| PDGFR_15 | $< 1.00 \times 10^{-12}$ | MSGRFSTMAGVVRLKDGTAVMLSFNVELLASEVGAEIAEIVKE ILEPVNPKVGEYLKELILKMQKEKDYEEQVKLVSKFLMMLKVL LGSDMDTEEALEILIEIMRRIIEAALAAGYPEELTAGMRKTAE EAEKLLEERRARGEGSGSHHWGSTHHHHHH |
| PDGFR_16 | $2.86 \times 10^{-9}$ | MSGSMKKEIDKLMAQLTGVVLGFLRKLGMSRKLRSLVLDGAQK LSEEVKEFIKEQLKLKDKVEAVRKLIKLAEELSKEPGKEKEVI FILALAFLLAYLAGDKELAREAVERLAEELKKSMPDKEEGERI AEELIARVDELLSGSGSHHWGSTHHHHHH |
| PDGFR_17 | $6.48 \times 10^{-9}$ | MSGSKLQKIKQLEQLEVKQMQRLLERMKKLWEETGDKDELLRE IDRLFARHAATAYAIGKIDKELSKLELEHAKWLADEIIKLAND PNVSIEEVFKVLEKIIEFEEEFAKDISKAKSIEEVIEIFKEYV EKIKEFLGSGSHHWGSTHHHHHH |
| PDGFR_18 | $< 1.00 \times 10^{-12}$ | MSGMVEKTLEKAKEAKELILKNTAAQLAMRSVLLRLEEDLDDD AIEVLIRLLKMTAALLRYLKLDEKLAELLREIARGLEELKKEV EKKLSDEEIKKKVEELSKEIQKKLKEILEKAKQPGVSLEETVA PEVMDLVIKAIETYAELLSGSGSHHWGSTHHHHHH |
| PDGFR_19 | $6.17 \times 10^{-11}$ | MSGSEIKELKLLLTKLVAAIRYASRHPEDFLDILRLAKEIFER VLEGIKKHPELIPFLSSALQQILAIIKKWMEDPRNQSAELLEA VAEFLEEIIKKLEEVLEYLKSSITPENEEELQKKIEILEATIK RLKELKEEILEKARKLKELEGSGSHHWGSTHHHHHH |
| PDGFR_20 | $1.97 \times 10^{-7}$ | MSGMKKEIEETKKYIEEVIKKVEDDPVLVQILERLLKVLELYE EGKIEEALKELSRVGLMLVALFEANYEDIVEAVYKLLKKKGID SKELAEKEAEKFGKEIKKTALVSYPDFIAKQVKEMVEFLEELY NLLKLKKGSGSHHWGSTHHHHHH |
| PDGFR_27 | $5.96 \times 10^{-8}$ | MSGMLSTLGKIAIVRAYKLIKKGLEEGKIEESLERAAALLFLA SKFEGNNKILLVLTTLIISIASPEDIRGAEAVLEKAKSEGLSE ELYEAVKEFIELMSKIKDEEKHKLAVERLAELLELAGKGELTE EKLTEHIKEMIEELKGSGSHHWGSTHHHHHH |
| PDGFR_29 | $1.24 \times 10^{-5}$ | MSGMDKELAKELLKLLEELEEKLRKEYRNLAGGLGKAMLIFLD IAPWLREKGYEKVAKRTLDILADITEYLRKAGVRTEKEFLEVL KEKVDELIEEAKEALKKGDFEYVFAVILALSILSVGSSKEVQE YLEEKKNELVELLKKEIGSGSHHWGSTHHHHHH |
| PDGFR_30 | $1.24 \times 10^{-9}$ | MSGRQVKVAVALKVKDKALRKMFEKLLEVINKLEDLEKANKGA LYLGATQHIIANETNYELTEENVEDVGEIIAKMIAALMDVSGI PEDDLVEAIKIMYEELVKEGREEMAKKLLETIAKELEKRGYPE LAKKVREILGSGSHHWGSTHHHHHH |

Continued on next page...

Table S7 (Continued)

| Design | $K_D$ (M) | Sequence |
|---|---|---|
| PDL1_07 | $5.26 \times 10^{-7}$ | MSGMDEEDLLRLIEKAKKLIEKLENEEIEDLKEKLRMLSLAAM YILLALKQPSAEKYLEELRELLERVKKLVEELKEKEGVEDKAT ELLLEDAERLLKEGSGSHHWGSTHHHHHH |
| PDL1_10 | $2.02 \times 10^{-7}$ | MSGMWEELEKTLSEDEEVKEEALKKVREYVEKILEEAKKKGKL TPQQEFELLKVVAMLTYVDPEFVEEIIEKVEEIDDERAKKLVE KMKRHLEVAKKRPLRGSGSHHWGSTHHHHHH |
| PDL1_11 | $4.17 \times 10^{-8}$ | MSGSEKEEKTLYLLKVAALFVRQYSKEAAELIEEALELFKQGK YEEALEKVEEARKEIEALGPEHKGLADLLDTVAKEIRELIGSG SHHWGSTHHHHHH |
| PDL1_13 | $5.57 \times 10^{-8}$ | MSGSMMELMEELFKEIGVLFEVMKVRPELEEELRKVMELIRQA LRLLADGEDLAKAEEKLREAWELLEKHKEELLKAIEEVIEKNK DKPKLVKELEEFKKKLEGSGSHHWGSTHHHHHH |
| PDL1_15 | $1.09 \times 10^{-6}$ | MSGMVKELIEAFEKGDEETVLKLLKEAVKLLKEGKLSKSDLIK ITIIATNFLDEPVTDEFIEELEKLDDEWAQRLAETLRRNKEGS GSHHWGSTHHHHHH |
| PDL1_17 | $5.03 \times 10^{-7}$ | MSGSEEELERLRDKILLLLRALILALKDFPELAELLEELREAI EEGDLERTEELLEEIEELVEERREELSKAVLDALESLLRDVRR YIELEKKGSGSHHWGSTHHHHHH |
| PDL1_18 | $9.20 \times 10^{-7}$ | MSGKEKEIEELKDKILLLLRALALALRDDPELVELIEKIAELL EKGDLEKAKELFEEFLELVEKRREELSKAVLDAIESLGKLFEE LLELKKEGSGSHHWGSTHHHHHH |
| PDL1_24 | $5.19 \times 10^{-8}$ | MSGSMSLEEKFKKIMEKMKEAGFIPDKSFSSLEEVAESEDITV LTKVTMFLTNDEEVAKKFLEILEKQYKKLEEEVKKNPEDEELK RKAEHLKKLIEHLRKNLGSGSHHWGSTHHHHHH |
| PDL1_25 | $8.30 \times 10^{-10}$ | MSGSSDLKAAERFLQNVLSFLLPEEAAKKLYEFILELVEKEPD LSREELKERIRELAAELGVELSDKELDSIVDIVELLRKLSGSG SHHWGSTHHHHHH |
| PDL1_26 | $5.14 \times 10^{-12}$ | MSGSSDLEAAKRFLYNVLKFLLPEEVAEKLYEFILELVENEPD LSDEELKKKIRELAKKLGAELTDKQLDSIVDIVNLLRKLSGSG SHHWGSTHHHHHH |
| PDL1_27 | $9.11 \times 10^{-8}$ | MSGDIFEDALAMLRAAALALRMVDEELQEAAEMIEEAVRLAKE GPDDPEALRKALEMVKEAREKVKELRKKSKNKKGLQSLIDMLD KAIELLEEYLKSGSGSHHWGSTHHHHHH |
| TRKA_05 | $7.41 \times 10^{-8}$ | MSGMTEEQLRGLEEIGREVAEVGSLREIMARRPELAGLTIALM LKYGSWEVTVEQVEEAKEEVRRAGEERLARGSGSHHWGSTHHH HHH |
| TRKA_20 | $8.74 \times 10^{-8}$ | MSGSITVSTASSVSVKEGSSLTLTATVTSDFDLSKHPVTVMLE RQFPGEEWELLGFISRRDGETTTAIDGRFQKAFNGGVSLSLTQ VSPHTLQVTLAIGNVKKEKDTAVYRFSVGVLDDEGWSRHNATG ADTKVTVTDGSGSHHWGSTHHHHHH |

Continued on next page...

Table S7 (Continued)

| **Design** | $K_D$ **(M)** | **Sequence** |
|---|---|---|
| VEGFA_01 | $1.11 \times 10^{-9}$ | MSGMKFKITIKLGNVPASEETKATLERILKKVAEEFGVKASISFDFEKGTGEFVFEIEGPLEEAQEAILTLLRAVGMFLADISLGLSFELTIESIEFETDDSGFESPVDLSSLRGKPVLEAARFLFDKVEELEKGSGSHHWGSTHHHHHH |
| VEGFA_03 | $3.33 \times 10^{-10}$ | MSGKKEDEETALFKRLLMVLVKDEKTTTETRALAIAMDIVLDSNKDLKVTKIEAEGLSLEEVEELIRNDIPVIGKFTVTIKKGDEEGKITINIKEEDGKIVVEIEGDPNDKLSSEIVEEIKKLAEMILEAEKAGSGSHHWGSTHHHHHH |
| VEGFA_04 | $< 1.00 \times 10^{-12}$ | MSGSEKERIRALFHRLLMLLVKDERTTDEARALAIAMEIYFTSTKDLEIVDINVNGMSQEEVEEKISKGEVILGKIEITFELDGKKHTIYIEIYEEDGKIKAEISADPNDKVSQELVKKIKELAEMILAAEKAGSGSHHWGSTHHHHHH |
| VEGFA_07 | $3.00 \times 10^{-9}$ | MSGSMAELIELSEEVKMVVYTYELPETNRLVLHILLEVPEELVEEIIEKIKELLKKKGLPELEEVEAGEVMAKGREGKNVYAVVPGEGKAIIEIMLVFEGPREESLKKAEEVLEEIIELLKKILGSGSHHWGSTHHHHHH |
| VEGFA_11 | $3.14 \times 10^{-9}$ | MSGMEEAVRLAVAALRELAKVLSRHLSPEARREIWEIVKRTAEEMARLAGIEIDGRDIARAEAMILAALDGSKDIPEELYEPAKRLGEAVAAVREALDDPAADVAAAIEELTEVMEEVLREGGAPEEAIEVLREGVERIIEIVRREGLAGSGSHHWGSTHHHHHH |
| VEGFA_12 | $1.21 \times 10^{-8}$ | MSGMFDELLKAYEGVKPVDLTEEQTPTLQEVVEELEARGIERPEGFSFDSARGVRQAFLHLGHRALQNPEEHLEEVLTIIEVLLERIKELVDLLKDPENSHLDKEVLKHTLAVLLDILMELVFLNFEHLPKDFQEKLLKVVKEASEVLKGSGSHHWGSTHHHHHH |
| VEGFA_24 | $1.12 \times 10^{-8}$ | MSGMHAPDPSMSVKDVLLLLKHFLEEVLDLLYALTQLEDPEKEADKLVELVRKLGDAYGIHPEIVELIQEVIEKHKDDAKKALEEIEKFLKLVEGLLSLAPILEEKGVSVDELVKLLVELLKKIEEEGIDPEELIKELKEELEGSGSHHWGSTHHHHHH |
| VEGFA_25 | $7.88 \times 10^{-9}$ | MSGMGQSALRVTMNIEVGDIKAGSNLDIIKLPDGTSLLIFEGPGLFDIRELSEEELEQPLRSIVVPLFQKSAENFAKIVIERLGSEDVEAKISVSAQVFDENGKPKIPELSASGTTKVKDLAKEVAELTTKLVDELEKKFGKTLKEILGSGSHHWGSTHHHHHH |
| VEGFA_26 | $1.85 \times 10^{-9}$ | MSGMNLDVASLLVQLSGFDIEKVKEVAEKFVEYLKKNVPDMVIRINVYSSEDSLWIEIHGALPLTEALYKMFKIIAENKDKPLEEIKKKATEEATEIIRRLLEEAGEPELGEKVVSHVEEITELLVELLEKGVESIDSVLNRPGRRPGSGSHHWGSTHHHHHH |
| VEGFA_28 | $1.78 \times 10^{-8}$ | MSGSFSFEYTLEADSEEEREALAVALIGSLYEMKLENPDIISSLTVRLLDSEEEALEESESAIPLVEKILKEKKVTKGSVIHVHLVFLPEKDGKYLLKVHVDVILATGMSKEEQEEVKKKIEELFEEVIKRAKEKVESFKDVLGSGSHHWGSTHHHHHH |
| VEGFA_30 | $2.24 \times 10^{-8}$ | MSGMEALRKLKALLINSSVGADDSTIIAALLLIEIVDASELRLKSKDPNLPEEEKKKLLEDVKWRIENLAKFVKELLEKYAPDVKLEDVVKKVEELAEKVAEKIKELKVKAEELVENLEEVAEELKKLGLDELAKALEEAAEKLKEVSKLGSGSHHWGSTHHHHHH |

Sequences listed include vector-derived residues resulting from subcloning into the pIVEX vector. The N-terminus contains an MSG prefix, and the C-terminus includes the extension GSGSHHWGSTHHHHHH, comprising a SNAC cleavage tag and a 6×His affinity tag.

**Table S8: Sequences and binding affinities of all VHHs per target.**

| Design | $K_D$ (M) | Sequence |
|---|---|---|
| CLL2_01 | $1.46 \times 10^{-6}$ | EVQLVESGGGLVQPGGSLRLSCAASGGTFSSYYMGWFRQAPGKGREL VAAISSSGIYTYYPDSVEGRFTISRDNAKRMVYLQMNSLRAEDTAVY YCAAAEKAIDVFFSASGAKWWGQGTQVTVSSGGGGSHHHHHH |
| CLL2_02 | $5.04 \times 10^{-7}$ | QVQLQESGGGLVQPGGSLRLSCAASGYTFSSYFMAWFRQAPGKERER VAKIYPGSGSTYLADSVKGRFTISQNNAKSTVYLQMNSLKPEDTAMY YCARIDRTKSYDYWGQGTQVTVSGGGGSHHHHHH |
| CLL2_04 | $1.59 \times 10^{-6}$ | QVQLQESGGGLVQPGGSLRLSCAASGFTTSYYAMAWFRQAPGKERER VAKISSSGRYTYLADSVKGRFTISQNNAKSTVYLQMNSLKPEDTAMY YCAASDISTSGFTTSSFSYWGQGTQVTVSGGGGSHHHHHH |
| CLL2_07 | $6.57 \times 10^{-7}$ | QVQLQESGGGLVQPGGSLRLSCAASGFTTSYYFMAWFRQAPGKERER VAKINPATGSTYLADSVKGRFTISQNNAKSTVYLQMNSLKPEDTAMY YCARQNSLTTYDSWGQGTQVTVSGGGGSHHHHHH |
| CLL2_08 | $1.77 \times 10^{-6}$ | EVQLVESGGGLVQPGNSLRLSCAASGFTFSSYRMSWVRQAPGKGLEW VSSISTSGGSKLYADSVKGRFTISRDNAKTTLYLQMNSLRPEDTAVY YCSRRLYTYGSLSSSDVSSQGTLVTVSSGGGGSHHHHHH |
| CLL2_12 | $2.06 \times 10^{-7}$ | EVQLVESGGGLVQPGNSLRLSCAASGFTFSTYRMSWVRQAPGKGLEW VSSISSGGTYKLYADSVKGRFTISRDNAKTTLYLQMNSLRPEDTAVY YCSAVLSSSSNPTSLVSSQGTLVTVSSGGGGSHHHHHH |
| CLL2_14 | $1.91 \times 10^{-7}$ | EVQLVESGGGLVQPGGSLRLSCAASGFTFSSYAMGWFRQAPGKGREL VAAISSGSRNTYYPDSVEGRFTISRDNAKRMVYLQMNSLRAEDTAVY YCARLKTYYSIPFSLTTSAFDSWGQGTQVTVSSGGGGSHHHHHH |
| CLL2_15 | $5.80 \times 10^{-8}$ | QVQLQESGGGLVQPGGSLRLSCAASGSFSIVDRMAWFRQAPGKERER VAKISLYSGNTYLADSVKGRFTISQNNAKSTVYLQMNSLKPEDTAMY YCARTLGYDGSDADYWGQGTQVTVSGGGGSHHHHHH |
| CLL2_18 | $3.04 \times 10^{-7}$ | QVQLQESGGGLVQPGGSLRLSCAASGTLSSVDRMAWFRQAPGKERER VAKMNLYNGNTYLADSVKGRFTISQNNAKSTVYLQMNSLKPEDTAMY YCVRSGGSGSSYWGQGTQVTVSGGGGSHHHHHH |
| CLL2_21 | $2.50 \times 10^{-10}$ | EVQLVESGGGLVQPGNSLRLSCAASGFTFSTYAMSWVRQAPGKGLEW VSSISSSGSYTLYADSVKGRFTISRDNAKTTLYLQMNSLRPEDTAVY YCSRLLYSFSLVKSTLKTSYFGSSSQGTLVTVSSGGGGSHHHHHH |
| CLL2_22 | $4.32 \times 10^{-7}$ | QVQLQESGGGLVQPGGSLRLSCAASGFTTSSAFMAWFRQAPGKERER VAYISSGGGNIYLADSVKGRFTISQNNAKSTVYLQMNSLKPEDTAMY YCASYYNDSSLNRGKRVWGQGTQVTVSGGGGSHHHHHH |
| CLL2_24 | $2.82 \times 10^{-6}$ | EVQLVESGGGLVQPGNSLRLSCAASGFRFSSYAMSWVRQAPGKGLEW VSSISWDGSNKLYADSVKGRFTISRDNAKTTLYLQMNSLRPEDTAVY YCRRTGSYDSTSLSSGLGSSQGTLVTVSSGGGGSHHHHHH |
| CLL2_27 | $8.41 \times 10^{-7}$ | EVQLVESGGGLVQPGGSLRLSCAASGGSFSYYRMGWFRQAPGKGREL VAAISYSGGSTYYPDSVEGRFTISRDNAKRMVYLQMNSLRAEDTAVY YCARLLTTRADSAPDYWGQGTQVTVSSGGGGSHHHHHH |

Continued on next page...

Table S8 (Continued)

| Design | $K_D$ (M) | Sequence |
|---|---|---|
| 1433E_04 | $5.71 \times 10^{-8}$ | EVQLVESGGGLVQPGGSLRLSCAASGDYGTDYNMAWYRQAPGKGREL VAGISYDGRYKSYADSVKGRFTISRDNAKNTLYLQMNSLRPEDTAVY YCRAARRDAYYVRSDSYVAWGQGTLVTVSSGGGGSHHHHHH |
| 1433E_10 | $1.13 \times 10^{-7}$ | EVQLVESGGGLVQPGGSLRLSCAASGALLDMSTMAWYRQAPGKGREL VAGINLYSGTTSYADSVKGRFTISRDNAKNTLYLQMNSLRPEDTAVY YCVSRTEFSDTAYARRHQAYFWGQGTLVTVSSGGGGSHHHHHH |
| 1433E_11 | $2.85 \times 10^{-8}$ | QVQLQESGGGLVQPGGSLRLSCAASGATSSVDYMAWFRQAPGKERER VAKMNVWYGNTYLADSVKGRFTISQNNAKSTVYLQMNSLKPEDTAMY YCAAVTTLFSGRAYAYWGQGTQVTVSGGGGSHHHHHH |
| 1433E_12 | $6.47 \times 10^{-9}$ | EVQLVESGGGLVQPGGSLRLSCAASGGTLSSYYMGWFRQAPGKGREL VAAISRSGIYTYYPDSVEGRFTISRDNAKRMVYLQMNSLRAEDTAVY YCAAVPYDDIDYWRRAGDYLYWGQGTQVTVSSGGGGSHHHHHH |
| 1433E_24 | $5.11 \times 10^{-8}$ | EVQLVESGGGLVQPGGSLRLSCAASGFTFSRAAMGWFRQAPGKGREL VAAISSDGSYHYYPDSVEGRFTISRDNAKRMVYLQMNSLRAEDTAVY YCATVYENYYSGMMYEAWGQGTQVTVSSGGGGSHHHHHH |
| 1433E_27 | $1.66 \times 10^{-8}$ | EVQLVESGGGLVQPGGSLRLSCAASGSYRGFDFMAWYRQAPGKGREL VAGINDGSGYTSYADSVKGRFTISRDNAKNTLYLQMNSLRPEDTAVY YCRAATIYFSGRATVTWGQGTLVTVSSGGGGSHHHHHH |
| PDGFR_01 | $4.44 \times 10^{-8}$ | EVQLVESGGGLVQPGGSLRLSCAASGGTFIETFMGWFRQAPGKGREL VAAISSSGSRTYYPDSVEGRFTISRDNAKRMVYLQMNSLRAEDTAVY YCAAYYLDRSSKTLKVSWGQGTQVTVSSGGGGSHHHHHH |
| PDGFR_03 | $1.69 \times 10^{-7}$ | EVQLVESGGGLVQPGGSLRLSCAASGSSSSSYFMAWFRQAPGQEREF VAGINWSGNRTLYADSVRGRFTNSRDNSKNTLYLQMNSLRAEDTAVY YCTTLDMRNRSVSYWGQGTLVTVSSGGGGSHHHHHH |
| PDGFR_04 | $3.61 \times 10^{-7}$ | EVQLVESGGGLVQPGGSLRLSCAASGASLHSGFMGWFRQAPGKGREL VAAINNRNGDTYYPDSVEGRFTISRDNAKRMVYLQMNSLRAEDTAVY YCAVMGYYTSYSSKTTNKAYWGQGTQVTVSSGGGGSHHHHHH |
| PDGFR_05 | $2.97 \times 10^{-8}$ | EVQLVESGGGLVQPGGSLRLSCAASYGNQIITMMAWFRQAPGQEREF VAGIYVASGYTLYADSVRGRFTNSRDNSKNTLYLQMNSLRAEDTAVY YCSSWYYSSSGSLTNTWGQGTLVTVSSGGGGSHHHHHH |
| PDGFR_06 | $1.12 \times 10^{-7}$ | EVQLVESGGGLVQPGGSLRLSCAASGSTSSYTAMAWFRQAPGQEREF VAGIYAGSGTTLYADSVRGRFTNSRDNSKNTLYLQMNSLRAEDTAVY YCASYLISGTLTVTWGQGTLVTVSSGGGGSHHHHHH |
| PDGFR_07 | $1.56 \times 10^{-8}$ | EVQLVESGGGLVQPGGSLRLSCAASGGTFTSYAMAWFRQAPGQEREF VAGINSSGGFTLYADSVRGRFTNSRDNSKNTLYLQMNSLRAEDTAVY YCARHSHSWITKYTSTFSPSYMYWGQGTLVTVSSGGGGSHHHHHH |
| PDGFR_08 | $4.11 \times 10^{-8}$ | EVQLVESGGGLVQPGGSLRLSCAASGTYAFVALMGWFRQAPGKGREL VAAISISSGYTYYPDSVEGRFTISRDNAKRMVYLQMNSLRAEDTAVY YCAAMTLSSGKSVISWGQGTQVTVSSGGGGSHHHHHH |
| PDGFR_09 | $9.24 \times 10^{-8}$ | EVQLVESGGGLVQPGGSLRLSCAASGFSGYAAMMAWFRQAPGQEREF VAGISWNGSYTLYADSVRGRFTNSRDNSKNTLYLQMNSLRAEDTAVY YCAQLDMKSGKSTVYWGQGTLVTVSSGGGGSHHHHHH |

Continued on next page...

Table S8 (Continued)

| Design | $K_D$ (M) | Sequence |
|---|---|---|
| PDGFR_11 | $8.09 \times 10^{-9}$ | EVQLVESGGGLVQPGGSLRLSCAASGGTFSNYFMAWFRQAPGQEREF VAGISWSGSRTLYADSVRGRFTNSRDNSKNTLYLQMNSLRAEDTAVY YCASIGIYSGTVKTYWGQGTLVTVSSGGGGSHHHHHH |
| PDGFR_12 | $3.10 \times 10^{-8}$ | EVQLVESGGGLVQPGGSLRLSCAASGGSFSTNFMAWYRQAPGKGREL VAGISFSGSRTSYADSVKGRFTISRDNAKNTLYLQMNSLRPEDTAVY YCSAFEIKRGKLTVYWGQGTLVTVSSGGGGSHHHHHH |
| PDGFR_13 | $1.56 \times 10^{-7}$ | EVQLVESGGGLVQPGGSLRLSCAASGGTFYQYFMAWFRQAPGQEREF VAGISVNGSYTLYADSVRGRFTNSRDNSKNTLYLQMNSLRAEDTAVY YCATLSVRGGSSTWGQGTLVTVSSGGGGSHHHHHH |
| PDGFR_14 | $3.96 \times 10^{-8}$ | EVQLVESGGGLVQPGGSLRLSCAASGDRGGVYPMAWYRQAPGKGREL VAGISSGGSQTSYADSVKGRFTISRDNAKNTLYLQMNSLRPEDTAVY YCARGYWTAATAWVAGMFAYYGYWGQGTLVTVSSGGGGSHHHHHH |
| PDGFR_15 | $1.57 \times 10^{-8}$ | EVQLVESGGGLVQPGGSLRLSCAASGSLTGGYWMAWFRQAPGQEREF VAGLLYHGTRTLYADSVRGRFTNSRDNSKNTLYLQMNSLRAEDTAVY YCAVMDLRNSSTSYWGQGTLVTVSSGGGGSHHHHHH |
| PDGFR_16 | $4.09 \times 10^{-9}$ | EVQLVESGGGLVQPGGSLRLSCAASGTTQSIVMMAWYRQAPGKGREL VAGISNTGRYISYADSVKGRFTISRDNAKNTLYLQMNSLRPEDTAVY YCAVWQISYSGSTTWTVYWGQGTLVTVSSGGGGSHHHHHH |
| PDGFR_17 | $1.89 \times 10^{-6}$ | EVQLVESGGGLVQPGGSLRLSCAASGGTFTSYFMAWYRQAPGKGREL VAGTSWSGIRKSYADSVKGRFTISRDNAKNTLYLQMNSLRPEDTAVY YCTALEIRGTSDGTARADYTYWGQGTLVTVSSGGGGSHHHHHH |
| PDGFR_18 | $2.42 \times 10^{-7}$ | EVQLVESGGGLVQPGGSLRLSCAASGTISSVVIMAWFRQAPGQEREF VAGIYVWSGSTLYADSVRGRFTNSRDNSKNTLYLQMNSLRAEDTAVY YCSTFAISSGTLTVTWGQGTLVTVSSGGGGSHHHHHH |
| PDGFR_20 | $6.37 \times 10^{-9}$ | EVQLVESGGGLVQPGGSLRLSCAASGTTGNIALMAWYRQAPGKGREL VAGISRSGSYTSYADSVKGRFTISRDNAKNTLYLQMNSLRPEDTAVY YCAAFAASYSGSSSWTVVWGQGTLVTVSSGGGGSHHHHHH |
| PDGFR_21 | $1.82 \times 10^{-8}$ | EVQLVESGGGLVQPGGSLRLSCAASGGTFSSYFMAWFRQAPGQEREF VAGISPGGGRTLYADSVRGRFTNSRDNSKNTLYLQMNSLRAEDTAVY YCATIPFSSSYRLSKDTSPYWGQGTLVTVSSGGGGSHHHHHH |
| PDGFR_22 | $8.26 \times 10^{-8}$ | EVQLVESGGGLVQPGGSLRLSCAASGGTFTNHFMAWFRQAPGQEREF VAGISPSGSTTLYADSVRGRFTNSRDNSKNTLYLQMNSLRAEDTAVY YCAAIDLRTGSTVWGQGTLVTVSSGGGGSHHHHHH |
| PDGFR_23 | $9.66 \times 10^{-8}$ | EVQLVESGGGLVQPGGSLRLSCAASGGNFHNYFMAWFRQAPGQEREF VAGISSSGSYTLYADSVRGRFTNSRDNSKNTLYLQMNSLRAEDTAVY YCAAYDYSGNTVWGQGTLVTVSSGGGGSHHHHHH |
| PDGFR_25 | $5.90 \times 10^{-8}$ | EVQLVESGGGLVQPGGSLRLSCAASGGTFSSYGMAWFRQAPGQEREF VAGISGSGSSTLYADSVRGRFTNSRDNSKNTLYLQMNSLRAEDTAVY YCAKDMFSGLRIRVTASSMLAWGQGTLVTVSSGGGGSHHHHHH |
| PDGFR_26 | $2.32 \times 10^{-7}$ | EVQLVESGGGLVQPGGSLRLSCAASGGTFISHFMGWFRQAPGKGREL VAAISWNGSSTYYPDSVEGRFTISRDNAKRMVYLQMNSLRAEDTAVY YCASWYWSSSGTLKVTWGQGTQVTVSSGGGGSHHHHHH |

Continued on next page...

Table S8 (Continued)

| Design | $K_D$ (M) | Sequence |
|---|---|---|
| PDGFR_27 | $1.53 \times 10^{-7}$ | EVQLVESGGGLVQPGGSLRLSCAASGSSSGASHMAWYRQAPGKGREL<br>VAGIDRGTGDTSYADSVKGRFTISRDNAKNTLYLQMNSLRPEDTAVY<br>YCASTYYAGKTFILGYITSLFRYWGQGTLVTVSSGGGGSHHHHHH |
| PDGFR_29 | $1.52 \times 10^{-7}$ | EVQLVESGGGLVQPGGSLRLSCAASGGSLAGGFMAWFRQAPGQEREF<br>VAGINLASGSTLYADSVRGRFTNSRDNSKNTLYLQMNSLRAEDTAVY<br>YCARLYWDRSTKKWKYVWGQGTLVTVSSGGGGSHHHHHH |
| PDGFR_30 | $8.16 \times 10^{-8}$ | EVQLVESGGGLVQPGGSLRLSCAASGGNLLDHFMGWFRQAPGKGREL<br>VAAISASGSSTYYPDSVEGRFTISRDNAKRMVYLQMNSLRAEDTAVY<br>YCAAISVYGSSAAWGQGTQVTVSSGGGGSHHHHHH |
| BHRF1_04 | $3.64 \times 10^{-7}$ | EVQLVESGGGLVQPGGSLRLSCAASGGTFSSYGMAWYRQAPGKGREL<br>VAGISSSGSSTSYADSVKGRFTISRDNAKNTLYLQMNSLRPEDTAVY<br>YCAAEDEISYYGLYIYATVAWGQGTLVTVSSGGGGSHHHHHH |
| BHRF1_05 | $2.38 \times 10^{-7}$ | EVQLVESGGGLVQPGGSLRLSCAASGGTFSLYAMGWFRQAPGKGREL<br>VAAISGGGSYTYYPDSVEGRFTISRDNAKRMVYLQMNSLRAEDTAVY<br>YCAREFFGTTIAWGQGTQVTVSSGGGGSHHHHHH |
| BHRF1_09 | $2.38 \times 10^{-7}$ | EVQLVESGGGLVQPGGSLRLSCAASGSSTTAMIMAWYRQAPGKGREL<br>VAGILNGLSITSYADSVKGRFTISRDNAKNTLYLQMNSLRPEDTAVY<br>YCARERFVSTSSGTVRWSTYWGQGTLVTVSSGGGGSHHHHHH |
| BHRF1_13 | $5.95 \times 10^{-7}$ | EVQLVESGGGLVQPGGSLRLSCAASGFTFSLWDMGWFRQAPGKGREL<br>VAAISNGGDYTYYPDSVEGRFTISRDNAKRMVYLQMNSLRAEDTAVY<br>YCAAVLGYDWIRSGLYWGQGTQVTVSSGGGGSHHHHHH |
| BHRF1_30 | $1.31 \times 10^{-7}$ | EVQLVESGGGLVQPGGSLRLSCAASGGTFSLWDMGWFRQAPGKGREL<br>VAAISSSGSSTYYPDSVEGRFTISRDNAKRMVYLQMNSLRAEDTAVY<br>YCARGGRYLDYFWGMSWGQGTQVTVSSGGGGSHHHHHH |
| EFNA1_02 | $6.51 \times 10^{-6}$ | EVQLVESGGGLVQPGGSLRLSCAASGSSGMPDIMGWFRQAPGKGREL<br>VAEMHIFTGDTYYPDSVEGRFTISRDNAKRMVYLQMNSLRAEDTAVY<br>YCAGRDSSSGGTVYWGQGTQVTVSSGGGGSHHHHHH |
| EFNA1_03 | $8.76 \times 10^{-6}$ | EVQLVESGGGLVQPGGSLRLSCAASGETSSVSYMGWFRQAPGKGREE<br>VAAINTVSGYTYYPDSVEGRFTISRDNAKRMVYLQMNSLRAEDTAVY<br>YCAARDARDDGISWGQGTQVTVSSGGGGSHHHHHH |
| EFNA1_04 | $3.12 \times 10^{-7}$ | EVQLVESGGGLVQPGGSLRLSCAASGLSTTGYAMAWFRQAPGQEREF<br>VAGISGDGSYTYYADSVRGRFTNSRDESKNTLYLQMNSLRAEDTAVY<br>YCARSTSSYYYGVEDMDYWGQGTLVTVSSGGGGSHHHHHH |
| EFNA1_06 | $7.17 \times 10^{-7}$ | EVQLVESGGGLVQPGGSLRLSCAASGETSVLSVMAWYRQAPGKGREL<br>VAGISYDGSITSYADSVKGRFTISRDNAKNTLYLQMNSLRPEDTAVY<br>YCAGSTSYSTSWELSSFSYWGQGTLVTVSSGGGGSHHHHHH |
| EFNA1_07 | $6.86 \times 10^{-7}$ | EVQLVESGGGLVQPGGSLRLSCAASGGTFSYYSMGWFRQAPGKGREL<br>VAAISWDGSETYYPDSVEGRFTISRDNAKRMVYLQMNSLRAEDTAVY<br>YCAAETSYYPYPGDTSYAYWGQGTQVTVSSGGGGSHHHHHH |
| EFNA1_08 | $8.22 \times 10^{-7}$ | EVQLVESGGGLVQPGGSLRLSCAASGGEISSYSMAWYRQAPGKGREL<br>VAGISADGRYTSYADSVKGRFTISRDNAKNTLYLQMNSLRPEDTAVY<br>YCATLTANYLNYDFTNTGDFDYWGQGTLVTVSSGGGGSHHHHHH |

Continued on next page...

Table S8 (Continued)

| **Design** | $K_D$ **(M)** | **Sequence** |
|---|---|---|
| EFNA1_09 | $7.21 \times 10^{-7}$ | EVQLVESGGGLVQPGGSLRLSCAASGIYDYASYMGWFRQAPGKGREL VAEMSIFTGNTYYPDSVEGRFTISRDNAKRMVYLQMNSLRAEDTAVY YCAALQSGGSYDVYWGQGTQVTVSSGGGGSHHHHHH |
| EFNA1_10 | $1.06 \times 10^{-6}$ | EVQLVESGGGLVQPGGSLRLSCAASGETSVMSVMAWYRQAPGKGREL VAGISYDGSITSYADSVKGRFTISRDNAKNTLYLQMNSLRPEDTAVY YCAASTSYSTSYELSSFDYWGQGTLVTVSSGGGGSHHHHHH |
| EFNA1_12 | $7.08 \times 10^{-7}$ | EVQLVESGGGLVQPGGSLRLSCAASGGTFSTYPMAWFRQAPGQEREF VAGIGAGGDYTLYADSVRGRFTNSRDNSKNTLYLQMNSLRAEDTAVY YCARYKYETDASNAWEFEDFDSWGQGTLVTVSSGGGGSHHHHHH |
| EFNA1_13 | $1.33 \times 10^{-6}$ | EVQLVESGGGLVQPGGSLRLSCAASGGTFSSYAMAWFRQAPGQEREF VAGINYNGNDTLYADSVRGRFTNSRDNSKNTLYLQMNSLRAEDTAVY YCARLSLQWEAYNEEDVDAWGQGTLVTVSSGGGGSHHHHHH |
| EFNA1_14 | $8.55 \times 10^{-7}$ | EVQLVESGGGLVQPGGSLRLSCAASGSSSSYSIMGWFRQAPGKGREL VAEMSTFTGNTYYPDSVEGRFTISRDNAKRMVYLQMNSLRAEDTAVY YCAALDMSGTYDYYWGQGTQVTVSSGGGGSHHHHHH |
| EFNA1_15 | $2.96 \times 10^{-6}$ | EVQLVESGGGLVQPGGSLRLSCAASGGTFSYYSMGWFRQAPGKGREL VAAITYDGSETYYPDSVEGRFTISRDNAKRMVYLQMNSLRAEDTAVY YCASASSYYAYSDDSSYYYWGQGTQVTVSSGGGGSHHHHHH |
| EFNA1_16 | $2.76 \times 10^{-7}$ | EVQLVESGGGLVQPGGSLRLSCAASGGDVSSRTMAWYRQAPGKGREL VAGISADGRYTSYADSVKGRFTISRDNAKNTLYLQMNSLRPEDTAVY YCATADRNYLNYDFTSTSDFSYWGQGTLVTVSSGGGGSHHHHHH |
| EFNA1_17 | $4.33 \times 10^{-7}$ | EVQLVESGGGLVQPGGSLRLSCAASGSTSIITSMAWYRQAPGKGREL VAGISRSGGYTSYADSVKGRFTISRDNAKNTLYLQMNSLRPEDTAVY YCAATTSLIYSYDSEIITDSMQFWGQGTLVTVSSGGGGSHHHHHH |
| EFNA1_18 | $8.09 \times 10^{-7}$ | EVQLVESGGGLVQPGGSLRLSCAASGPADDLSVMAWYRQAPGKGREL VAGIWYDGSNTSYADSVKGRFTISRDNAKNTLYLQMNSLRPEDTAVY YCAGQLSYYTSYETSSFQYWGQGTLVTVSSGGGGSHHHHHH |
| EFNA1_19 | $1.29 \times 10^{-6}$ | EVQLVESGGGLVQPGGSLRLSCAASGFTFSYYAMGWFRQAPGKGREL VAAITYDGSETYYPDSVEGRFTISRDNAKRMVYLQMNSLRAEDTAVY YCAAVSSSYPYLDDTSYAYWGQGTQVTVSSGGGGSHHHHHH |
| EFNA1_20 | $5.66 \times 10^{-7}$ | EVQLVESGGGLVQPGGSLRLSCAASGFTFSSYMMGWFRQAPGKGREL VAAISYDGSSTYYPDSVEGRFTISRDNAKRMVYLQMNSLRAEDTAVY YCARLGVHSSTAEEIAARNVDSWGQGTQVTVSSGGGGSHHHHHH |
| EFNA1_21 | $1.01 \times 10^{-6}$ | EVQLVESGGGLVQPGGSLRLSCAASGDSDYFAIMGWFRQAPGKGREL VAEMHVFTGNTYYPDSVEGRFTISRDNAKRMVYLQMNSLRAEDTAVY YCAALNMAGTYDYYWGQGTQVTVSSGGGGSHHHHHH |
| EFNA1_22 | $9.30 \times 10^{-7}$ | EVQLVESGGGLVQPGGSLRLSCAASGDASSVQTMAWYRQAPGKGFEL VAGMNRDGSSTSYADSVKGRFTISRDNAKNTLYLQMNSLRPEDTAVY YCVASGSTRTSDIALSYADWGQGTLVTVSSGGGGSHHHHHH |
| EFNA1_24 | $2.79 \times 10^{-6}$ | EVQLVESGGGLVQPGGSLRLSCAASGFTFSYYAMGWFRQAPGKGREL VAAITNDGSETYYPDSVEGRFTISRDNAKRMVYLQMNSLRAEDTAVY YCAATSSSYPYLDDTSYRYWGQGTQVTVSSGGGGSHHHHHH |

Continued on next page...

Table S8 (Continued)

| Design | $K_D$ (M) | Sequence |
|---|---|---|
| EFNA1_25 | $4.47 \times 10^{-6}$ | EVQLVESGGGLVQPGGSLRLSCAASGDYDSVSIMGWFRQAPGKGREE VAAINMVSGHTYYPDSVEGRFTISRDNAKRMVYLQMNSLRAEDTAVY YCAARDASNGGVVWGQGTQVTVSSGGGGSHHHHHH |
| S100A4_01 | $3.67 \times 10^{-7}$ | QVQLQESGGGLVQPGGSLRLSCAASGYTTSYYWMAWFRQAPGKERER VAKIIPILGYTYLADSVKGRFTISQNNAKSTVYLQMNSLKPEDTAMY YCARSKSWTYDSTSFDYWGQGTQVTVSGGGGSHHHHHH |
| S100A4_03 | $2.38 \times 10^{-7}$ | EVQLVESGGGLVQPGGSLRLSCAASGQDAYIWAMAWYRQAPGKGREL VAGIYFGGGTTSYADSVKGRFTISRDNAKNTLYLQMNSLRPEDTAVY YCATYTYSGSLSRTWGQGTLVTVSSGGGGSHHHHHH |
| S100A4_07 | $3.71 \times 10^{-7}$ | EVQLVESGGGLVQPGGSLRLSCAASGGPDYFFYMAWYRQAPGKGREL VAGINFMGGSISYADSVKGRFTISRDNAKNTLYLQMNSLRPEDTAVY YCRFAQFASLSSNGGSVLTYSWGQGTLVTVSSGGGGSHHHHHH |
| S100A4_17 | $7.17 \times 10^{-7}$ | QVQLQESGGGLVQPGGSLRLSCAASGSSNAWIAMAWFRQAPGKERER VAKIYFFGTNKYLADSVKGRFTISQNNAKSTVYLQMNSLKPEDTAMY YCVSYSPSTSSSGGYTRAWGQGTQVTVSGGGGSHHHHHH |
| S100A4_20 | $5.41 \times 10^{-7}$ | EVQLVESGGGLVQPGGSLRLSCAASGDTSFIIAMAWYRQAPGKGREL VAGLNRLTSSISYADSVKGRFTISRDNAKNTLYLQMNSLRPEDTAVY YCAAARVLGGTTERAWGQGTLVTVSSGGGGSHHHHHH |
| HNMT_03 | $7.68 \times 10^{-8}$ | EVQLVESGGGLVQPGNSLRLSCAASGFSFSDYWMSWVRQAPGKGLEW VSSISWNGSRKLYADSVKGRFTISRDNAKTTLYLQMNSLRPEDTAVY YCYRLGRFVSSLNVEKDTFLGSSQGTLVTVSSGGGGSHHHHHH |
| HNMT_10 | $2.66 \times 10^{-9}$ | EVQLVESGGGLVQPGNSLRLSCAASGFSFSRYAMSWVRQAPGKGLEW VSSISGSGSYKLYADSVKGRFTISRDNAKTTLYLQMNSLRPEDTAVY YCTRLIDSDADMDIVRSSYVSSQGTLVTVSSGGGGSHHHHHH |
| HNMT_12 | $2.51 \times 10^{-9}$ | EVQLVESGGGLVQPGNSLRLSCAASGFSFSRYGMSWVRQAPGKGLEW VSSISSSGGTKLYADSVKGRFTISRDNAKTTLYLQMNSLRPEDTAVY YCSRYWYFDELLFTYYSSQGTLVTVSSGGGGSHHHHHH |
| HNMT_18 | $4.37 \times 10^{-8}$ | EVQLVESGGGLVQPGGSLRLSCAASGYTGSDYYMAWYRQAPGKGREL VAGISWTGITKSYADSVKGRFTISRDNAKNTLYLQMNSLRPEDTAVY YCTSSRFRSYDGRAEPDAYWGQGTLVTVSSGGGGSHHHHHH |
| HNMT_23 | $2.50 \times 10^{-9}$ | EVQLVESGGGLVQPGGSLRLSCAASGGDVSYWSMAWYRQAPGKGREL VAGISGSGRFTSYADSVKGRFTISRDNAKNTLYLQMNSLRPEDTAVY YCAVGGVGDTGLFWGQGTLVTVSSGGGGSHHHHHH |
| HNMT_25 | $1.15 \times 10^{-7}$ | EVQLVESGGGLVQPGGSLRLSCAASGFAFSLYSMGWFRQAPGKGREL VAAIKGWSGNTYYPDSVEGRFTISRDNAKRMVYLQMNSLRAEDTAVY YCASGRWFGPLTTSDFSSWGQGTQVTVSSGGGGSHHHHHH |

For expression in CHO cells (pCDNA3.4 vector), an N-terminal signal peptide (MGWSCIILFLVATATGVHS) was fused to the listed sequences but is omitted from the table. The C-terminus includes the shown extension GGGGSHHHHHH (flexible linker and 6×His tag).

### References

[1] Ian Sillitoe et al. “CATH: increased structural coverage of functional space”. In: *Nucleic acids research* 49.D1 (2021), pp. D266–D273.

[2] Helen M Berman et al. “The protein data bank”. In: *Nucleic acids research* 28.1 (2000), pp. 235–242.

[3] Martin Steinegger and Johannes Söding. “MMseqs2 enables sensitive protein sequence searching for the analysis of massive data sets”. In: *Nature biotechnology* 35.11 (2017), pp. 1026–1028.

[4] James Dunbar et al. “SAbDab: the structural antibody database”. In: *Nucleic acids research* 42.D1 (2014), pp. D1140–D1146.

[5] John Jumper et al. “Highly accurate protein structure prediction with AlphaFold”. In: *Nature* 596.7873 (2021), pp. 583–589.

[6] Jason Yim et al. “Fast protein backbone generation with SE(3) flow matching”. In: *arXiv preprint arXiv:2310.05297* (2023).
